# The resting and ligand-bound states of the membrane-embedded human T-cell receptor–CD3 complex

**DOI:** 10.1101/2023.08.22.554360

**Authors:** Ryan Q. Notti, Fei Yi, Søren Heissel, Martin W. Bush, Zaki Molvi, Pujita Das, Henrik Molina, Christopher A. Klebanoff, Thomas Walz

## Abstract

The T-cell receptor (TCR) initiates T-lymphocyte activation, but mechanistic questions remain(*1–4*). Here, we present cryogenic electron microscopy structures for the unliganded and human leukocyte antigen (HLA)-bound human TCR–CD3 complex in nanodiscs that provide a native-like lipid environment. Distinct from the “open and extended” conformation seen in detergent(*5–8*), the unliganded TCR–CD3 in nanodiscs adopts two related “closed and compacted” conformations that represent its physiologic resting state *in vivo*. By contrast, the HLA-bound complex adopts the open and extended conformation, and conformation-locking disulfide mutants show that ectodomain opening is necessary for maximal ligand-dependent T-cell activation. Together, these results reveal allosteric conformational change during TCR activation and highlight the importance of native-like lipid environments for membrane protein structure determination.

## Main Text

T cells are central to a wide array of both adaptive and pathologic immune responses. The T-cell receptor (TCR), in association with members of the CD3 protein family, allows T cells to recognize and respond to antigenic peptides presented on human leukocyte antigen (HLA) complexes of other cells. While the identities of the TCR–CD3 complex constituents and its downstream signaling elements have been known for some time, questions remain regarding its activation mechanism.

The TCR–CD3 is a hetero-octameric membrane protein complex(*5–8*). The αβ-type TCR is comprised in part by two variable chains, TCRα and TCRβ, which directly interface with the HLA–antigen complex. Antigen specificity is generated through somatic recombination of the variable segments in the TCRα and TCRβ genes. The remainder of the core complex is comprised of six invariant CD3 family chains: the CD3γ-CD3 and CD3δ-CD3 heterodimers, and a CD3ζ homodimer. Herein, we differentiate the CD3 protomers by the CD3 component with which each pairs (CD3_D_ pairs with CD3δ, and CD3 _G_ pairs with CD3γ) and the CD3ζ protomers as CD3ζ and CD3ζ’.

While the TCRα and TCRβ chains lack intracellular signaling domains, the CD3 proteins possess cytoplasmic immunoreceptor tyrosine activation motifs (ITAMs) that become phosphorylated upon TCR-HLA interaction and then bind and activate downstream signaling kinases. Dynamic interactions between ITAMs and the membrane regulate signaling(*9*, *10*), but the mechanism by which ligand binding to the extracellular TCRα-TCRβ domains results in phosphorylation of cytoplasmic CD3 ITAMs remains a matter of debate. Specifically, it is unclear whether ligand binding and activation involves conformational change in the TCR–CD3 complex. While reconstitution of TCR–CD3 signaling *in vivo* with minimal constructs has argued against transmembrane conformational change(*11*), biochemical evidence in support of conformational change(*12–15*) and specifically for reversible structural extension of the TCR under mechanical force suggests the opposite(*16*), although the structural details of such conformational transitions remain to be determined.

Structures of the hetero-octameric αβ-type TCR–CD3 have been determined by cryogenic electron microscopy (cryo-EM)(*5–8*). These studies revealed an antenna-like structure, with the TCRα and TCRβ variable domains projecting over a ring formed by the CD3 ectodomains and the TCRα and TCRβ constant domains, with numerous interactions between their respective transmembrane (TM) helices (Fig. S1). Addition of HLA–antigen to this complex did not meaningfully alter the conformation of the TCR–CD3(*7*, *8*), arguing against models of intrinsic conformational change in the ligand-dependent TCR–CD3 activation process (Fig. S1).

However, the TCR–CD3 complexes used in these studies were solubilized in detergent. Biochemical data have previously shown important roles for the lipid bilayer in the regulation of TCR–CD3 signaling specifically(*6*, *17*, *18*), raising questions as to the physiologic meaning of the determined structures. Membrane protein structure determination is generally preferable in a native-like membrane environment that increases the likelihood that important protein–lipid interactions are captured in the obtained molecular models(*19*).

Here, we report cryo-EM structures of a human TCR–CD3 stabilized both in detergent micelles and nanodiscs. The structures reveal novel conformations for the TCR–CD3 in a native-like lipid environment, distinct from those previously reported, and cross-linking data support the notion that these conformations represent the resting state of an unstimulated TCR–CD3 on T cells. We then determined the structure of the same nanodisc-embedded TCR–CD3 complex bound to HLA, revealing substantial conformational change upon physiologic ligand binding, and show that exit from the resting state conformation is required for maximal TCR–CD3 signaling.

## Results

### Structure of the NY-ESO-1-specific 1G4 TCR–CD3 in detergent micelles

In this study, we investigated the structure of the 1G4 (LY-variant) TCR–CD3. We chose the 1G4 TCR, which binds to an HLA-A*02:01-restricted 9 amino acid epitope derived from the NY-ESO-1 cancer-testis antigen, because it is both a model immunoreceptor and being developed as an anti-cancer therapeutic(*20*). To confirm that 1G4 has a similar structure to the other TCR– CD3 complexes analyzed in detergent, we purified this complex in glyco-diosgenin (GDN) and determined its structure by cryo-EM to a nominal resolution of 3.3 Å (Fig. 1A, S1-3; Tables S1-3). Herein, this model and the conformation it represents will be abbreviated as “GDN.” Overall, GDN was very similar to the structures of three other TCR–CD3 constructs, including those with and without HLA bound (Fig. S3E). This is not surprising, given the nearly identical CD3 and TCR constant region sequences shared by these receptors (Fig. S3F).

**Fig. 1:**
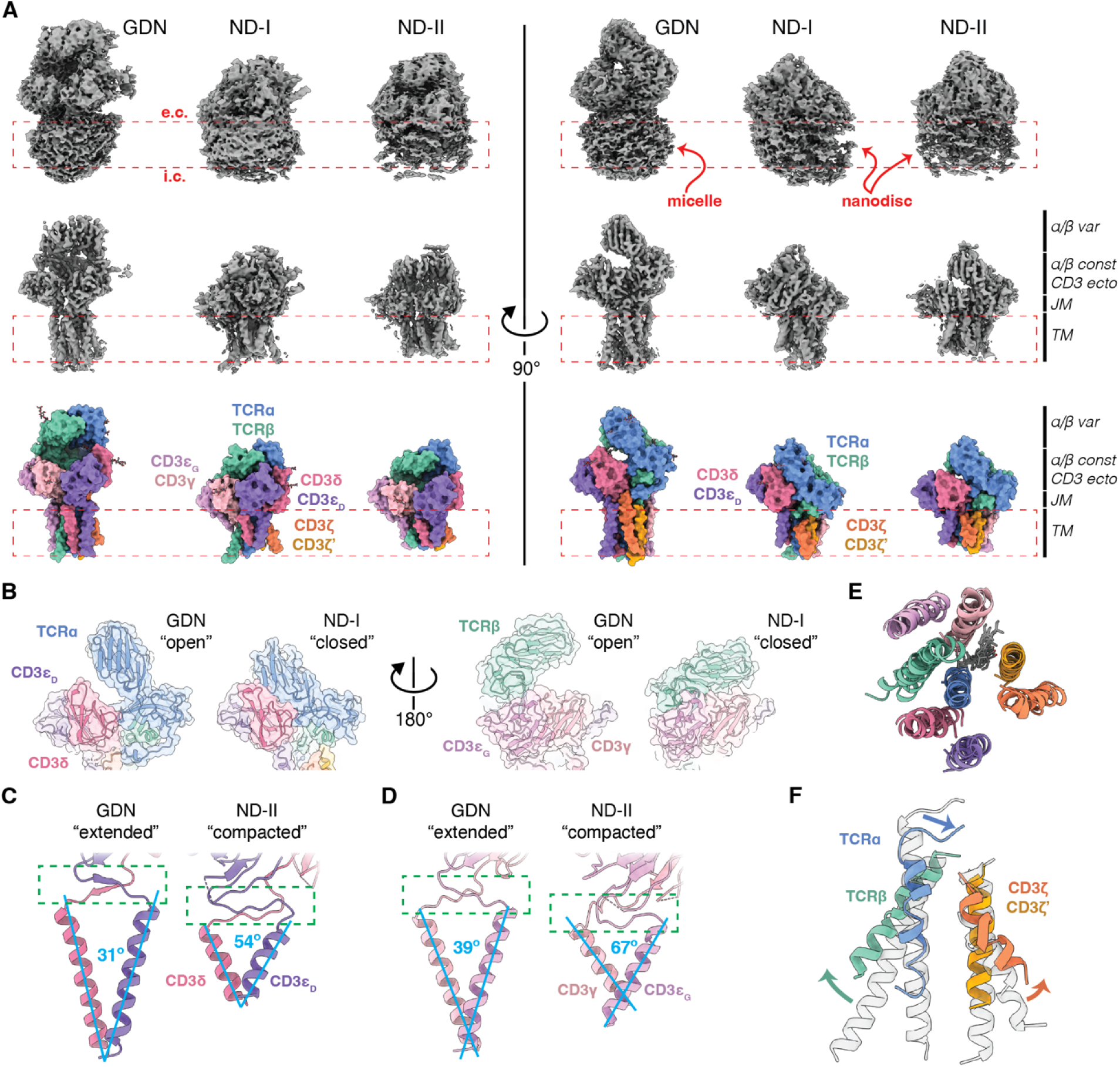
Architecture of the TCR–CD3 in a native-like lipid environment. (**A**) Cryo-EM maps (top, middle) and space-filling solvent-excluded surfaces for the models (bottom; herein, model surfaces) of the TCR–CD3 in a detergent micelle (GDN) and in nanodiscs (conformations ND-I and ND-II). Top row shows cryo-EM density (without local refinement) after B-factor sharpening, contoured to a low threshold to highlight micelle and nanodisc positioning. Red dashed box indicates approximate plasma membrane span as inferred from the micelle and nanodisc densities (red arrows and labels). Extracellular and intracellular faces indicated as e.c. and i.c., respectively. Middle row shows DeepEMhancer-sharpened maps after local refinement, accentuating the protein and glycan densities. Protein regions indicated at far right abbreviated as follows: var, variable domains; ecto, ectodomains; const, constant domains; JM, juxtamembrane linkers; TM, transmembrane helices. (**B**) Comparison of GDN and ND-I models, highlighting ectodomain closure in the context of a lipid bilayer. Model backbone cartoon inside transparent model surfaces shown. (**C,D**) Comparison of GDN and ND-II models, highlighting juxtamembrane linker compaction in the direction normal to the lipid bilayer (green dashed box) and coincident splay of the TM helices (cyan lines and angular measures). The CD3 δ and CD3 γ heterodimers are shown in (C) and (D), respectively. (**E**) Overlay of the TM helices in the detergent micelle and lipid bilayer (both ND-I and ND-II), viewed from the extracellular space showing similar overall arrangement. (**F**) Comparison of GDN (transparent gray) and ND-II (solid, color-coded) transmembrane helices aligned to TCRα residues 245-260 and viewed parallel to the membrane. Green and orange arrows highlight increased splay of the TCRβ transmembrane helix and the carboxy-terminal end of the CD3ζ TM helix, respectively. Blue arrow highlights the unstructuring and deflection of the first helical turn at the amino-terminus of the TCRα transmembrane helix in the lipid bilayer.

### Structure of the TCR–CD3 in a native-like lipid environment

To determine the structure of the 1G4 TCR–CD3 in a native-like lipid environment, we reconstituted the TCR–CD3 complex into membrane scaffold protein-based nanodiscs loaded with mammalian polar lipids and cholesteryl hemisuccinate (Fig. S4). Computational processing of cryo-EM data yielded two distinct maps determined to nominal resolutions of 3.3 to 3.1 Å (for locally refined maps used for model building and refinement; Fig. 1, S5-9; Tables S1-3). These maps, which we denote as nanodisc (ND)-I and ND-II, were clearly different from GDN (Fig. 1) and the previously published TCR structures. Attempts to bias the particle search towards the published conformations with projections of the structure in detergent or that same structure docked into a lipid nanodisc model did not yield any maps approximating the previously published conformation (Fig. S5).

Despite equal or higher nominal overall resolutions than GDN, ND-I and ND-II were not as high quality (compare local resolution estimations in Fig. S7D,H with Fig. S3D). This was likely due to a high degree of conformational variability, but also due in part to map anisotropy (compare Fig. S7C,G with Fig. S3C). Efforts to improve the orientation distribution of the particles with graphene oxide grids and/or fluorinated detergents did not improve reconstruction quality. Nonetheless, with much of the density having estimated local resolution of better than 4 Å, these maps were of sufficient quality to build models into the density.

Molecular models were built for ND-I and ND-II by docking the GDN structure into the maps and refining them by standard approaches (see Materials and Methods for details; Fig. S5-9; Tables S1-3). These models reveal two distinct but related conformations of the TCR–CD3 complex (Fig. 1A). The ND structures show two large conformational changes relative to GDN: a “closure” of the TCR ectodomains against the CD3 ectodomains (Fig. 1A,B) and a “compaction” of the juxtamembrane (JM) linker peptides (Fig. 1A,C,D). Collectively, these changes shorten the TCR projection into the extracellular milieu by approximately 35 Å. Closure of the TCR ectodomains (Fig. 1B) results in part from rotation of the variable domains towards their respective constant domains about the interdomain hinge region (Fig. S10A,B). Relative to the “extended” JM linker conformations seen in GDN, the CD3 JM linkers are compacted, with closer approximation of the CD3 ectodomains to the amino-terminal ends of their TM helices (Fig. 1C,D).

The conformational changes in the TCR ectodomains are accompanied by tilting of TM helices. While the in-plane arrangement of the TM helices is largely similar when viewed along the membrane normal (Fig. 1E), examination from orthogonal axes reveals differences in TM helix conformation. Compaction of the CD3 JM linkers requires opening of the angle between the TM helices of the CD3 δ and CD3 γ pairs by 23° (from 31° to 54°) and 28° (from 39° to 67°), respectively, when comparing ND-II with GDN (Fig. 1C,D). For ND-I, the TM helices of the CD3 δ are bent, with the first helical turn displaying a wide splay (76°, Fig. S10C), but the overall angle between the modeled helices (46°) is intermediate to ND-II and GDN. The CD3 _G_ TM helix was not modeled, as its register could not be confirmed, and thus the CD3 γ TM angle was not measured. Additionally, there is splaying of the TCRβ and distal CD3ζ TM helices away from the membrane normal (Fig. 1F). In GDN, the TCRα TM helix extends outward past the amino-terminal ends of the CD3 TM helices, where it is ensconced by the extended CD3 JM linkers. Compaction of the JM linkers in nanodiscs necessitates reorganization of the first turn of the TCRα TM helix, which is unstructured and displaced towards the lipid bilayer (Fig. 1F). Taken together, these cryo-EM maps reveal marked conformational differences between the TCR–CD3 in detergent micelles and a lipid bilayer.

Additionally, the transmembrane helices of ND-I and ND-II appear shorter than in GDN. Specifically, these structures lack clear density for the cluster of basic and aromatic amino acids visualized in GDN that extend the TM helices (Fig. S10D). In a membrane, these extensions would likely be interfacial or extend into the cytoplasm(*21*). Although there is additional density beyond the modeled carboxy-termini of some of these shortened helices in ND-I and ND-II, it is not of sufficient quality to model with confidence (Fig. S10E,F). This suggests the TCR–CD3 TM helices are less rigid in the inner leaflet of a lipid bilayer than in a detergent micelle.

### Lipid interactions with the TCR–CD3

The TCR–CD3 is known to interface directly with membrane cholesterol(*22*), and the binding sites have been visualized in previously published structures(*6*). In GDN, ND-I, and ND-II, density corresponding to two cholesterols in the previously published structures was again apparent (Fig. S10G). The two cholesterol molecules in ND-I and ND-II are oriented tail-to-tail in a groove between CD3ζ’, TCRα, TCRβ, and CD3γ, as in GDN (Fig. 2A). However, the compaction-associated changes in the TCR–CD3 TM helices result in a sliding of the cholesterol molecules towards each other, with deflection of their branched aliphatic tails to prevent clashing (Fig. 2A, insets). We refer to the cholesterol molecules as Chol_A_ and Chol_B_, corresponding to the TCR–CD3 TM helix with which each is approximated. The deflected tail of Chol_B_ is interposed between adjacent TM helices, accommodated by their increased splay in the compacted conformation (Fig. 2B, inset). The cholesterol-binding groove is more open to the membrane bilayer adjacent to Chol_B_ in nanodiscs, creating a possible path for cholesterol loading into the assembled TCR–CD3 and/or exchange with the membrane environment (Fig. 2B, bottom right).

**Fig. 2:**
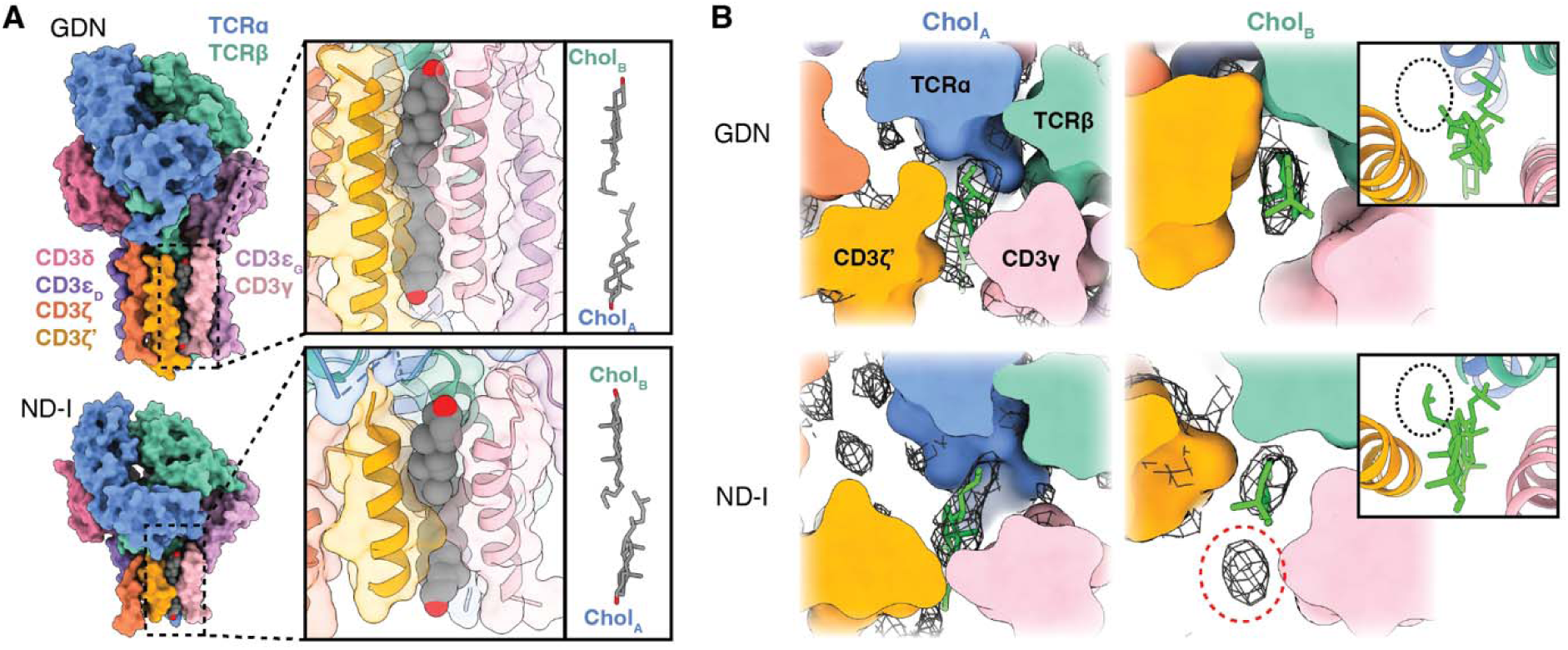
TCR–CD3–lipid interactions. (**A**) Comparison of the cholesterol-binding sites of the TCR in detergent (GDN, top) and in nanodiscs (ND-I, bottom). Overview shown at left and insets show the cholesterol-binding sites in detail (left) and with protein removed (right) to highlight the deflection of the branched aliphatic tails. Chol_B_ denotes the cholesterol proximal to the ectodomains and the TCRβ transmembrane helix; Chol_A_ denotes the cholesterol proximal to the cytoplasmic face and the TCRα transmembrane helix. (**B**) Sections parallel to the membrane plane highlighting the cholesterol-binding groove. Sections are shown at the levels of Chol_A_ and Chol_B_, as indicated. Density from locally refined and B-factor-sharpened cryo-EM maps is shown as black mesh, and cholesterols in lime green. Red dashed circle highlights structured lipid density in the nanodisc bilayer adjacent to the opened cholesterol-binding groove. Insets show the cholesterol-binding groove from the same angle as the main panels, but with both cholesterols included in the slice, highlighting intercalation of the deflected Chol_B_ aliphatic chain between TM helices in nanodiscs (black dashed oval) as compared to the structure in GDN.

Imaging of the TCR–CD3 in a lipid bilayer revealed additional sites of protein–lipid interaction. Unmodeled tubular densities consistent with phospholipid acyl chains are present at several sites adjacent to the TCR–CD3 TM helices (yellow densities in Fig. S10H-K). Structured lipid is seen parallel to the tail-to-tail cholesterol dimer, filling the aforementioned channel between the cholesterol-binding groove and the bulk membrane lipids (red dashed circle in Fig. 2B, bottom left; Fig. S10H). Splay of the CD3ζ TM helix away from that of CD3 _D_ enlarges a cavity, which is filled by several presumptive lipid acyl chains in the nanodisc (Fig. S10K). Additional probable sites of structured lipid include a shroud of density over the CD3ζ TM helices (Fig. S10I) and in the cleft over the TCRαβ TM helices between CD3δ and CD3_G_ (Fig. S10J).

The ND-I and ND-II maps show clear density for the proteolipid nanodisc that enshrouds not only the TM helices, but also the membrane-proximal faces of the TCR–CD3 ectodomains (Fig. 3A, left). The lipid contacts with the CD3 ectodomains are apparent even in maps contoured at a higher threshold, consistent with high occupancy (Fig. 3A, right). While the CD3 ectodomain surfaces are generally poorly evolutionarily conserved in comparison to the TM helices, patches on the membrane-proximal surfaces of the CD3 ectodomains do show high sequence conservation (Fig. 3A,C). These highly conserved ectodomain surface patches (black triangles in Fig. 3B-E) interface with the extracellular membrane leaflet and are neutral to electropositive (Fig. 3E), facilitating phospholipid head group interactions. Of note, we observe a similar conserved electropositive patch at the interface of CD79αβ(*23*), the analogous ITAM-bearing chains in the B-cell receptor, raising the prospect that this interface may also favor membrane interactions and a different conformation in the lipid bilayer from that reported in detergent micelles (Fig. S11).

**Fig. 3:**
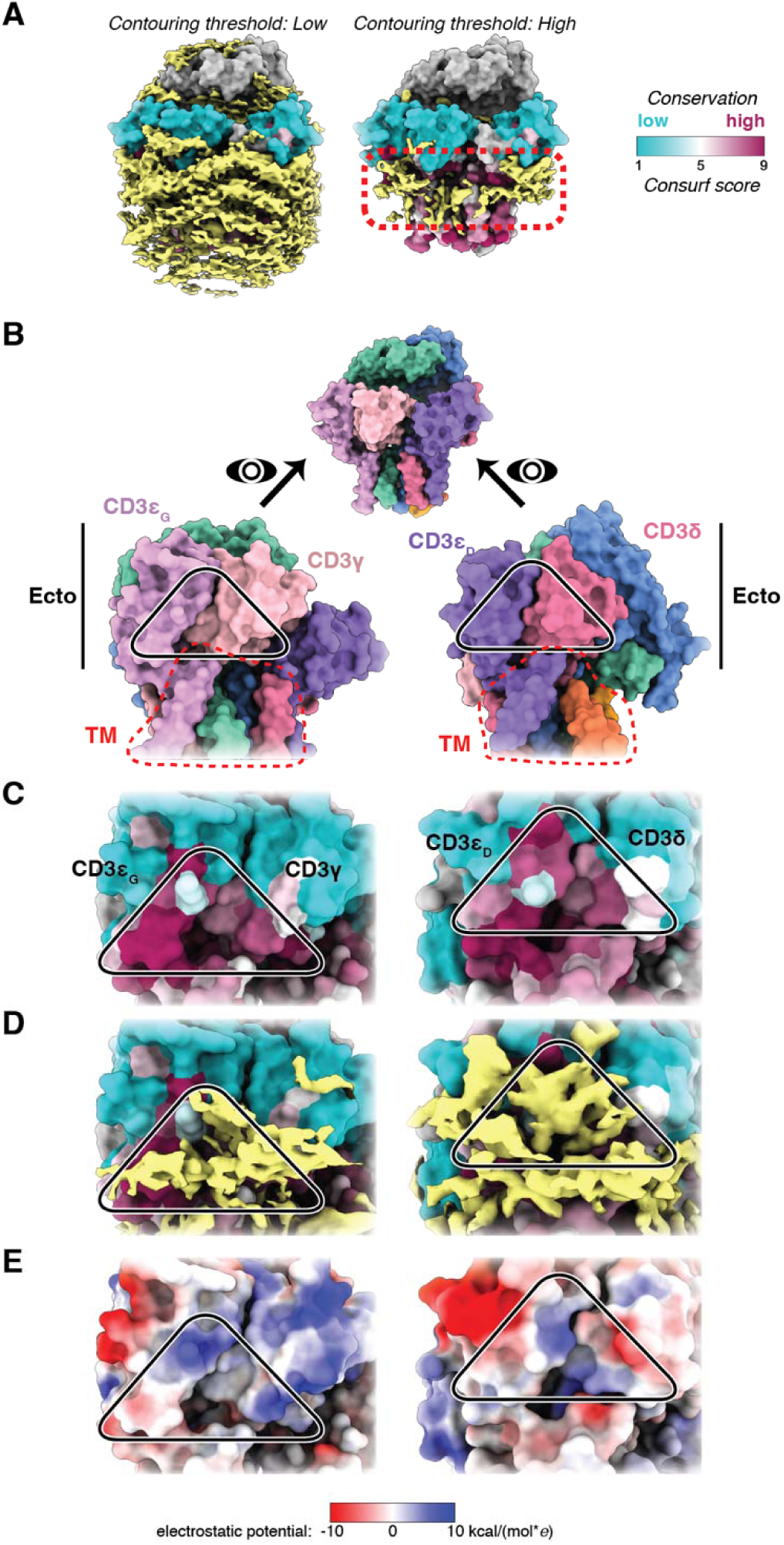
Conserved CD3 surface patches form extensive membrane contacts. (**A**) ND-II model surface colored by sequence conservation (Consurf score 1 through 9) and yellow unmodeled map density from the nanodisc enshrouding the TCR–CD3 TM helices and membrane-proximal ectodomain surfaces. When visualized with a higher contouring threshold (right), structured lipid density adjacent to the CD3 ectodomains persisted (red dotted box). Note also the relative sequence conservation in the CD3 TM helices and membrane-adjacent ectodomain surfaces, but not solvent-exposed surfaces. (**B**) Top, ND-II space-filling model for orientation. Arrows and eyes indicate the viewing direction for inspection of the CD3 γ and CD3 δ interfaces examined in the lower panels. Bottom, overviews orienting the close-up views in panels C through E. Red dashed lines denote the border between transmembrane (TM) and ectodomain (Ecto) surfaces. (**C**) Close-up views of the membrane-proximal face of the CD3 heterodimers, with black triangles highlighting the conserved surface patch at the subunit interface. (**D**) Lipid density (yellow) is most robust over this conserved patch, visualized at high contouring threshold (as in panel A, right). (**E**) The conserved ectodomain patch is relatively electropositive compared to the surrounding ectodomain surfaces colored by electrostatic potential (red to blue, −10 to 10 kcal/(mol\**e*)).

### Probing TCR conformation in situ

Which conformation of the TCR–CD3 is the physiologic resting state on the cell membrane: the open/extended conformation seen in detergent micelles, or the closed/compacted conformations seen in the lipid bilayer of the nanodisc? Given that the lipid bilayer of a nanodisc more closely approximates the physicochemical milieu of the plasma membrane, we hypothesized that the TCR–CD3 is closed and compacted in the absence of activating ligand *in vivo.* To test this notion, we engineered a series of disulfide bonds that would only be geometrically favorable in ND-I and/or ND-II but not GDN (or *vice versa*), and we read out their presence or absence by one of the following approaches: (1) non-reducing sodium dodecyl sulfate-polyacrylamide gel electrophoresis (SDS-PAGE) and Western blot of whole-cell lysates for interchain disulfides, or (2) mass spectrometry on affinity-purified protein for intrachain disulfides. Analysis of disulfide bonds in the Protein Databank have shown that disulfide bonds almost never have a Cys-Cys Cα-Cα distance greater than 7 Å, with most being between 5 and 6.5 Å(*24*). Accordingly, we sought residue pairs in the TCR–CD3 ectodomains with Cα-Cα distances below 7 Å in one conformation and above 7 Å in the other as candidates for disulfide engineering.

The interface between CD3δ and TCRα contains several pairs of residues with geometry favorable for engineered disulfide-bond formation in ND-I but not GDN (Fig. 4A). As TCRα and TCRβ are linked by an interchain disulfide natively (Fig. S12A), formation of a TCRα–CD3δ disulfide would result in an upward gel shift of the TCRαβ heterodimer in nonreducing SDS-PAGE. For all three tested engineered disulfide pairs meeting the 7 Å Cα-Cα distance constraint, a gel shift consistent with TCRα–CD3δ disulfide formation was observed (Fig. 4B, asterisks; full-length gels in Fig. S13). Confirming that this mass shift was due to disulfide formation, no gel shift was apparent under reducing conditions (Fig. S13B). Moreover, the mass shift was due to the formation of the engineered disulfide bond and not an off-target bond, as adduct formation was only seen when both residues in a given target pair were mutated to cysteines (Fig. 4B). As an internal negative control, a permutation of cysteine mutants at this interface not meeting geometric criteria for disulfide-bond formation in ND-I (TCRα S78C – CD3δ N38C, Fig. 4A, right, orange labels) was produced. It did not form a disulfide bond *in vivo* (Fig. 4B, orange) despite S78C and N38C successfully pairing with cognate cysteine mutants that did meet the inter-Cα threshold. This control shows the structural precision and specificity of the engineered disulfide bond approach. These experiments were performed using whole-cell lysates, but we obtained similar results with NiNTA affinity-purified protein (Fig. S12B), and we were able to reconstitute the two G79C-crosslinked mutants into nanodiscs (Fig. S12C,D), suggesting proper folding and assembly of the holo-TCR–CD3.

**Fig. 4:**
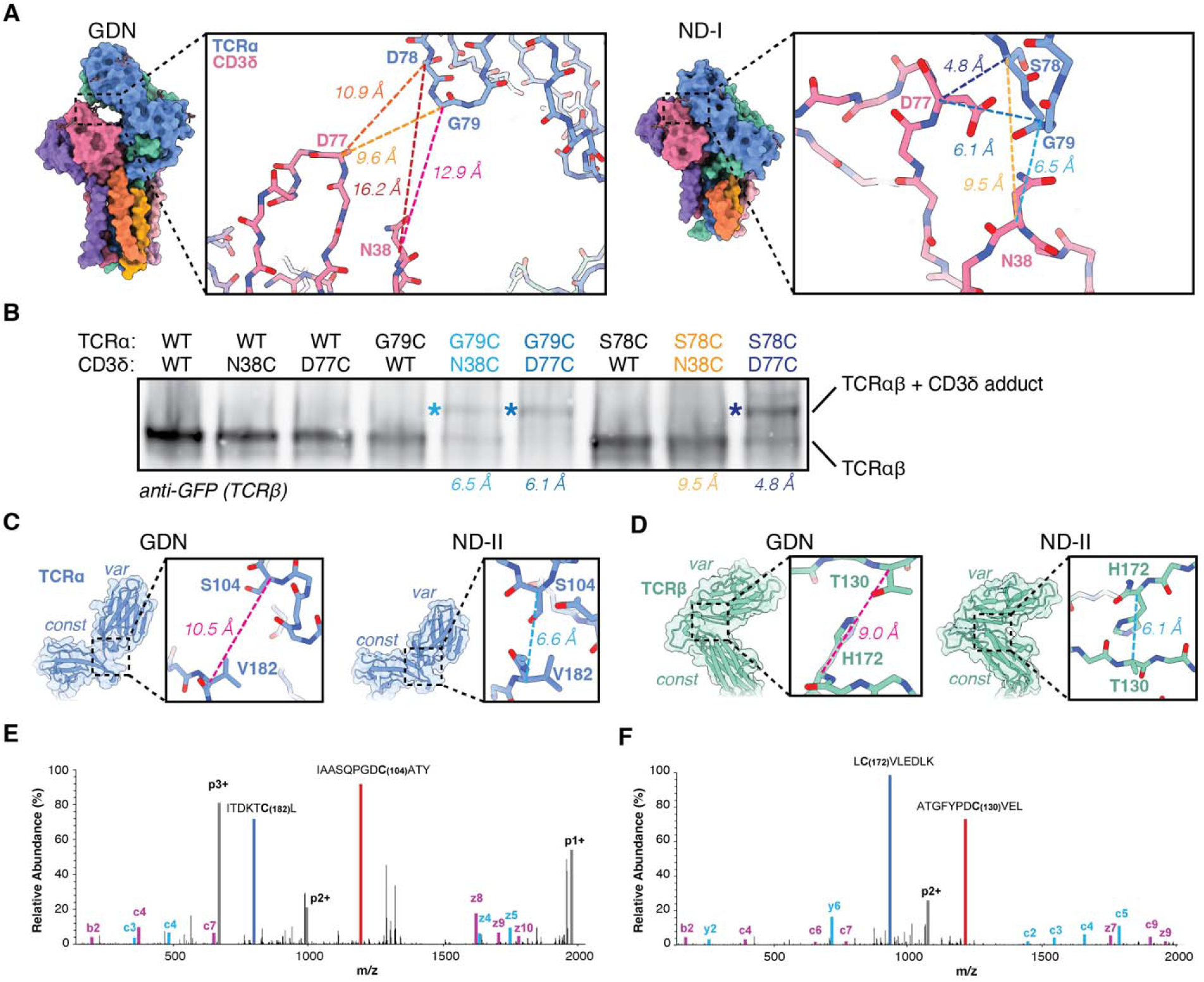
Confirmation of the TCR–CD3 resting conformation *in situ.* (**A**) Comparison of Cα-Cα distances between apposing faces of CD3δ and TCRα in detergent and in a lipid bilayer, highlighting plausible engineered disulfide positions in ND-I (distances in blue shades) that are not geometrically plausible in GDN (distances in yellow, orange, red, magenta). (**B**) Non-reducing SDS-PAGE gel of whole-cell lysates from HEK293T cells infected with the indicated TCR–CD3 virus genotypes Western blotted for EGFP-tagged TCRβ. Engineered disulfides yielding a TCRαβ–CD3δ covalent adduct resulted in an upward gel shift (asterisks). Color coding and distances as in panel A. (**C,D**) Design of intrachain disulfide bonds in the closed conformation of TCRα (C) and TCRβ (D). Distances color coded as above. (**E,F**) Mass spectrometric detection of intrachain disulfide bonds in TCRα (E) and TCRβ (F). EThcD fragment spectra of disulfide-linked dipeptides are shown. Gas-phase reduced linear peptides are shown in red (peptide A) and blue (peptide B) with the reduced Cys highlighted in bold. Fragment ions of the two linear peptides are shown in magenta (fragments of peptide A) and cyan (fragments of peptide B). Charge-reduced precursor ions are shown in gray.

The region around the TCRα and TCRβ interdomain hinges is also notably different in the closed/compacted conformation compared to the open/extended one, with the variable domains coming into closer apposition with their respective constant domains, allowing the engineering of conformation-specific disulfide pairs in this region (Fig. 4C,D). We probed formation of these intrachain disulfides by mass spectrometry. Proteolytic digestion of the TCR– CD3 with trypsin and chymotrypsin yielded distinct peptides, several of which were linked by disulfide bonds. Electron-transfer/higher-energy collision dissociation (EThcD) fragmentation was used to fragment the disulfide-bonded peptides providing *in vacuo* reduction to the linear peptide constituents along with peptide backbone fragmentation, thereby allowing unambiguous identification of the engineered disulfides(*25*). All native TCR–CD3 disulfides spanning two proteolytic fragments (Fig. S12A) were observed in the wild-type 1G4 TCR–CD3 by this approach (Table S4). For both ND conformation-compatible engineered disulfides tested (Fig. 4C,D), we observed formation of the engineered disulfide (Fig. 4E,F; Table S4). In contrast, an engineered disulfide that would only be present in the open/extended conformation (Fig. S12E) did not form *in vivo* (Table S4). Taken together, these data support the conclusion that a TCR– CD3 in the unliganded, resting state adopts the closed/compacted conformation *in vivo*.

### Conformational variability of the TCR–CD3 in a lipid bilayer

To allow direct comparison of the two distinct, but related, closed/compacted structures (ND-I and ND-II), they were aligned by their TCRα TM helices. TCRα was chosen because it is relatively perpendicular to the membrane plane (alongside CD3ζ’) and thus might be least affected by any differences in TM helix splay. Indeed, this alignment strategy brought the nanodisc density of their maps into alignment as well (Fig. S14A), giving both models the same orientation relative to the membrane bilayer. In contrast to the strong overall alignment of the TCR–CD3 TM helices by this approach (Fig. S14B,C), the ectodomains of ND-I and ND-II have marked conformational differences, as seen in the aligned cryo-EM densities (Fig. S14A,B) and the resulting protein models (Fig. 5A). Comparison of the structures reveals the motions allowing exchange between these conformations (Fig. 5B,C; Movie S1). ND-I is the most closed of the three structures, and transition to ND-II requires several motions: the depression of the CD3 _D_ (Fig. 5B) and CD3γ ectodomains (Fig. 5C) away from the TCR variable domains and towards the bilayer outer leaflet; rotation of the of the TCRα variable domain about the interdomain hinge (Fig. 5B); and retraction of the TCRβ variable domain (Fig. 5C). This is accompanied by sliding of CD3ζ along its helical axis towards the ectodomains (Fig. 5B). While the TCRα and CD3ζ’ TM helices remain relatively fixed in this frame of reference, the TCRβ and CD3γ TM helices shift away from the cholesterol-binding groove (Fig. 5D, S14C). To accommodate this movement, there is rotation of Chol_A_ and elevation of Chol_B_ (Fig. 5D), showing cholesterol to be an integral and dynamic component of the TCR–CD3 structure.

**Fig. 5:**
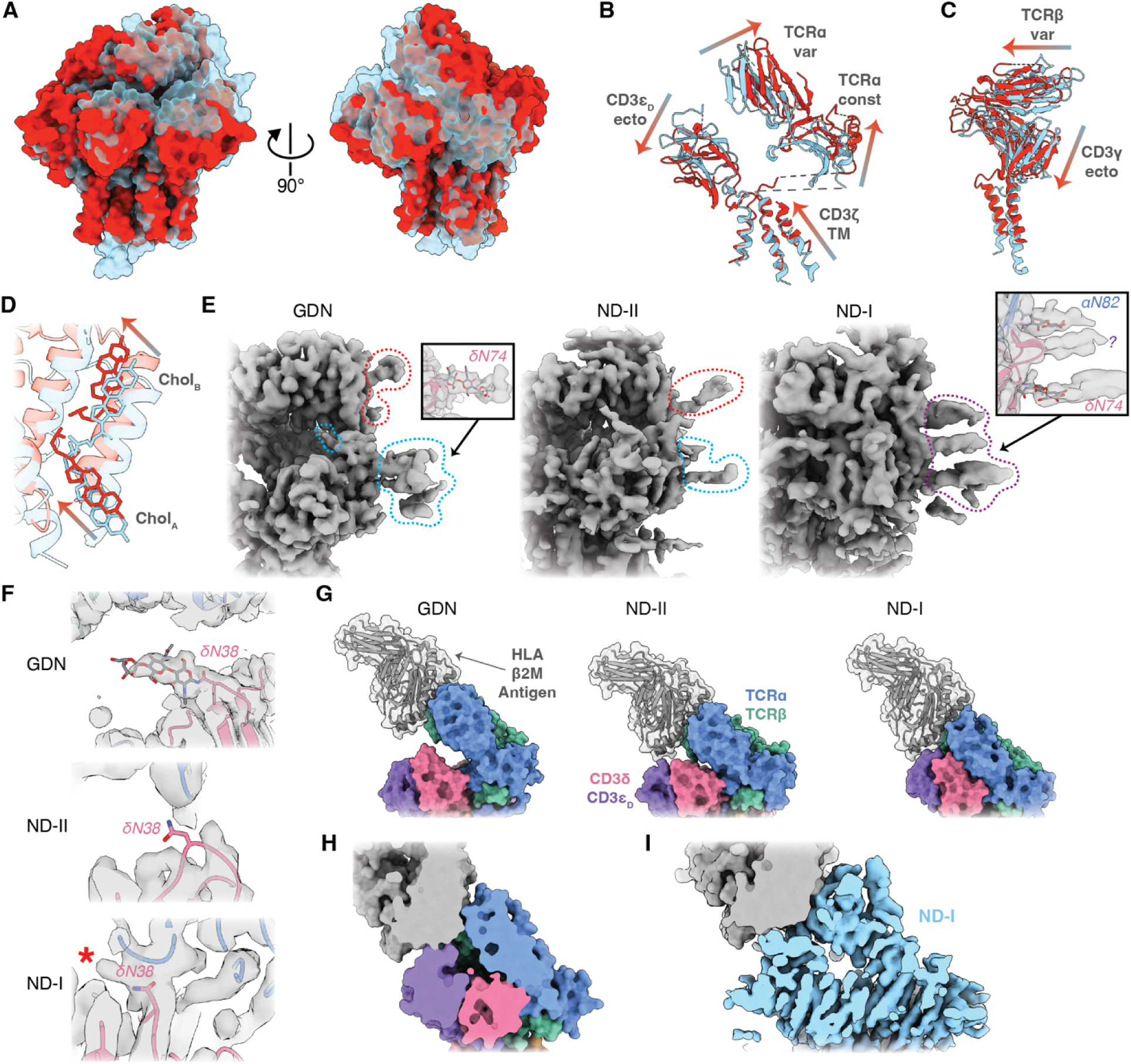
Conformational variability of the TCR–CD3 in the resting state. (**A**) Comparison of ND-I (transparent cyan) and ND-II (red), shown here as model surfaces, aligned on the TCRα transmembrane helix. (**B,C**) Gradient arrows highlight the domain movements underlying the transition between ND-I (cyan) and ND-II (red). (**D**) TCRβ and CD3γ TM helix movement away from the cholesterol-binding site during the transition from ND-I to ND-II, with arrows indicating the accompanying movements of the cholesterols in the sterol-binding groove. (**E**) Comparison of the TCRα and CD3δ lateral glycosylation sites (red and cyan dashed lines, respectively) in the density maps showing increasing ordering of the glycan array with ectodomain closure (purple dashed line). Insets highlight modeled asparagine-linked glycan cores (or partial cores) modeled into these densities in GDN and ND-I. (**F**) Comparison of the CD3δ N38 residue in GDN, ND-II and ND-I. Gray surface is the locally refined, DeepEMhancer-sharpened cryo-EM density. (**G**) Modeling HLA–β2M–antigen (gray) binding to the TCR in the indicated conformations. (**H**) Cross section through the HLA–TCR interface in ND-I shows close association of HLA and CD3 _D_, but no large clashes. (**I**) Similarly, modeling HLA onto the locally refined, DeepEMhancer-sharpened ND-I density does not show any large clashes between HLA and unmodelled density.

The GDN map displays densities for many of the known asparagine (N)-linked glycans in the TCR–CD3 (Fig. S14D). There were fewer densities modellable as N-linked glycans in the ND-I and ND-II maps (Fig. S14D). However, all three maps show an array of densities lateral to the CD3δ and TCRα ectodomains, corresponding to sites of known N-linked glycosylation. In ND-II, the TCRα (linkage unclear) and CD3δ N74-linked glycans are visible, emerging radially from the TCR–CD3, as in GDN, but are not well structured (Fig. 5E). By comparison, ND-I reveals clear reorganization of the glycans (Fig. 5E), with increased stacking of adjacent glycans brought into close proximity by tight ectodomain closure. This arraying of N-linked glycans is reminiscent of that seen on HIV Env(*26*), where interacting adjacent glycans form a structured arcade of glycan cores (Fig. S14E). Two densities in the array are attributable to CD3δ N74- and TCRα N82-linked glycans, with a third density emerging between them without definitive attribution (purple question mark in Fig. 5E, right inset). In GDN, as in other structures of TCR– CD3 in detergent, the CD3δ N38-linked glycan is oriented inward towards the cleft between the TCR and CD3 ectodomains (Fig. 5F, top). In ND-II, CD3δ N38 is unstructured (Fig. 5F, middle), but is grossly positioned at the entrance to the cleft, and the linked glycan could conceivably orient either into or away from the cleft. In ND-I, CD3δ N38 is structured (Fig. 5F, bottom), but its glycan cannot be confidently modeled. However, density protrudes from N38 into the open cleft (Fig. 5F, asterisk) and additional unmodeled density is seen within the cleft, consistent with the glycan remaining within the cleft in the ND-I conformation.

The differences in ectodomain closure seen between ND-I and ND-II could have consequences for the accessibility of the TCR variable domains to HLA–antigen complexes. To assess the impact of ectodomain cleft opening and closure on TCR–HLA interactions, we modeled the binding geometry of HLA onto GDN, ND-I, and ND-II by alignment of the TCR-HLA crystal structure to the TCR variable domains in our models (Fig. 5G). There appears to be adequate clearance for the TCR variable domains to interact with HLA in GDN and ND-II. For ND-I, the modeled fit is tighter, with the CD3 _D_ ectodomain predicted to be in close contact with HLA. However, on assessment of space-filling model cross-sections, no overt steric clashes are evident (Fig. 5H). Because a portion of a potentially clashing surface loop in the CD3 _D_ ectodomain is not modeled in ND-I, the TCR-aligned HLA model was docked into the ND-I cryo-EM density (Fig. 5I), but no unmodeled density appears to clash with HLA. The CD3 _D_ ectodomain from GDN is completely modeled and aligns closely with that from ND-I (Fig. S14F, inset). As an additional sensitivity analysis, the GDN CD3 _D_ ectodomain was modeled into ND-I and the HLA interaction again assessed. While there is a small overlap in their space-filling models (Fig. S14F), no large clashes are observed. Thus, while it remains unclear what quantitative effect ectodomain cleft closure might have on HLA affinity, particularly in the physiologic context of a glycosylated HLA molecule, it is at least plausible that both ND-I and ND-II may be receptive to HLA binding.

While our modeling data suggest that HLA can bind to TCR–CD3 in the closed/compacted resting state conformations, TCR–CD3 opening and/or extension may be necessary for coreceptor interaction with HLA bound to the TCR–CD3. Modeling the HLA class II–CD4 complex onto GDN, ND-I, and ND-II reveals an unfavorable geometry for CD4 interaction with a class II HLA when bound to a TCR–CD3 in the ND-I or ND-II conformation because of the proximity of the TCR variable regions to the membrane (Fig. S14G), at least for the CD4 conformation reported in the CD4-HLA-TCR ectodomains ternary complex crystal structure(*27*). In contrast, CD4–HLA can be accommodated in the open/extended GDN conformation (Fig. S14G). The full ectodomain structure of the CD8 coreceptor has not been described, precluding a similar analysis applicable to TCRs recognizing HLA class I-restricted epitopes, like the one used in this study.

### TCR–CD3 conformational change upon HLA binding

Whether or not the TCR–CD3 complex undergoes conformational change upon binding to HLA remains an open question. To determine the structure of the HLA-liganded TCR–CD3 complex in a lipid bilayer, we reconstituted TCR–CD3 in nanodiscs as above, incubated the complex with an excess of single-chain trimeric HLA–β2M–NY-ESO-1(C9V) (herein, “HLA”), and determined the structure by single-particle cryo-EM (Fig. S15, S16; Tables S1-3). Despite picking particles using a template model of the closed/compacted TCR–CD3 complex bound to HLA, cryo-EM reconstruction showed the HLA-bound complex to be in the open/extended conformation (Fig. 6A,B). Indeed, the HLA-bound conformation was nearly identical to that seen in GDN (Fig. 6C). There was subtle tilting of HLA relative to the TCR when comparing the HLA-bound nanodisc structure to the crystal structure of the 1G4 TCRαβ ectodomain heterodimer bound to HLA, but the binding interface was otherwise similar (Fig. S17A).

**Fig. 6:**
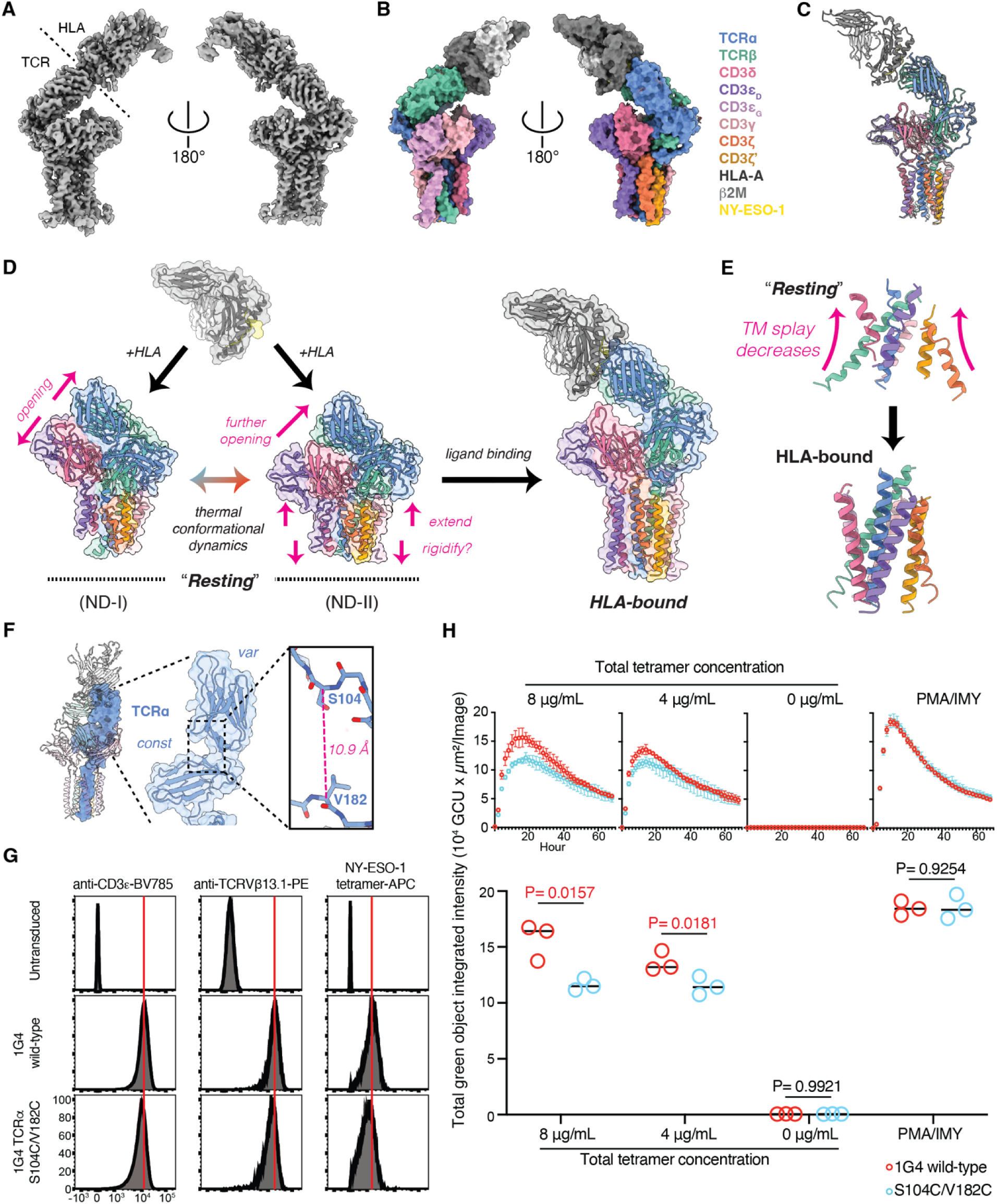
Structure of the HLA-bound TCR–CD3 complex in nanodiscs and model of TCR– CD3 activation. (**A,B**) Cryo-EM maps (A) and model surfaces (B) of HLA-bound TCR–CD3 in nanodiscs. Maps shown in (A) were DeepEMhancer-sharpened after local refinement to remove nanodisc density. Legend in (B) applies to (B) – (F). (**C**) Comparison of GDN and HLA-bound models, highlighting the similarity in TCR–CD3 conformation. HLA-bound model is colored by chain and GDN is transparent gray. (**D**) Proposed model of TCR–CD3 conformational dynamics. The unliganded TCR–CD3 explores resting subconformations (between ND-I and ND-II). HLA binding induces a conformational transition from ND-II to an open/extended conformation. Magenta arrows and text indicate movements required to transition to the next conformation in the series. (**E**) Changes in the TM helices between “resting” (ND-I shown) and HLA-bound conformations. (**F**) Increasingly zoomed-in views of the TCRα variable and constant domains in the HLA-bound conformation highlighting the Cα-Cα distance between S104 and V182. (**G**) Flow cytometry histogram from J8Zb2m-α-β-cells post tetramer-APC enrichment and stained as indicated (n = 3 replicates). Cells were untransduced (top) or transduced with a virus expressing the TCRαβ 1G4 genotype indicated (middle, bottom). Vertical axis is cell count normalized to peak; horizontal axis is fluorescence in arbitrary units. Red line indicates fluorescence intensity mode (histogram peak) for wild-type 1G4. (**H**) Time-dependent T-cell activation quantified using an NFAT-ZsGreen reporter by live cell imaging. Top shows ZsGreen fluorescence over time after addition of indicated concentrations of HLA-tetramer or PMA/ionomycin for 1G4 wild-type (red) or TCRα S104C/V182C mutant-transduced cells (cyan). Data shown are mean +/- standard deviation using biologic triplicates. Peak TCR–CD3 activation amplitudes are compared for each condition at the 8 μg/mL HLA-tetramer peak (t=16h) for all HLA-tetramer concentrations or at the PMA/Ionomycin peak (t=10h) specifically for that condition (bottom). P values calculated by unpaired, two-tailed t-test shown for each comparison. P < 0.05 highlighted in red.

Comparison of the TCR variable domains in the HLA-bound nanodisc and GDN structures showed them to be nearly identical, despite the absence of ligand in the GDN structure, and both were similar to the crystal structure of the 1G4 TCRαβ ectodomain heterodimer bound to HLA (Fig. S17B). By contrast, ligand binding induced a number of structural changes in the TCR variable domains relative to ND-I and ND-II. Specifically, in ND-I and ND-II the amino-terminal strands and several surface loops in the variable domains were unstructured, but these were ordered in the HLA-bound structure (Table S3). Qualitatively, the HLA-bound map showed better resolved protein density throughout (compare Fig. 6A with Fig. 1A), consistent with a general rigidification of the TCR structure upon HLA binding, as has been observed previously for the TCR ectodomains by NMR(*15*) and hydrogen-deuterium exchange(*28*). This rigidification was apparent in the density corresponding to the TCR variable domains, particularly for TCRβ (Fig. S17C,D).

Structural comparison of the resting and HLA-bound TCR–CD3 in lipid bilayers demonstrates that HLA binding is sufficient to induce substantial conformational changes in the TCR–CD3 complex. Our data support a model in which the unliganded “resting” TCR–CD3 explores a thermally accessible spectrum of closed/compacted conformations from the most closed ND-I to the more open ND-II, with HLA-binding triggering a change to the open/extended conformation (Fig. 6D). We favor this transition occurring via the ND-II conformation, which is more open than ND-I and thus closer to the HLA-bound state along this trajectory. Morphing the closed/compacted conformation to the open/extended one (Movie S2) highlights the straightening of the TM helices towards the membrane normal (that is, decreased splay, Fig. 6E). These changes in TM conformation add to the plausibility of the model, as they provide a means to transmit conformational changes in the ectodomains to the cytoplasmic ITAMs.

While some published data support pure allosteric activation models for the TCR, in which HLA binding is sufficient to induce T-cell receptor activation(*29*), other data support models in which ligand-induced conformational change is necessary but not sufficient, requiring subsequent receptor clustering(*12*) or mechanical force application(*30*). However, all of these models propose that TCR–CD3 conformational change is a necessary step in the activation process, and thus we hypothesized that constraining conformational change would impair ligand-induced T-cell activation. Transition from the closed/compacted to the open/extended conformations revealed here requires further ectodomain opening from the ND-II state (Fig. 6D). We hypothesized that the intra-TCRα disulfide bond-forming double mutation S104C–V182C (characterized above, Fig. 4C) would prevent ectodomain opening to the HLA-bound conformation, as the Cα-Cα distance between these residues in that conformation is outside the limits of a disulfide bond (Fig. 6F). To investigate the impact of restrained TCR–CD3 opening on ligand-dependent activation of T cells, we introduced wild-type and TCRα S104C–V182C mutant TCR constructs into TCRαβ-negative Jurkat T cells bearing an NFAT-responsive fluorescent reporter. TCR–CD3 assembly and surface localization (as assessed by anti-CD3 antibody binding) and HLA-binding competence were confirmed by flow cytometry for the mutant and were comparable to cells transduced with wild-type 1G4 (Fig. 6G). In the absence of HLA–antigen tetramers, there was no detectable reporter fluorescence in these cells, but upon exposure to HLA–antigen tetramers, cells expressing wild-type TCR–CD3 became activated, with reporter fluorescence peaking 16 h after HLA-tetramer addition (Fig. 6H). By contrast, peak reporter fluorescence was significantly decreased in the TCRα S104C–V182C mutant, with peak activation attenuated by 26% relative to wild-type on exposure to 8 μg/mL of tetramers (P < 0.05, Fig. 6H; source data in Table S5). The TCRα S104C–V182C mutant and wild-type controls were equally responsive to phorbol 12-myristate 13-acetate (PMA) and ionomycin, which bypass TCR–CD3 signaling to activate T cells, showing this phenotype to be specific to ligand-induced TCR–CD3 activation.

## Discussion

Here, we report structures of a human TCR–CD3 both in a lipid bilayer and in detergent micelles. We find that the TCR–CD3 hetero-octamer assumes a spectrum of more closed and compacted conformations when embedded in a lipid bilayer relative to that seen in detergent micelles, and that the closed/compacted conformation is indeed the native unliganded state of the TCR–CD3. Addition of HLA, the physiologic ligand for the TCR, induced a change in conformation from the resting state to an open/extended conformation like that previously seen in detergent, and exit from the resting conformation by ectodomain opening is necessary for maximal ligand-dependent activation of the TCR–CD3 complex. While this latter finding does not rule out roles for TCR–CD3-extrinsic phenomena (e.g., higher-order clustering, steric exclusion of inhibitory phosphatases) in the activation of the TCR–CD3 or regulation thereof, it argues that intrinsic conformational change in the TCR–CD3 is necessary for early TCR–CD3 activation and argues against previous models that have excluded this requirement(*11*, *31*).

Similarly, it does not rule out a role for mechanical force application or catch bond formation in the maximal activation of the TCR–CD3 but shows that the earliest stage of TCR activation (i.e., conformational change out of the resting state) can be mediated by ligand binding alone.

Broadly, these findings reinforce the importance of structural interrogation of membrane proteins in lipid environments that closely resemble their physiologic environment. Comparison of TCR–CD3s in detergent micelles in the absence and presence of activating ligand had revealed no change in TCR–CD3 structure(*7*, *8*), consistent with models excluding TCR–CD3 conformational change in the activation process. Although these studies were technically well performed, our results now reveal that they did not identify the true resting state of the TCR– CD3 complex in lipid bilayers, preventing observation of the conformational changes during ligand-dependent activation.

Additionally, the structures in detergent did not capture the full scope of TCR–CD3 interaction with the membrane. Not only are membrane lipids closely interwoven with the TM helices, but large conserved surface patches on the ectodomains mediate lipid–protein interactions, likely partially counteracting the thermodynamic costs of JM linker compaction and “spring loading” the TCR with potential energy for conformational change during activation. In this model, ligand binding provides the activation energy to release the membrane-binding surfaces of the CD3 ectodomains and allow JM extension. Detergent extraction would overcome this thermodynamic barrier by removing the lipid bilayer all together. During reconstitution into nanodiscs, we first extracted the TCR–CD3 complex in detergent, opening and extending the receptor, which suggests that the addition of a lipid bilayer is sufficient to reset the TCR–CD3 to its resting conformation without the need for extrinsic chaperones. Lastly, we note that a similar conserved surface patch is present on the CD79 heterodimer in the B-cell receptor(*23*), raising the possibility that other hetero-multimeric ITAM-containing immunoreceptors exploit protein surface interactions with the lipid bilayer to constrain conformation.

As with the TCR–CD3 structures in detergent, the CD3 ITAMs are not visible in our structures. The ITAMs are membrane-associated in the unliganded state(*9*, *10*), but are likely to be radially disordered along the intracellular membrane face. However, the TM helix movements seen between our closed/compacted and the open/extended conformations suggest a means for communication of extracellular conformational changes in the TCR–CD3 complex to the ITAMs. Within the cytoplasmic domains of CD3 proteins, the basic residues proximal to the CD3 TM helix are critical for ITAM association with the membrane(*9*, *10*). Upon transition from the receptive to the HLA-bound conformation, the decreased splay in the CD3 TM helices could alter basic patch interaction with the membrane and favor ITAM release.

Analysis of the basal conformational heterogeneity of the TCR–CD3 suggests novel means of TCR–CD3 regulation. The TCRα and CD3δ glycan ordering observed in the ND-I conformation is notably different from the glycan arrangement in the more open ND-II conformation or the TCR–CD3 in detergent. While molecular modeling of the branched glycan sugars was not possible with sufficient confidence, the stacked arrangement of glycan core densities raises the possibility of extensive hydrogen bonding between the glycan branches distally and perhaps also with TCR–CD3 surface residues. While glycan–glycan and glycan– protein interactions are well described *in trans*, there are fewer examples of structured glycan– glycan interactions within a given protein or stable complex. The human immunodeficiency virus-1 Env protein is an example of the latter(*26*, *32*) and the arrangement of the radial glycan array in ND-I is reminiscent of that seen in crystallographic studies of Env. However, the stacking of glycans observed in the TCR–CD3 is unique in being protein conformation-dependent. Breaking an extensive glycan hydrogen-bonding network to transition out of ND-I would be energetically costly, and thus modification of TCR–CD3 glycosylation could serve as a means of posttranslational regulation of the TCR–CD3 activation threshold by increasing or decreasing the free energy barrier to ND-I exit.

Previous studies have explored the role of cholesterol in the regulation of TCR–CD3 activation(*6*, *18*, *22*). The substantive movements of the two cholesterol molecules seen between TCR–CD3 conformations raise the possibility that changes in the sterol composition of the T-cell membrane (either global or local) could alter TCR–CD3 activity even prior to ligand binding by shaping the basal TCR–CD3 conformational landscape. Moreover, our structures reveal channels for cholesterol loading or exchange with the lipid bilayer not previously observed, making it plausible that already assembled TCR–CD3 could respond to changes in membrane sterol content. Non-sterol lipid species are also known regulators of T-cell signaling(*17*), including phosphoinositol lipids(*33*, *34*) and oxidized lipids in the tumor microenvironment(*35*). This work delineates multiple sites of structured interaction between the TCR–CD3 and membrane lipids beyond the sterol-binding site, which may allow specific lipids to modulate TCR–CD3 conformation and activity.

The extensive contacts between TCR variable domain residues and those of the TCR constant domains or CD3 ectodomains in the closed/compacted conformations raises questions about how variation at these sites might confer specific baseline sensitivities to given TCR alleles. Whether use of certain V or J alleles changes the occupancy of different segments of the TCR–CD3 conformational landscape will need to be determined and has broad implications for vaccine design and immunotherapy engineering. Likewise, whether individually rare germline coding variants in the TCR constant domain *loci* or CD3 proteins affect an individual’s basal T-cell reactivity is unclear outside of known deleterious loss-of-function alleles(*36*). Such information might predict individual propensity for autoimmunity or responsiveness to immune-oncology agents and will provide insights for engineering TCRs with favorable activity profiles(*37*).

## Acknowledgements

The authors thank the members of the Walz, T. de Lange (RU), and R. K. Hite (MSKCC) labs and W. D. Tap (MSKCC) for insightful discussions. We thank M. Ebrahim, J. Sotiris, and H. Ng at the Evelyn Gruss Lipper Cryo-EM Resource Center of The Rockefeller University (RRID:SCR_021146) for assistance with cryo-EM data collection. Mass spectrometry data were generated by the Proteomics Resource Center at The Rockefeller University (RRID:SCR_017797).

## Funding

NIH T32CA9207 (RQN)

NIH KL2TR001865 (RQN)

NIH UL1TR001866 (RQN)

The Black Family Metastasis Center at Rockefeller University (RQN)

Shapiro-Silverberg Fund for the Advancement of Translational Research (RQN)

The Mark Foundation for Cancer Research (RQN and TW)

NIH R37 CA259177 (CAK)

NIH R01 CA269733 (CAK)

NIH R01 CA286507 (CAK)

NIH P30 CA008748 (CAK)

NIH P50 CA217694 (CAK)

Cycle for Survival (CAK)

The Shteinbuk and Mead Family (CAK)

Metropoulos Family Foundation (CAK)

The Parker Institute for Cancer Immunotherapy (CAK)

The Sarcoma Center at MSKCC (CAK)

Sohn Conferences Foundation (HM)

Leona M. and Harry B. Helmsley Charitable Trust (HM)

## Author contributions

Conceptualization: RQN, TW

Methodology: RQN, FY, SH, MWB, ZM, HM, CAK, TW

Investigation: RQN, RY, SH, MWB, ZM, PD

Visualization: RQN, FY, SH, MWB

Funding acquisition: RQN, HM, CAK, TW

Project administration: RQN, HM, CAK, TW

Supervision: RQN, HM, CAK, TW

Writing – original draft: RQN

Writing – review & editing: RQN, FY, SH, MWB, HM, CAK, TW

## Data and materials availability

Cryo-EM maps and atomic coordinates for GDN, ND-I, ND-II, and HLA-bound have been deposited in the Protein Data Bank and Electron Microscopy Data Bank with the following accession numbers: GDN, PDB 9BBC, EMD-44417; ND-I, PDB 8TW4, EMD-41658; ND-II, PDB 8TW6, EMD-41660; HLA-bound, PDB 9C3E, EMD-45166. Novel plasmids and cell lines are available by material transfer agreement with the respective institutions.

## Supplementary Materials

Figs. S1 to S17 Tables S1 to S8

Movies S1 to S2

## Materials and Methods

Protein sequences, plasmids, and antibodies used in this study are listed in Tables S6-8.

### Cell lines

Eukaryotic cells lines were maintained under standard sterile conditions using aseptic technique with the following parameters: SF-9 cells (ATCC CRL-1711) were maintained at 5e5-10e6 cells/mL in Sf-900 II SFM medium (Thermo Fisher Scientific) supplemented with antibiotic-antimycotic solution (Thermo Fisher Scientific) at 27°C, ambient CO_2_. HEK293GP cells (Takara Bio, 631458) were maintained at 10-90% confluency in RPMI medium 1640 (RPMI, Thermo Fisher Scientific) supplemented with 10% fetal bovine serum (FBS, Sigma Aldrich), 10 mM HEPES, pH 7.5, and penicillin/streptomycin at 37°C, 5% CO_2_. HEK293F FreeStyle cells (Thermo Fisher Scientific) were maintained at 5e5 to 2e6 cells/mL in FreeStyle 293 expression medium (Thermo Fisher Scientific) supplemented with antibiotic-antimycotic solution at 37°C, 8% CO_2_. Modified Jurkat T cells (see below) were maintained at 2e5 to 2e6 cells/mL in RPMI medium supplemented with 10% FBS and penicillin-streptomycin solution at 37°C, 5% CO_2_.

### Plasmids and sequences

A modified version of the BacMam expression system(*38*) was used for the expression of the TCR–CD3 complex in HEK cells with furin cleavage and P2A ribosomal skipping sites separating each protein sequence (see “Protein expression and purification” and “Engineered disulfide crosslinking”). The modified viral vector also expressed the VSV-g sequence (kind gift from Titia de Lange, Rockefeller University) under a baculoviral promoter to enhance mammalian cell tropism. A self-inactivating retroviral plasmid encoding the 1G4(LY) TCRαβ sequence (“wild type” for this study) for transduction of SUP-T1 cells was a kind gift from S. A. Rosenberg, NIH. CD3ζ was cloned from Jurkat cell cDNA using standard approaches. CD3, CD3γ, CD3δ, EGFP, mCherry, and 8xHis tags were cloned from synthetic double-stranded DNA blocks (IDT DNA). Mutant TCR–CD3 sequences were produced by overlap-extension PCR or synthesized *de novo* (Genscript). Protein sequences used in this study are listed in Table S6, and plasmids used in this study are listed in Table S7.

### Baculovirus production

Bacmid recombineering was performed as per manufacturer specifications with the FastBac system in DH10Bac cells (Thermo Fisher Scientific). Bacmid was purified by phenol/chloroform/isoamyl alcohol extraction, and 25 mg bacmid DNA was transfected into SF-9 cells adherent in 6-well plates using Cellfectin II reagent (Thermo Fisher Scientific) in Grace’s Insect Medium (Thermo Fisher Scientific) for 6 h at 27°C before transfer to standard SF-9 medium (above). After 5 days, the cell supernatant (P1 virus) was removed and used to inoculate a 40-mL culture of SF-9 cells at 1e6 cells/mL. After >50% of the cells had died (∼day 5), supernatant (P2 virus) was harvested by centrifugation of the culture at 3,500xg for 20 min, decanted, supplemented with 2% FBS, and stored in the dark at 4°C. For large-scale expression, P3 virus was produced by inoculating 40-mL cultures of SF-9 cells at 3e6 cells/mL with 2 mL P2 virus and cultured, harvested, and stored as for P2 virus.

### Protein expression, purification, and nanodisc reconstitution

See overview schematic in Fig. S4A. For structural studies, ∼250-mL cultures of HEK293F FreeStyle cells at 2e6 cells/mL shaking in 1-L baffled polycarbonate flasks were inoculated with 15 mL each of P3 virus, and sodium butyrate (Alfa Aesar) was added to a final concentration of 10 mM to enhance expression. For preparation of the TCR–CD3 in detergent for cryo-EM, TCR–CD3 was expressed using two baculoviruses: (1) TCRα-[furin/P2A]-TCRβ::mCherry, and (2) CD3 - [furin/P2A]-CD3ζ-[furin/P2A]-CD3δ-[furin/P2A]-CD3γ::8HIS. Cells were incubated at 30°C, 5% CO_2_ for 3 days to allow expression and harvested by centrifugation at 4,000xg for 10 min at 4°C. Cell pellets were resuspended in 25 mM HEPES (Sigma Aldrich), pH 7.5, 1 mM phenylmethanesulfonylfluoride (Sigma Aldrich) and lysed with a Dounce homogenizer. Lysate was clarified by centrifugation at 4,000xg for 10 min at 4°C, and the supernatant was decanted and ultracentrifuged at 100,000xg for 1 h at 4°C to recover the membrane fraction. Membrane pellets were resuspended in 25 mM HEPES, pH 7.5, 150 mM NaCl (Thermo Fisher Scientific), (herein, HBS), and n-dodecyl-β-D-maltoside (DDM, Anatrace) and cholesteryl hemisuccinate (CHS, Anatrace) were added to final concentrations of 1% and 0.1%, respectively. The membrane pellets were solubilized on a rotator for 2 h at 4°C. Debris was removed by centrifugation at 4,000xg for 30 min at 4°C. The clarified supernatant was loaded onto NiNTA agarose resin (Qiagen). Resin was washed with 50 column volumes of HBS with 0.02% glyco-diosgenin (GDN) and 30 mM imidazole (Thermo Fisher Scientific). The TCR–CD3 was eluted from the NiNTA-agarose resin with 360 mM imidazole and 0.02% GDN in HBS. The mCherry fusion to TCRβ was removed by incubation with human rhinovirus protease 3C overnight at 4°C. Protein was concentrated with Amicon centrifugal filters (Millipore) and further purified by size-exclusion chromatography in HBS with 0.02% GDN over a Superose6 column (GE Healthcare).

For expression of the TCR–CD3 for affinity purification and nanodisc reconstitution, we found that co-expression of three viral constructs gave acceptable yields of material for cryo-EM analysis: (1) TCRα-[furin/P2A]-TCRβ::mCherry-[furin/P2A]-CD3δ-[furin/P2A]-CD3γ-[furin/P2A]-CD3-[furin/P2A]-CD3ζ::EGFP, (2) CD3δ-[furin/P2A]-CD3γ::8HIS, and (3) CD3-[furin/P2A]-CD3ζ::EGFP. Cells were cultured, infected, harvested, and lysed as above. Lysate was clarified, membranes isolated and extracted, and extracts loaded on to NiNTA-agarose resin as above. Resin was washed with 50 column volumes of HBS with 0.1% DDM, 0.01% CHS, and 30 mM imidazole. The resin was resuspended in HBS with 0.1% DDM, 0.01% CHS for on-bead nanodisc reconstitution. For the TCR–CD3 complex in nanodiscs subsequently incubated with HLA–β2M–NY-ESO-1, a simplified two-virus expression scheme was used: (1) TCRα-[furin/P2A]-TCRβ::EGFP, and (2) CD3-[furin/P2A]-CD3ζ-[furin/P2A]-CD3δ-[furin/P2A]-CD3γ::8HIS. Protein complexes were otherwise purified as above.

T-cell subsets differ in plasma membrane lipid composition, and individual T cells have spatially heterogenous membrane compositions(*17*). To create a lipid environment with a diversity of lipid constituents, including the negatively charged phosphatidylserine and phosphatidylinositol lipids important for ITAM–membrane interactions(*9*, *10*), porcine brain polar lipids (Avanti Polar Lipids #141101C) were used to reconstitute the lipid bilayer. Avanti reports the composition of this lipid mixture as follows: 12.6% phosphatidylcholines, 33.1% phosphatidylethanolamines, 4.1% phosphatidylinositides, 18.5% phosphatidylserines, 0.8% phosphatidic acid, 30.9% unknown. The porcine brain polar lipids dissolved in 10% DDM were added to a final concentration of 33 μg/mL (approximately 43 μM using the molecular weight of 1-palmitoyl-2-oleoyl-glycero-3-phosphocholine as a standard phospholipid) and membrane-scaffold protein (MSP) 1D1DH5 was added to a final concentration of 20 μM. MSP 1D1DH5 was expressed and purified as previously described(*39*). Components were allowed to incubate for 15 min on ice before addition of SM-2 BioBeads (Bio-Rad) to approximately ¼ of the reaction volume. The mixture was incubated overnight on a rotator at 4°C. The reaction mixture was again returned to a disposable plastic column and washed with 20 column volumes of HBS, 50 column volumes of HBS with NaCl increased to 300 mM and 30 mM imidazole, washed with 20 column volumes of HBS, then eluted in HBS with 360 mM imidazole. Protein was concentrated with Amicon centrifugal filters (Millipore) and further purified by size-exclusion chromatography in HBS over a Superose6 column (GE Healthcare). Peak fractions were collected and their quality ensured by SDS-PAGE followed by silver staining as per manufacturer specifications (Invitrogen) or Coomassie blue staining and negative-stain electron microscopy as described previously(*40*) (Fig. S4B-D).

### Cryo-EM specimen preparation and data collection

For the TCR–CD3 in GDN, Superose6 peak fractions were concentrated with Amicon centrifugal concentrators to 5 mg/mL as determined with a NanoDrop spectrophotometer (Thermo Fisher Scientific) assuming 1 AU = 1 mg/mL. Aliquots of 4 μL were applied to glow-discharged 400 mesh R1.2/1.3 Cu grids with graphene oxide (Quantifoil) using a Vitrobot Mark VI (Thermo Fisher Scientific) set at 4°C and 100% humidity. After 20 s, grids were blotted for 0.5 s with a blot force of −2 and plunged into liquid nitrogen-cooled ethane.

For the TCR–CD3 in nanodiscs, Superose6 peak fractions were concentrated with Amicon centrifugal concentrators to 330 mg/mL as determined with a NanoDrop spectrophotometer assuming 1 AU = 1 mg/mL. Aliquots of 4 μL were applied to glow-discharged 400 mesh R1.2/1.3 Cu grids using a Vitrobot Mark VI set at 4°C and 100% humidity. After 5 s, grids were blotted for 4 s with a blot force of −2 and plunged into liquid nitrogen-cooled ethane. For the TCR–CD3 in nanodiscs with HLA–β2M–NY-ESO-1, the Superose6 peak fractions were concentrated with Amicon centrifugal concentrators to 600 μg/mL. Lyophilized HLA single-chain trimer (herein, “HLA”; fusion of NY-ESO-1 C9V [SLLMWITQV], β2M I21-M119, and HLA-A*02:01 G25-T305; Kactus Biologics) was dissolved in deionized water per manufacturer instructions and then re-purified by size-exclusion chromatography over a Superose6 column into HBS to remove PBS and trehalose and concentrated to 1 mg/mL. The TCR–CD3 complex and HLA single-chain trimer mixed at a 2:1 volume ratio (1:5 approximate molar ratio). Aliquots of 4 μL were applied to glow-discharged graphene oxide-coated 400 mesh R1.2/1.3 Cu grids (Quantifoil) using a Vitrobot Mark VI (Thermo Fisher Scientific) set at 4°C and 100% humidity. After a wait time of 20 s, grids were blotted for 0.5 s with a blot force of −2 and plunged into liquid nitrogen-cooled ethane.

Cryo-EM imaging was performed in the Cryo-EM Resource Center at the Rockefeller University using SerialEM(*41*). Data collection parameters are summarized in Table S1. In brief, for all samples, data were collected on a 300-kV Titan Krios electron microscope at a nominal magnification of 64,000×, corresponding to a calibrated pixel size of 1.08 Å on the specimen level. For the TCR–CD3 in detergent sample, images were collected using a defocus range of −0.8 to −2.5 μm with a K3 direct electron detector (Gatan) in super-resolution counting mode. The ‘superfast mode’ in SerialEM was used, in which 3 × 3 holes are exposed using beam tilt and image shift before moving the stage to the next position(*42*). Exposures of 2.0 s were dose-fractionated into 50 frames (40 ms per frame) with a dose rate of 30 electrons per pixel per s (approximately 1.03 electrons per Å^2^ per frame), resulting in a total dose of 51 electrons per Å^2^.

For the TCR–CD3 in nanodisc sample, images were collected using a defocus range of −0.8 to −2.0 μm with a K3 direct electron detector (Gatan) in super-resolution counting mode with the ‘superfast mode’ in SerialEM. Exposures of 2.2 s were dose-fractionated into 55 frames (40 ms per frame) with a dose rate of 30 electrons per pixel per s (approximately 1.03 electrons per Å^2^ per frame), resulting in a total dose of 57 electrons per Å^2^.

For the TCR–CD3 in nanodiscs with HLA sample, images were collected using a defocus range of −0.8 to −2.0 μm with a K3 direct electron detector (Gatan) in super-resolution counting mode. The ‘superfast mode’ in SerialEM was used, in which 3 × 3 holes are exposed using beam tilt and image shift before moving the stage to the next position. Exposures of 2.0 s were dose-fractionated into 50 frames (40 ms per frame) with a dose rate of 30 electrons per pixel per s (approximately 1.03 electrons per Å2 per frame), resulting in a total dose of 51 electrons per Å^2^.

### Cryo-EM image processing for the TCR–CD3 in detergent

The 15,726 collected movie stacks were gain-normalized, motion-corrected, dose-weighted and binned over 2 × 2 pixels in Motioncorr2(*43*) and imported into cryoSPARC v4.2 for all further processing(*44*) (see data processing overview and motion-corrected micrograph examples in Fig. S2). A subset of the larger dataset, comprising 4,278 micrographs, was used to generate an initial model. The contrast transfer function (CTF) parameters were determined with CTFFIND4(*45*) implemented in cryoSPARC. Micrographs were curated by average defocus value, CTF fit resolution and CTF cross correlation, yielding 3,292 motion-corrected micrographs with CTF estimations for downstream processing.

Particles were picked with the cryoSPARC blob-picker using a size range of 80 to 180 Å. These initial particles were filtered by power level and normalized cross-correlation score. Particles were extracted with 2×2 binning and subjected to four rounds of 2D classification. Each round, particles with the best alignment, as determined quantitatively by resolution estimate or qualitatively by appearance of fine structure, were advanced to the next round of 2D classification and selection. This iterative 2D classification procedure yielded 133k particles for further analysis. These particles were then re-extracted without binning in 300×300-pixel boxes, yielding 131k particles. One additional round of 2D classification was performed. The best 82k particles were used for *ab initio modeling*, specifying three models. One model was clearly recognizable as TCR–CD3 in a detergent micelle; the other two models were amorphous blobs. All three models were used to seed 3D heterogeneous refinement using the same 82k particles, and the resulting TCR–CD3 model was used as an initial model in downstream processing.

We returned to the full dataset of 15,726 motion-corrected micrographs. CTF parameters were estimated using Patch CTF in cryoSPARC. Micrographs were curated by average defocus value, defocus range, CTF fit resolution, and relative ice thickness, yielding 15,153 micrographs for particle picking. Particles were picked with 20-Å low pass-filtered templates from the model of the TCR–CD3 in detergent determined above. These initial particles were filtered by power level and normalized cross-correlation score to yield 2.9M particles, of which 2.6M could be bounded into a 256×256-pixel box. Particles were extracted with 2×2 binning and subjected to three rounds of 2D classification. Well-aligning 2D classes were scarce in this particle stack, even after three rounds of 2D classification, and resulting 3D reconstructions were of low resolution (over 6 Å). For this and all nominal resolutions listed herein, resolutions are reported from gold-standard Fourier shell correlation (FSC) curves with a cut-off criterion of 0.143.

We endeavored to improve the quality of the picked particles using the Topaz machine learning suite embedded in cryoSPARC. The best 36k particles, coming from 4 distinct 2D classes in the final round of classification, were used to train a Topaz particle picking model in cryoSPARC. The Topaz model was trained over 10 epochs, with an expected particle number of 100 per micrograph using 400 micrographs chosen at random from the 8,125 micrographs with the highest median pick cross-correlation value from template picking. The model was then used to pick particles from the 8,125 micrograph set, yielding 615k particles, of which 595k could be bounded in a 256×256-pixel box. Particles were subjected to two rounds of serial 2D classification, and the best 342k particles were advanced to serial 3D multi-reference (“heterogeneous”) refinement. Heterogeneous refinement was seeded with the initial model from the exploratory micrograph subset, along with three decoy noise models to remove poorly aligning particles. Decoy classes were created by aborting *ab initio* modeling jobs after the first round of modeling before any structure was visible. After each round of 3D heterogenous refinement, particles from the classes with the highest resolution were advanced to the next round and their respective maps (or, in later rounds, the best available maps) used to seed the refinement job in competition with decoy maps. After two rounds of iterative 3D classification against noise decoys, the best 94k particles were used for non-uniform refinement seeded with the TCR–CD3 model from that same round of heterogenous refinement. This map refined to a nominal resolution of 4.7 Å.

To improve map quality, we performed multiple rounds of Topaz model training and iterative 2D and 3D classifications as above. To start, we retrained the first Topaz model with an expected particle count of 400 per micrograph. The number of particles picked increased to 1.3M, of which 1.2M could be bounded by a 256×256-pixel box and were extracted with 2×2 binning. These particles were subjected to two rounds of serial 2D classification, yielding 977k particles, which were in turn subjected to four rounds of serial 3D classification seeded with the best prior model against 3 noise models each, yielding 145k particles for non-uniform refinement. Map quality was marginally improved to a nominal resolution of 4.6 Å. To identify latent heterogeneity within the particle stack, cryoSPARC 3D variability analysis was performed on three eigenvectors, and reconstructions from opposing ends of the first eigenvector were used to seed a single round of heterogeneous refinement. The better resolving class, comprised of 85k particles, was subjected to non-uniform refinement, and the resulting map improved to a nominal resolution of 4.5 Å. These 85k particles were used to train a new Topaz model, again expecting 400 particles per micrograph, but number of epochs increased to 20. This model was used to pick particles from the entire 15,153 micrograph set, yielding 1.6M particles. These were extracted unbinned in a 256×256-pixel box. After two rounds of serial 2D classification, 1.5M particles remained, which were subjected to four rounds of 3D classification seeded initially with the two models from 3D variability analysis and three decoy noise classes. The 248k particles in the best class of the last round of heterogeneous refinement were subjected to non-uniform refinement, with now significant improvement in map quality (nominal resolution 3.4 Å). Re-extraction into a larger 320×320-pixel box resulted in an improvement in resolution to 3.3 Å, and these 242k particles were used to train a final Topaz model, with an expected particle number of 250 per micrograph and 30 epochs of training. Picking from the 15,153-micrograph set with this model yielded 2M particles, which were extracted in 256×256-pixel boxes with 2×2 binning. These particles were subjected to two rounds of serial 2D classification, and the best 1.2M particles were extracted unbinned in 256×256-pixel boxes. These particles were subjected to one round of 2D classification and the best 951k particles were entered into three rounds of serial 3D classification seeded with the best prior model and 3 noise decoys. This yielded 311k particles which were subjected to an additional round of 2D classification, from which 248k particles were selected. These particles, along with those from the best prior Topaz-picked model, were re-extracted in 300×300-pixel boxes and pooled to ensure that all high-quality particles were captured for final reconstruction. One round of 2D classification was performed on the extracted particles and duplicates removed, yielding 388k unique particles. Of these, the 310k best-aligning particles were selected to advance into a final cascade of five rounds of serial 3D classification seeded with the best prior model and 3 noise decoys. The final 252k particle set emerging from these classifications was subjected to non-uniform refinement, yielding a map with nominal resolution of 3.3 Å. Local refinement with a mask including the protein components but excluding the nanodisc density further improved map quality and interpretability.

### Cryo-EM image processing for the TCR–CD3 in nanodiscs

The 17,766 collected movie stacks were gain-normalized, motion-corrected, dose-weighted and binned over 2 × 2 pixels in Motioncorr2(*43*) and imported into cryoSPARC v4.2 for all further processing(*44*) (see initial processing streams and motion-corrected micrograph examples in Fig. S5). The contrast transfer function (CTF) parameters were determined with CTFFIND4(*45*) implemented in cryoSPARC. Micrographs were curated by CTF fit resolution and cross correlation, yielding 14,852 motion-corrected micrographs with CTF estimations for downstream processing.

Three parallel processing streams were pursued based on different approaches to particle picking (Fig. S5). Particles were picked in cryoSPARC using (1) an unbiased blob-picking approach, (2) a template-based approach with a model of the TCR–CD3 in detergent (PBD 6JXR(*5*)) docked into the MSP 1D1DH5 nanodisc using the CHARMM-GUI(*46*) nanodisc builder(*47*), and (3) a template-based approach using the first published structure of the TCR– CD3 in detergent (PBD 6JXR(*5*)). For each processing stream, 2D image classification was performed on 4×4 binned particle images to eliminate classes of noise or poorly aligning particles and *ab initio* modeling was performed. At this stage, it was clear that the particles picked using 6JXR templates were of poor quality and did not yield reconstructions substantially different from those obtained with the other two processing streams (see below) and this avenue was thus aborted.

Data were processed in the blob-picking stream as follows: Particles were picked using a particle size range of 60 to 120 Å, and initial particles were filtered by power level and normalized cross-correlation. Particles were extracted with 4×4 binning and subjected to 4 rounds of 2D classification. Each round, particles with the best alignment, as determined quantitatively by resolution estimate or qualitatively by appearance of fine structure, were advanced to the next round of 2D classification and selection. This iterative 2D classification procedure yielded 6.3M particles for further analysis. *Ab initio* modeling was performed with this particle set specifying four models. Of these four initial models, two were recognizable as protein in nanodiscs, and these models were used to seed 3D heterogenous refinement.

Additionally, two decoy noise models were added as seeds for 3D heterogenous refinement to remove poorly aligning particles; these classes were created by aborting *ab initio* modeling jobs after the first round of modeling before any structure was visible. After each round of 3D heterogenous refinement, particles from the classes with the highest resolution were advanced to the next round and their respective maps used to seed the refinement job in competition with decoy maps. After two rounds of 3D heterogenous refinement with two iteratively improving classes and two decoy classes, the number of particles remaining was 2.6M. To exhaustively capture all well-aligning particles, the same iterative process was performed on the 6.3M particle set with a single *ab initio* model and two rounds of heterogeneous refinement, yielding an additional 595k unique particles that were pooled with the 2.6M particle set and re-extracted with a 256×256-pixel box but only 2×2 binning. These 3.1M particles were subjected to one round of 2D classification and selection, yielding 3M particles. Of these, the best aligning 670k particles were used for a new round *ab initio* modeling specifying 4 classes to attempt to expand the diversity of conformations, seeding further 3D heterogeneous refinement and selection. These four models were refined by one round of heterogeneous refinement and one round of non-uniform refinement and then used (along with two decoy maps) to seed a round of heterogeneous refinement with the 3M 128×128-pixel particles. A single class with 883k particles refining to a nominal resolution of 4.5 Å was selected for re-extraction in a 256×256-pixel box without binning. One round of heterogenous refinement with two copies of the advancing maps and two decoy maps was performed, yielding two classes refining to higher resolution (3.4 and 3.7 Å; 420k and 394k particles, respectively). To further subclassify these two particle sets, the pooled 814k particles were subjected to 3D variability analysis(*48*) with four eigenvectors. To avoid classification against rotational and translational variation in nanodisc position relative to the TCR–CD3, a custom mask was employed to remove nanodisc density. Four maps were generated using clustering of the particles in 3D variability latent space. Three of these maps with distinct spatial features were used to seed a 3D heterogenous refinement against three decoy maps. Two rounds of 3D heterogeneous refinement were performed, and the particles in the three iteratively improving classes were then re-extracted in 320×320 pixel boxes and subjected to non-uniform refinement(*49*) and DeepEMhancer sharpening(*50*) as shown in Fig. S5, bottom left (181k particles, 3.7 Å resolution; 228k particles, 3.4 Å resolution; 212k particles, 3.7 Å resolution). These particles were pooled with the output from the 6JXR-in-nanodisc templated stream, as below.

Data were processed in the 6JXR-in-nanodisc template-picking stream as follows: Particles were picked with a 20-Å low pass-filtered model of the TCR–CD3 in a nanodisc (see above). These initial particles were filtered by power level and normalized cross-correlation then extracted with 4×4 binning and subjected to two rounds of 2D classification and selection as described above. Iterative 2D classification yielded 7.2M particles for further analysis. *Ab initio* modeling was performed with this particle set specifying four initial models. Of these four initial models, two were recognizable as protein in nanodiscs, and these models were used to seed 3D heterogenous refinement against decoy maps, as described above. After two rounds of 3D heterogenous refinement with two iteratively improving classes and two decoy classes, the number of particles remaining was 2.9M. To exhaustively capture all well-aligning particles, the same iterative process was performed on the 7.2M particle set with two *ab initio* models and two rounds of heterogeneous refinement, yielding an additional 612k unique particles that were pooled with the 2.9M particle set and re-extracted with a 256×256-pixel box but only 2×2 binning. These 3.4M particles were subjected to one round of 2D classification and selection, yielding 3.1M particles. Of these, the best aligning 605k particles were used for a new round of *ab initio* modeling specifying two classes. Only one class resembled protein-in-nanodisc, and it was used to seed four rounds of iterative heterogeneous refinement against one decoy map. This reduced the number of particles from 3.1M to 1.2M. To expand the diversity of conformations seeding further 3D heterogeneous refinement and selection, the 605k particles used previously for *ab initio* modeling of two classes were subjected to *ab initio* modeling specifying four classes. The resulting four models were refined by one round of heterogeneous refinement and one round of non-uniform refinement and then used (along with two decoy maps) to seed a round of heterogeneous refinement with the 3.1M 128×128-pixel particles. The two classes yielding the highest resolution maps comprised of 1.2M particles and the maps were used to seed another round of 3D heterogeneous refinement against two decoy maps, yielding 1.1M particles in the two highest resolution classes. Pooling the particles from these two parallel approaches (starting with *ab initio* modeling of two and four classes from 2×2-binned particles) yielded 1.7M unique particles in three classes, which were advanced for re-extraction in a 256×256-pixel box without binning. To further subclassify this particle set, the pooled 1.7M particles were subjected to 3D variability analysis(*48*) with four eigenvectors. The third eigenvector showed the most concerted variation. The particles were segmented into six classes along this latent space dimension and the respective reconstructions used to seed one round of 3D heterogeneous refinement. The two highest-resolution classes, comprised of 710k particles, were subjected to 3D variability analysis(*48*) with three eigenvectors. Four maps were generated using clustering of the particles in 3D variability latent space and were used to seed one round of 3D heterogenous refinement. The particles in the two highest-resolution classes were re-extracted in 320×320-pixel boxes and subjected to non-uniform refinement and DeepEMhancer sharpening as shown in Fig. S5, bottom center (198k particles, 3.8 Å resolution; 203k particles, 3.5 Å resolution).

To further improve map quality and resolution, the 902k unique particles from the two aforementioned processing streams were pooled and subjected to one round of 3D heterogeneous refinement using the five input maps. This allowed the identification of three maps denoted ND1, ND2, and ND3, with nominal resolutions of 3.5, 3.5, and 3.3Å, respectively (Fig. S6).

Given the map quality improvements seen with in iterative Topaz picking models in processing the TCR–CD3-in-GDN dataset, we employed a similar approach here (Fig. S6). Additionally, initial CTF estimates were recalculated for all 17,766 micrographs using the cryoSPARC Patch CTF utility. The particles used to reconstruct ND1-3 were used to train a Topaz model over 30 epochs on a subset of 938 micrographs with an expectation of 250 particles per micrograph. This yielded 4.5M particles picks that were curated by micrograph CTF resolution estimate, average defocus, and pick power score to give 4M particles, which were subsequently extracted in 256×256-pixel boxes with 2×2 binning. After three rounds of serial 2D classification, selection, and re-extraction into 256×256-pixel boxes without binning, 954k particles remained. These particles were used to train a new Topaz picking model over 100 epochs on a subset of 896 micrographs with an expectation of 250 particles per micrograph, yielding 4.6M particles extracted into 256×256-pixel boxes with 2×2 binning. After one round of 2D classification and selection, 4.4M particles remained, which were subjected to one round of 3D heterogeneous refinement seeded with the ND1-3 maps and two noise classes. The 3.3M particles aligning to the ND1-3 starting models were re-extracted without binning and subjected to two rounds of 2D classification and selection, yielding 1.3M particles.

To further expand the pool of particles available for 3D reconstruction, the ND1-3 maps were used to generate templates for particle picking. 31M particles were curated by power score and normalized cross correlation, followed by extraction into 256×256-pixel boxes with 4×4 binning to yield 11.6M particles. After three rounds of serial 2D classification 4.5M particles remained, which were re-extracted into the same size boxes, but with 2×2 binning. These particles were subjected to three rounds of serial 3D heterogeneous refinement seeded with the ND1-3 maps and three noise classes, after which 1.6M particles remained. These particles were re-extracted in the same box size without binning and subjected to one more round of 3D heterogeneous refinement as above, and the resulting 1.5M particles were used to train a Topaz model in 20 epochs on a subset of 999 micrographs, with an expectation of 250 particles per micrograph. The resulting 4.6M particles were curated by micrograph CTF fit resolution estimation and particle power score, then extracted into a 256×256-pixel box with 2×2 binning to yield 4M particles. One round of 2D classification and selection yielded 3.7M particles. These particles were separated into three groups by an initial round of 3D heterogeneous refinement seeded with the ND1-3 models. For each of these three parallel streams, the particles were subjected to three rounds of 3D heterogeneous refinement for each model against three noise classes, and the resulting particles for the three streams pooled. These 1.4M particles were re-extracted into a 300×300-pixel box without binning and subjected to a single round of 3D heterogeneous refinement against the highest resolution maps from the preceding three parallel streams. The intermediate (ND2-like) class had the highest resolution, and its 883k particles were kept for further refinement.

Having now exhaustively sampled the micrographs for well-aligning particles, the best particles from the several approaches above were pooled: the original ND1-3 sets (638k particles), the particles originating from the two Topaz models trained on this set (954k and 1.3M particles), and the particles originating from ND1-3 template picking and subsequent Topaz-guided picking (883k particles). After removal of duplicate particles, 2.2M unique particles were extracted into 256×256-pixel boxes without binning. After one round of 2D classification and selection, 1.8M particles remained. At this point, we hypothesized that the ND2 conformation likely represented a transitional state on a continuum between the more distinct ND1 and ND3 conformations, with the ND2 maps showing some averaging of features from both ND1 and ND3 maps. Thus, these 1.8M particles were subjected to two round of serial 3D heterogenous refinement seeded with the ND1 and ND3 states along with two noise classes, yielding 630k and 809k particles respectively. After re-extraction into 300×300-pixel boxes without binning, the two classes contained 620k and 797k particles, respectively, and refined to nominal resolutions of 3.2 Å and 3.0 Å, respectively. In parallel, we performed the same serial four-class 3D heterogeneous refinement with ND1, ND2, ND3, and a noise class as seeds. The maps from the ND1/ND3-seeded refinements were of higher quality and nominal resolution than the ND1/ND2/ND3-seeded refinements (Fig. S6), supporting the hypothesis that the ND1 and ND3 states represent more highly populated and stable conformations than the transitional ND2. We thus renamed ND1 and ND3 as ND-I and ND-II, respectively, and used them as the final maps for further analysis.

### Cryo-EM image processing for the TCR–CD3 in nanodiscs incubated with HLA

The 13,644 collected movie stacks were gain-normalized, motion-corrected, dose-weighted and binned over 2 × 2 pixels in Motioncorr2(*43*) and imported into cryoSPARC v4.2 for all further processing(*44*) (see processing scheme and motion-corrected micrograph examples in Fig. S15). The contrast transfer function (CTF) parameters were determined with CTFFIND4(*45*) implemented in cryoSPARC. Micrographs were curated by average defocus, astigmatism, CTF fit resolution, and CTF fit cross correlation, yielding 10,610 motion-corrected micrographs with CTF estimations for downstream processing.

Particles were initially picked by a template-based approach using a model of the TCR– CD3 in nanodiscs ND-II conformation with HLA from PDB 2BNQ(*51*) docked by alignment of the TCR variable domains. Templates were 20-Å low pass-filtered prior to particle picking.

These initial particles were filtered by power level and normalized cross-correlation score then extracted with 4×4 binning and subjected to five rounds of 2D classification and selection as described above. From iterative 2D classification, the best 324K particles were selected re-extraction in 288×288-pixel boxes without binning. After one round of 2D classification, the best-aligning 84k particles were used to train a Topaz picking model (500 micrographs, 200 particles expected per micrograph, 50 training epochs). For Topaz picking, a 10,761-micrograph set was used, which had been curated as above for the 10,610 micrograph set, but with slightly liberalized cutoffs. After picking particles with the Topaz model, micrographs were further curated by the number of particles picked per micrograph and CTF fit cross correlation, yielding 1.6M particles from 7,003 micrographs. These particles were extracted into 288×288-pixel boxes with 2×2 binning. After one round of 2D classification, the best-aligning 106k particles were used to generate a single *ab initio* model. This model was refined with non-uniform refinement and then used to seed 3D heterogenous refinement against two decoy maps, as described above. After one round of 3D heterogenous refinement 82k particles remained. These particles were used to train a new Topaz model (500 micrographs, 400 particles expected per micrograph, 50 training epochs). This model was used to pick 2.8M particles, of which 2.7M particles could be extracted into 288×288-pixel boxes, without binning. After one round of 2D classification, 2.6M particles were advanced to 3D heterogeneous refinement seeded with the prior best map and two decoy classes, from which 1.3M particles aligned to the target map and advanced. These 1.3M particles were subjected to 3D classification without alignment, specifying six target classes seeded by principal component analysis of 100 reconstructions (1000 random particles per reconstruction). All 1.3M particles were then subjected to 3D heterogeneous refinement, seeded with the six volumes output by the prior 3D classification job. One class comprising 310k particles refined to substantially higher resolution than the others (3.2 Å after subsequent non-uniform refinement and re-extraction into 300×300-pixel boxes). These particles were used to train a new Topaz model (500 micrographs, 200 particles expected per micrograph, 50 training epochs). 2.4M particles were picked with this new model, and extracted into 288×288-pixel boxes with 4×4 binning. After one round of 2D classification, the best-aligning 2.3M particles were re-extracted into 288×288-pixel boxes without binning. One round of 2D classification reduced the number of best-aligning particles to 2.2M, and these particles were re-extracted unbinned in 320×320-pixel boxes. At this point, 2D classes showed clear density for a TCR–CD3 complex in a nanodisc with additional blurry density for the HLA emanating from the apex of the TCR variable domains. The 165k particles with the best HLA density were selected for 3D classification without alignment using a custom focus mask inclusive of the TCR variable domains and HLA. 3D classification was seeded by principal component analysis, as above, and two classes were specified. The two resulting volumes showed strong vs. weak density for the bound HLA. 3D heterogenous refinement was performed with the 165k particle set and seeded with the two output volumes from 3D classification without alignment. 77k particles segregated into the class with stronger HLA density. To further enrich for particles with strong HLA density, these 77k particles were subjected to 3D heterogeneous refinement, again seeded with the output volumes from 3D classification without alignment. The resulting 58k particles were used to seed a new round of Topaz picking (500 micrographs, 100 particles expected per micrograph, 50 training epochs), which identified 1.8M particles. To ensure exhaustive sampling of the available data, these particle picks were combined with those from the previous two rounds of Topaz-aided particle picking, and duplicate particles were removed, yielding 3.3M unique particles. These particles were extracted in 400×400-pixel boxes (binned 2×2) to ensure adequate accommodation for delocalized signal around the elongated TCR–CD3–HLA particles. One round of 3D heterogenous refinement was performed, seeded with the HLA-strong and HLA-weak outputs from the prior 3D classification and 2 decoy classes. The 895k particles aligning to the HLA-strong model were subjected to an additional cycle of 3D classification without alignment, seeded by principal component analysis and specifying six classes. The six output volumes were used as the starting classes for one round of heterogeneous refinement with then 895k-particle set. One class comprising 192k particles refined to higher resolution than the others. The same analysis (3D classification without alignment followed by 3D heterogenous refinement) was performed for the HLA-weak set, yielding a single high-resolution class with 215k particles. These two sets of particles were re-extracted into a 400×400-pixel box without binning, and combined with the prior best particle stack, comprising 297k particles, also extracted now in a 400×400-pixel box. Duplicate particles were removed, yielding 587k unique particles. These particles were subjected to four rounds of serial 3D heterogeneous refinement seeded with the HLA-strong and HLA-weak models, and the particles classifying in the HLA-strong class were advanced to the next round. This yielded 155k particles in the final set. Non-uniform refinement of this particle set yielded a map with a nominal resolution of 3.6 Å. Local refinement was performed with a custom mask excluding the bulk lipid of the nanodisc, yielding a map with a nominal resolution of 3.5 Å. This map was used for refinement in Phenix. Additional local refinements on subsets of the structure (e.g., TCR variable domains–HLA) were performed and used to guide manual model building but were not used for final refinement or calculation of refinement statistics.

### Cryo-EM map assessment and post-processing

Gold-standard FSC curves were generated in cryoSPARC during refinement and nominal resolutions are reported with a cut-off criterion of 0.143. FSC curves for final maps are shown in Fig. S3, S7, and S15. Map anisotropy was assessed with 3DFSC(*52*) implemented in cryoSPARC and local resolution analysis was also performed in cryoSPARC (Fig. S3, S7, S15). Although all models were refined against unsharpened maps (see *Model building and refinement*, below), sharpened maps were used for visual interpretation, manual model building, and figure generation. Map sharpening (Fig. S8) was performed either by global B-factor approaches (better visualization of lipid densities) or DeepEMhancer(*50*) (better visualization of protein densities).

### Model building and refinement

The model of TCR–CD3 in GDN micelles was built starting with rigid-body docking of prior models in ChimeraX(*53*). Specifically, PDB 6JXR was used as a starting model for the CD3 proteins and the TCR constant regions and transmembrane helices, PDB 7FJD was used to dock the cholesterols, and PDB 6BNU(*51*) was used as a starting model for the 1G4 TCR variable regions. This starting model readily docked into the map, anchored by the clear alignment of immunoglobulin folds to the map features and continuous density for all chains through the ectodomains, juxtamembrane linkers, and transmembrane helices (see example densities in Fig. S9A). As with prior TCR–CD3 structures, the cytoplasmic unstructured regions were not resolved in the map. The model was improved by iterative cycles of refinement with phenix.real_space_refine(*54*), manual adjustment in Coot(*55*), and brief map-constrained relaxation in ISOLDE(*56*). Residues without clear side chain density beyond Cβ were manually modeled as stubs. Cores for the asparagine-linked glycans were built in Coot using tools included in the package(*57*) (modeled glycans indicated graphically in Fig. S14D). For all models, refinement data are shown in Table S2, and model completeness is shown in Table S3. All cryo-EM map and molecular model figures were generated in ChimeraX.

For the ND-I and ND-II models, the TCR–CD3-in-GDN model built above was used as a starting model and domains were docked into the map in ChimeraX. Again, the immunoglobulin folds in the TCR and CD3 ectodomains were readily anchored into the map features, with bulky hydrophobic residues and disulfide bonds periodically confirming register (map density overviews and sections shown in Fig. S9). Regions without clear density in the maps were removed and residues without clear side chain density beyond Cβ were manually modeled as stubs. For the CD3, CD3γ, and CD3δ proteins, the register for the juxtamembrane linker and transmembrane helices were anchored by the placement of the C-X-X-C lariats at the beginning of the juxtamembrane region (Fig. S9D). Bulky hydrophobic sidechains provided further confirmation of register within the transmembrane helices. In ND-I, the register at the amino-terminus of the CD3 _G_ transmembrane helix could not be unambiguously determined, so this helix was left unmodeled. For CD3ζ, the GDN model of these chains was docked in place and manually refit, using the interchain disulfide bond and bulky hydrophobic residues to confirm register (Fig. S9E). For TCRα and TCRβ, the linker peptides between their constant domains and the transmembrane helices were poorly structured and left unmodeled. Their transmembrane helices were docked from the GDN structure relative to the surrounding helices and bulky side chains were used to confirm register (Fig. S9F). Model building and refinement was performed in Phenix, Coot, ChimeraX, and ISOLDE as described above for the TCR–CD3-in-GDN structure.

For the TCR–CD3–HLA structure, the GDN structure readily docked into the cryo-EM map. The HLA–β2M–NY-ESO-1 structure from PDB 2BNQ(*51*) was docked into the map. The structure was refined in Phenix as described above. Example density with constructed model is shown in Fig. S16.

### Sequence conservation analysis

TCR and CD79-BCR sequences were analyzed using the Consurf(*58*) server and colored according to relative sequence conservation level in ChimeraX. For computation of sequence identity among GDN and the published TCR–CD3 structures in detergent, sequences were aligned with Clustal Omega.

### Engineered disulfide bond analysis

Cysteine mutant pairs were designed with the Cα-Cα parameters described in the main text. For Western blot analysis, HEK293F cells were plated at 1.5e6 cells/mL in 6-well plates, 2 mL per well. Cells were infected with baculovirus expressing the indicated TCR–CD3 mutants at a volume 1/30 the culture volume and the indicated CD3 mutants at 1/15 the culture volume. All TCRα mutants were paired with the same wild-type TCRβ sequence. 10 mM sodium butyrate was added at the same time as viral infection to enhance expression. Cultures were incubated for 2 days at 37°C, 8% CO_2_. Cells were harvested by centrifugation at 20,800xg, and resuspended in Laemmli sample buffer with or without 5% 2-mercaptoethanol (Sigma Aldrich). Samples were heated for 5 min at 95°C prior to separation by SDS-PAGE. Gels were electroblotted to immobilon-P PVDF membranes (Millipore), rinsed in 200 mM NaCl, 20 mM Tris (Research Products International), pH 8 (TBS), supplemented with 0.05% Tween-20 (TBST, Sigma Aldrich)), and blocked in neat Intercept Blocking Buffer (“block,” Li-Cor) for 1 h at room temperature. Blots were incubated with mouse anti-GFP (1:3,000, Roche/Sigma Aldrich) in TBST with 10% block overnight at 4°C. Blots were washed with TBST at 25°C, incubated with Alexa 700-conjugated goat anti-mouse IgG (1:10,000, Invitrogen) in TBST with 10% block at 25°C for 1 h, and again washed sequentially with TBST and TBS. Blots were imaged on an Amersham Typhoon infrared fluorescence scanner. To confirm the results seen in whole cell lysates, mutant TCR–CD3 was also detergent extracted from membrane pellets and NiNTA-affinity purified, as was done for cryo-EM analyses, and NiNTA elutions in DDM/CHS were processed for Western blot. To further confirm that these crosslinks were formed in properly assembled TCR–CD3 complexes, select disulfide mutants were also reconstituted into nanodiscs and purified by size-exclusion chromatography, by the same protocol used for producing TCR–CD3 in nanodiscs for cryo-EM analysis. For these samples, protein constituents were visualized in gels using Acquastain as per manufacturer specifications. For mass spectrometry readout, protein was produced as for structural studies but using the indicated mutant TCR–CD3 constructs with TCRβ fused to EGFP and CD3ζ untagged. The CD3 proteins, including 8xHis-tagged CD3γ were expressed from a single virus. Instead of reconstitution into nanodiscs, the TCR–CD3 complex was eluted from NiNTA agarose resin in HBS containing DDM/CHS, free cysteines were blocked with 20 mM iodoacetamide (Sigma Aldrich) in the dark for 1 h at room temperature. The reaction was quenched and detergent removed by acetone precipitation.

### Mass spectrometry

Protein complexes were deglycosylated with PNGase F and sequentially digested using sequencing-grade trypsin for 3 h, followed by a second digestion step using chymotrypsin for another 3 h. Enzymatic activity was quenched by acidification using trifluoroacetic acid and the resulting peptides were purified using in-house constructed C18 micropurification tips prior to LC-MSMS analysis(*59*). Peptides were separated across a 70-min linear gradient, in which the fraction of solvent B (80% acetonitrile, 0.1% formic acid) over solvent A (1% acetonitrile, 0.1% formic acid) was increased from 2% to 35% using an Easy nLC 1200 Ultra-High Performance Liquid Chromatography system (Thermo Fisher Scientific) equipped with a 100 μm-by-120 μm pulled-emitter column packed with 3-μm C18 material (Nikkyo Technos). Analysis was carried out using a Fusion Lumos Tribrid mass spectrometer (Thermo Fisher Scientific) operating in positive-ion DDA, FT-mode. Ions were fragmented using EThcD. Disulfide bonds were identified using the XlinkX software through Proteome Discoverer v. 2.5 (Thermo Fisher Scientific).

### Viral transduction of T-cell lines

SFG-1G4, and SFG-1G4 S104C/V182C plasmids were synthesized by Genscript. For transfection of the plasmids mentioned above, 293GP cells (Takara Bio, 631458) were co-transfected with 4.5 μg SFG plasmid and 2.25 μg RD114 feline envelop plasmid (provided by S. A. Rosenburg, NIH) using Lipofectamine™ 3000 Transfection Reagent (Thermo Fisher Scientific, L3000075). Viral supernatant was harvested on day 2 post transfection and always used freshly.

T-cell line transduction was mediated by retronectin (Takara, T100B). In brief, retronectin was coated on a non-tissue-culture-treated plate at 10 μg/mL in PBS overnight. Coated plates were washed by PBS and blocked by 2% FBS in PBS for 30 min at room temperature. The blocking solution was aspirated, viral supernatant was added to the coated plates, and the plates were centrifuged at 2500xg for 2 h at 32°C. Viral supernatant was aspirated after centrifugation. Cells were loaded on plates and spun at 300g for 30 min at 32°C. On day 2 post transduction, transduction rates were measured by flow cytometry (see below) after staining with phycoerythrin-labeled anti-human TCR Vβ13.1 antibody (Biolegend, 362410). Cells were stained by iTAg tetramer/APC – HLA-A*02:01 NY-ESO-1 (SLLMWITQC) (MPL, TB-M011-2) and enriched using EasySep™ Human APC Positive Selection Kit II (Stemcell Technologies, 17661).

### Flow cytometry

For cell-surface staining, cells were stained with antibodies for 45 min in PBS + 0.5% FBS at 4°C. Cells were washed twice with PBS + 0.5% FBS and flow cytometry data were acquired on an LSRFortessa X-20 (BD). Antibodies used in our surface staining are Brilliant Violet 785™ anti-human CD3 antibody (Biolegend, 317330), and PE anti-human TCR Vβ13.1 antibody. To visualize HLA binding, iTAg tetramer/APC-HLA-A*02:01 NY-ESO-1 (SLLMWITQC) (MPL, TB-M011-2) was used. Cells were counterstained with the viability reagent LIVE/DEAD Fixable Aqua Dead Cell Stain Kit as per manufacturer’s specifications (Thermo Fisher Scientific, L34966).

### Generation of an NFAT-reporter cell line

The J8Zb2m^-^α^-^β^-^ cell line for TCR–CD3 display was generated as follows. The Jurkat-derived cell line J.RT3-T3.5 (ATCC TIB-153) lacking endogenous TCRβ chain was modified to express an NFAT reporter and CD8 via transduction with retroviral supernatant from 293Galv9 producer cells transfected with the 8xNFAT-ZsG-hCD8 plasmid (Addgene: #153417). 48 h post transduction, transductants were stimulated with PMA/Ionomycin overnight and sorted on CD8^+^ZsGreen^+^ cells using a Sony SH800. Sorted cells were allowed to recover *in vitro* for 5 days and subsequently nucleofected using a Nucleofector IIb device (Program X-001) with β2M sgRNA (5’-CGCGAGCACAGCUAAGGCCA-3’; Synthego) pre-complexed with TrueCut Cas9 V2 (Invitrogen, A26498) according to the manufacturer’s recommendation. Nucleofected cells were allowed to recover in RPMI + 2 mM L-glutamine + 20% FCS without antibiotics for 2 days and thereafter HLA-ABC^-^ live cells were sorted using a Sony SH800.

To then render the resulting cell line deficient for the TCRα chain, TRAC gene with sgRNA (5’-TCTCTCAGCTGGTACACGGC-3’; IDT) and Alt-R S.p. Cas9 Nuclease V3 (IDT 1081058) were introduced using a Lonza 4D-Nulceofector. TRAC knock-out was confirmed by ICE CRISPR Analysis Tool (Synthego) using Sanger sequencing data flanking the edited cut site. Primer pair: Forward 5’-AGCTTGTGCCTGTCCCTGAG-3’, Reverse 5’-GGGCTGGGGAAGAAGGTGT-3’. The sorted cell line, termed J8Zb2m^-^α^-^β^-^, was expanded and maintained *in vitro* in RPMI + 2 mM L-glutamine + 10% FCS + 100 μg/mL penicillin/streptomycin.

### TCR–CD3 activation assay with Incucyte system

J8Zb2m^-^α^-^β^-^cells with or without transduction were loaded onto 96 well plates (Corning, 353072) at 2e5 cells per 100 μL complete medium in triplicates. Plates were left at room temperature for 20 min and transferred to Incucyte SX1 system (Sartorius) for baseline images. The plate was then removed from Incucyte and loaded with 100 μL of iTAg tetramer/APC-HLA-A*02:01 NY-ESO-1 at 8 μg/mL, 4 μg/mL or 0 μg/mL as designated. Cell Stimulation Cocktail (PMA and ionomycin, Thermo Fisher Scientific, 00-4970-93) was added at 1:1000 as a positive control. Again, the plate was left at room temperature for 20 min and transferred to Incucyte system for imaging. Instrumental setting: 20x objective lens, 9 images per well, 2-h scan interval. Analysis was done using Incucyte Basic Analysis Software. P values were determined by unpaired, two-tailed t-tests using Prism 10 software.

## Other Supplementary Materials for this manuscript include the following

Movies S1 to S2

---

**Fig. S1:**
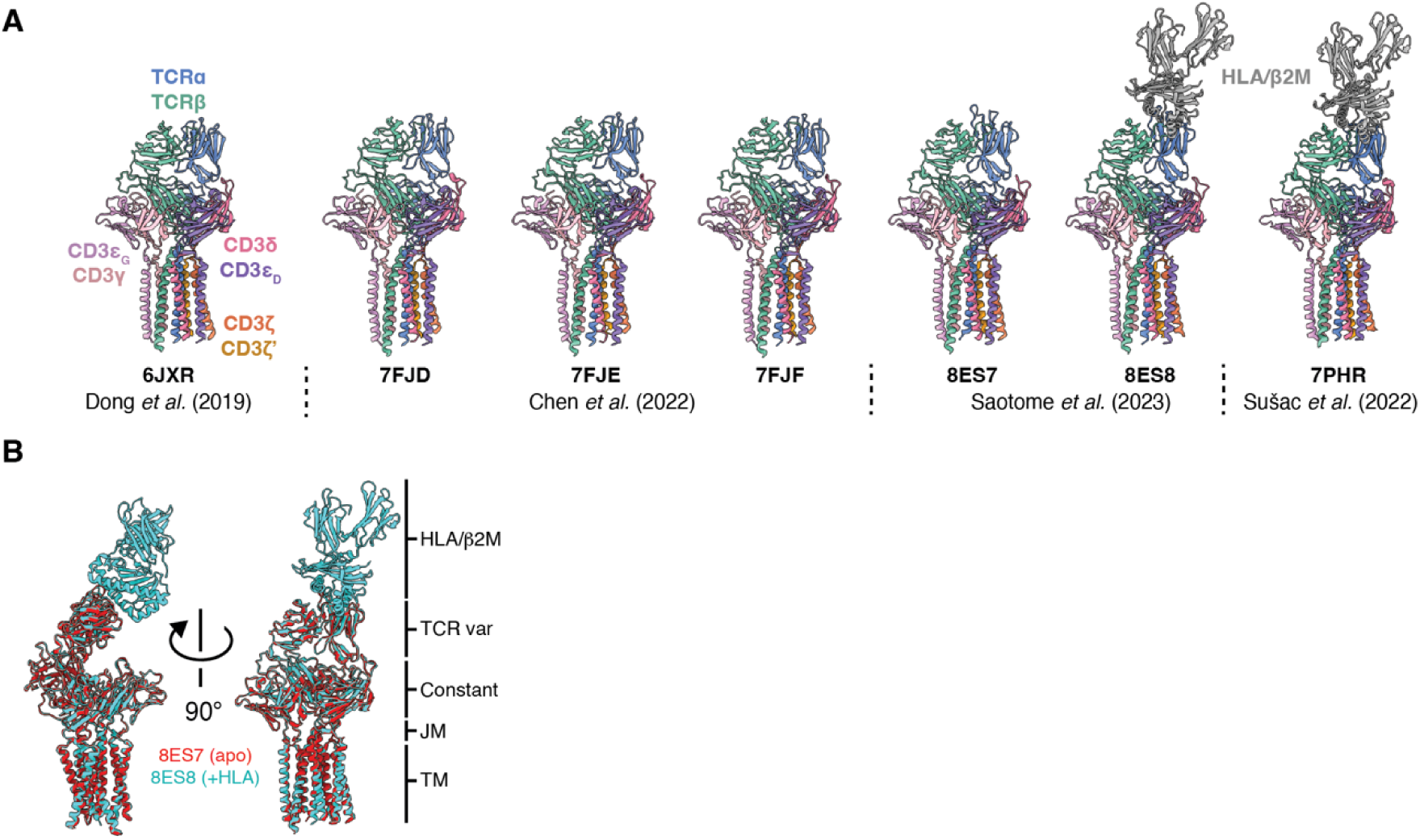
Previously published structures of the TCR–CD3 in detergent. (**A**) Cartoon representations of previously published structures of the TCR–CD3 by itself and in complex with HLA–β2M–antigen determined in detergent micelles by cryo-EM. Models are colored by chain as indicated. (**B**) Superposition of published structures of a TCR–CD3 by itself (red) or in complex with HLA–β2M–antigen (cyan), showing no meaningful structural change throughout the TCR structure upon ligand binding.

**Fig. S2:**
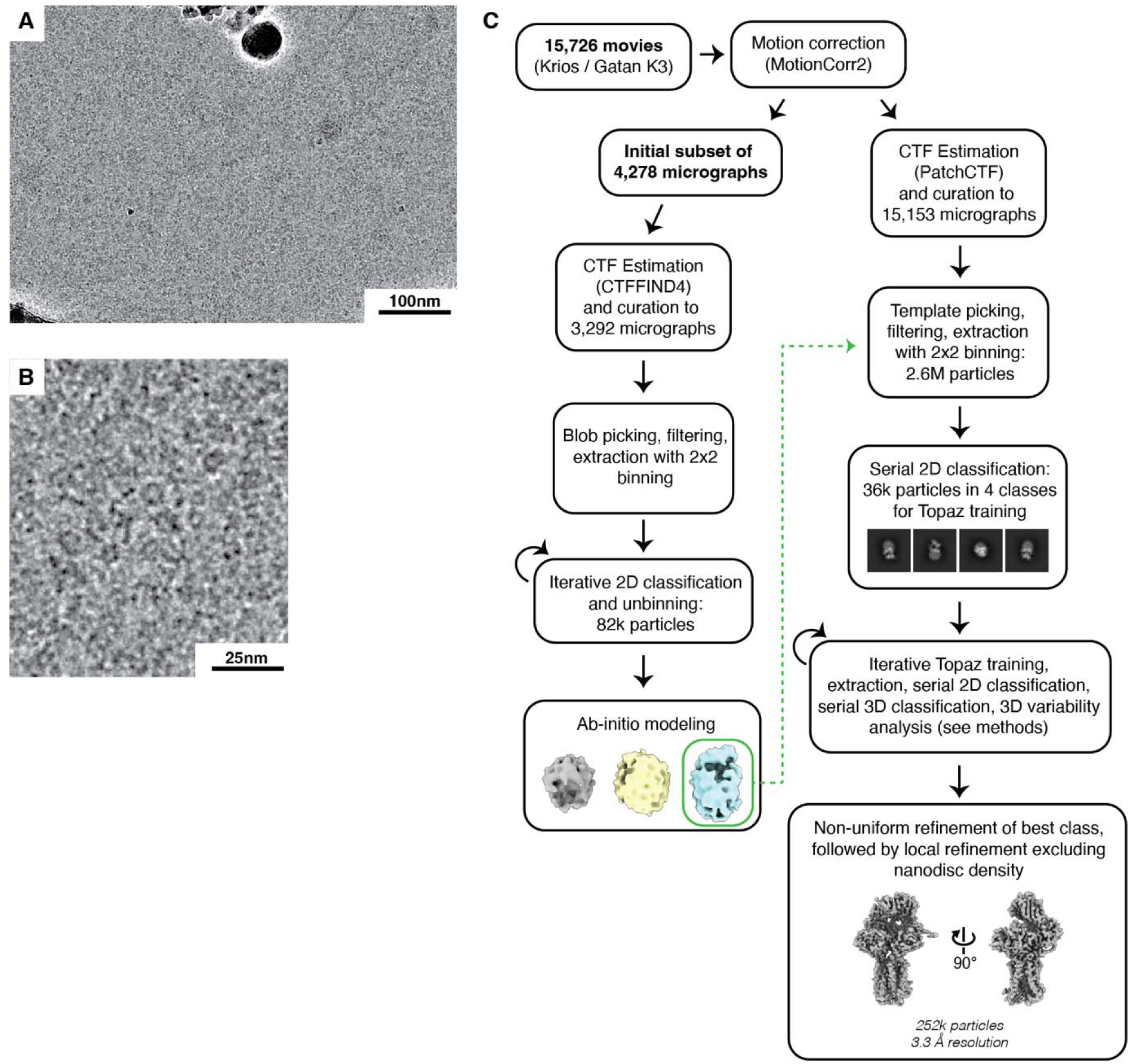
Cryo-EM image processing for the 1G4(LY) TCR–CD3 in detergent. (**A**) Representative cryo-electron micrograph from this study. (**B**) Zoomed-in region of the image in panel A. (**C**) Flow chart of data collection and image processing as described in Materials and Methods. Data were processed in cryoSPARC v4.

**Fig. S3:**
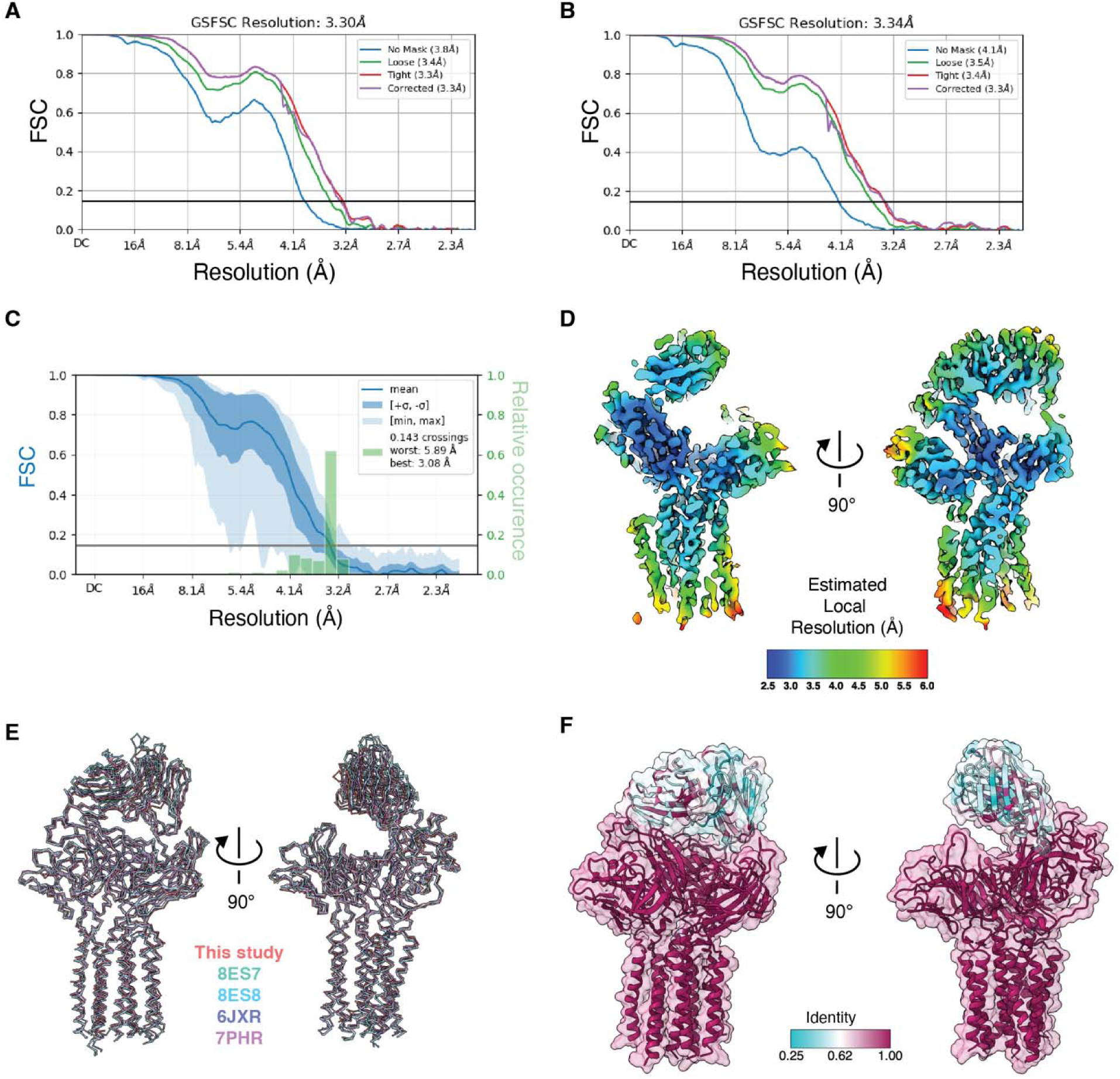
Structure of the 1G4(LY) TCR–CD3 in detergent. (**A**) Gold-standard FSC curves for the map of the TCR–CD3 in GDN (herein, simply “GDN”) determined in this study after non-uniform refinement. (**B**) FSC curve for the locally refined GDN map (mask excluded micelle density). While resolution was not nominally improved by masking out the micelle, map interpretability was qualitatively improved, and this map was used for further analysis. (**C**) 3D FSC analysis for the locally refined GDN map, showing relatively little map anisotropy. (**D**) Central slices of the locally refined GDN map, colored by local resolution estimate, as indicated in the panel. (**E**) Backbone representation of the GDN structure compared with those of TCR– CD3 sequences previously determined in other studies. Models containing HLA–β2M–antigen have had those atoms removed for this analysis. (**F**) GDN model colored according to sequence identity among the four TCR–CD3 sequences (this study and three priors) with cryo-EM structures determined in detergent micelles.

**Fig. S4:**
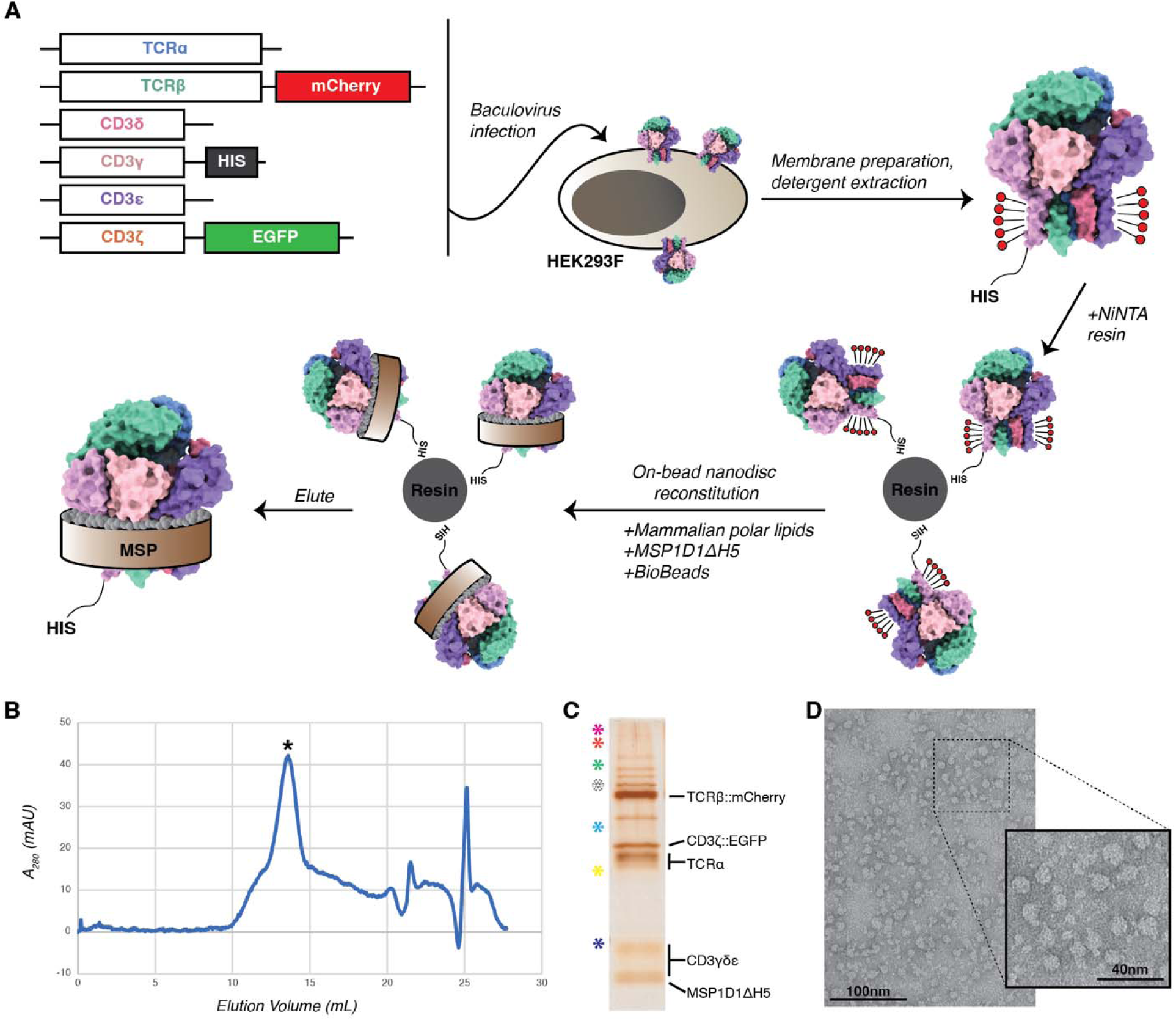
Overview of TCR–CD3 expression, purification, and reconstitution into nanodiscs. (**A**) Schematic representation of the expression and purification of TCR–CD3 and its reconstitution into nanodiscs as described in Materials and Methods. (**B**) Representative size-exclusion chromatogram of nanodisc-embedded TCR–CD3. Asterisk indicates peak of TCR– CD3 in nanodiscs. (**C**) Representative silver-stained SDS-PAGE gel of pooled size-exclusion chromatography peak fractions (right lane). Constituents identified by expected molecular weight. CD3γ, δ, and ε cannot be distinguished by this approach. Molecular weight ladder is in the left lane, with asterisks indicating the following molecular weight values: magenta, 250 kDa; orange, 150 kDa; green, 100 kDa; white, 75 kDa; cyan, 50 kDa; yellow, 37 kDa; navy, 25 kDa. (**D**) Representative negative-stain electron micrograph of pooled size-exclusion chromatography peak fractions. Inset, zoomed-in view. Note presumed TCR–CD3 density protruding from the circular nanodisc structures.

**Fig. S5:**
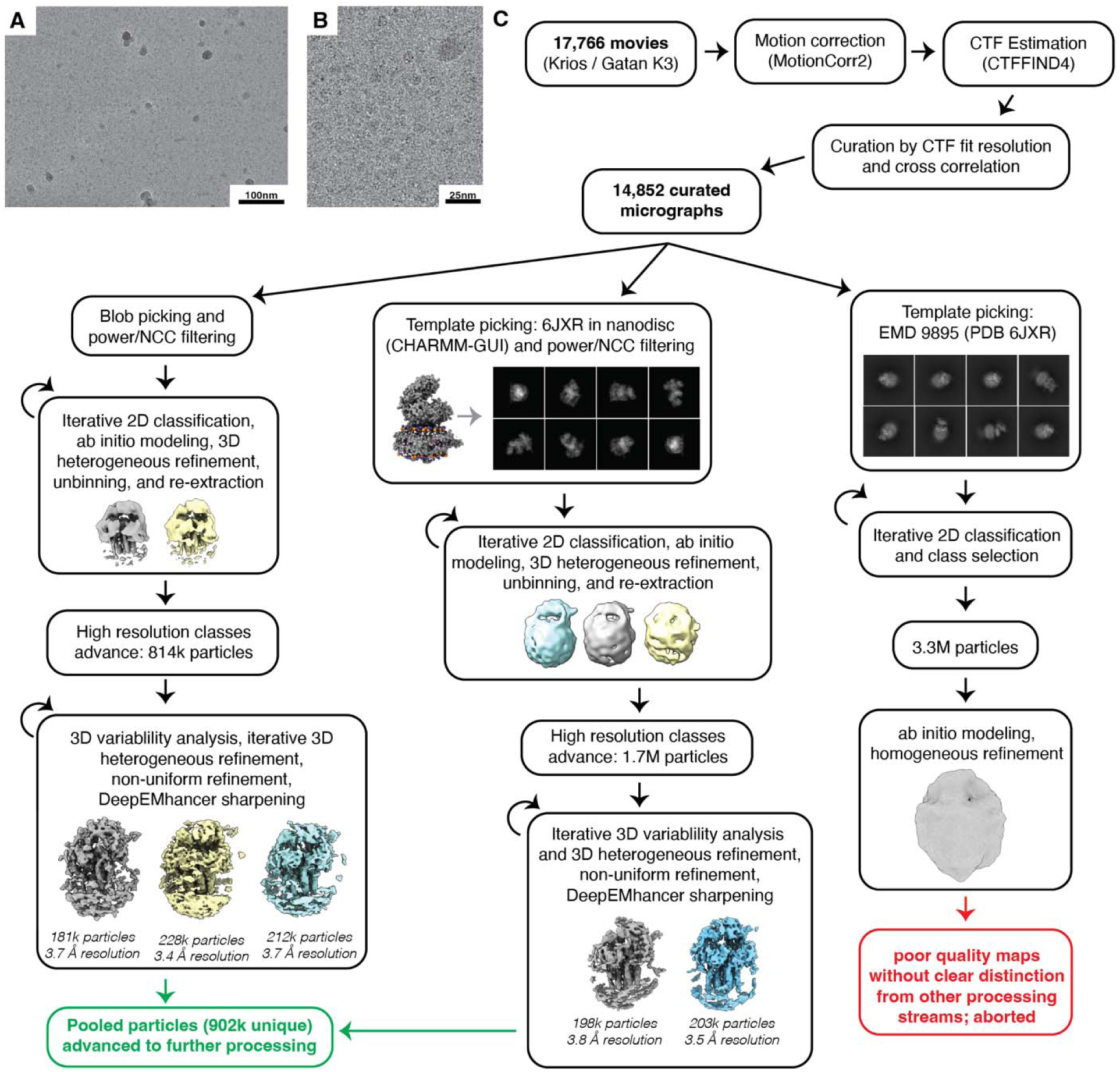
Initial cryo-EM image processing for the TCR–CD3 in nanodiscs. (**A**) Representative cryo-electron micrograph from this study. (**B**) Zoomed-in region of the image in panel B. (**C**) Flow chart of data collection and image processing as described in Materials and Methods. Data were processed in cryoSPARC v4 in three parallel streams using particle projections picked using an unbiased circular “blob” template (left), and using templates based on PDB 6JXR embedded in an MSP1D1DH5 nanodisc (center), or the original EM map used to create the PDB 6JXR model (right). The particle classes yielding the highest resolution reconstructions were pooled and advanced to further processing as detailed in Fig. S6.

**Fig. S6:**
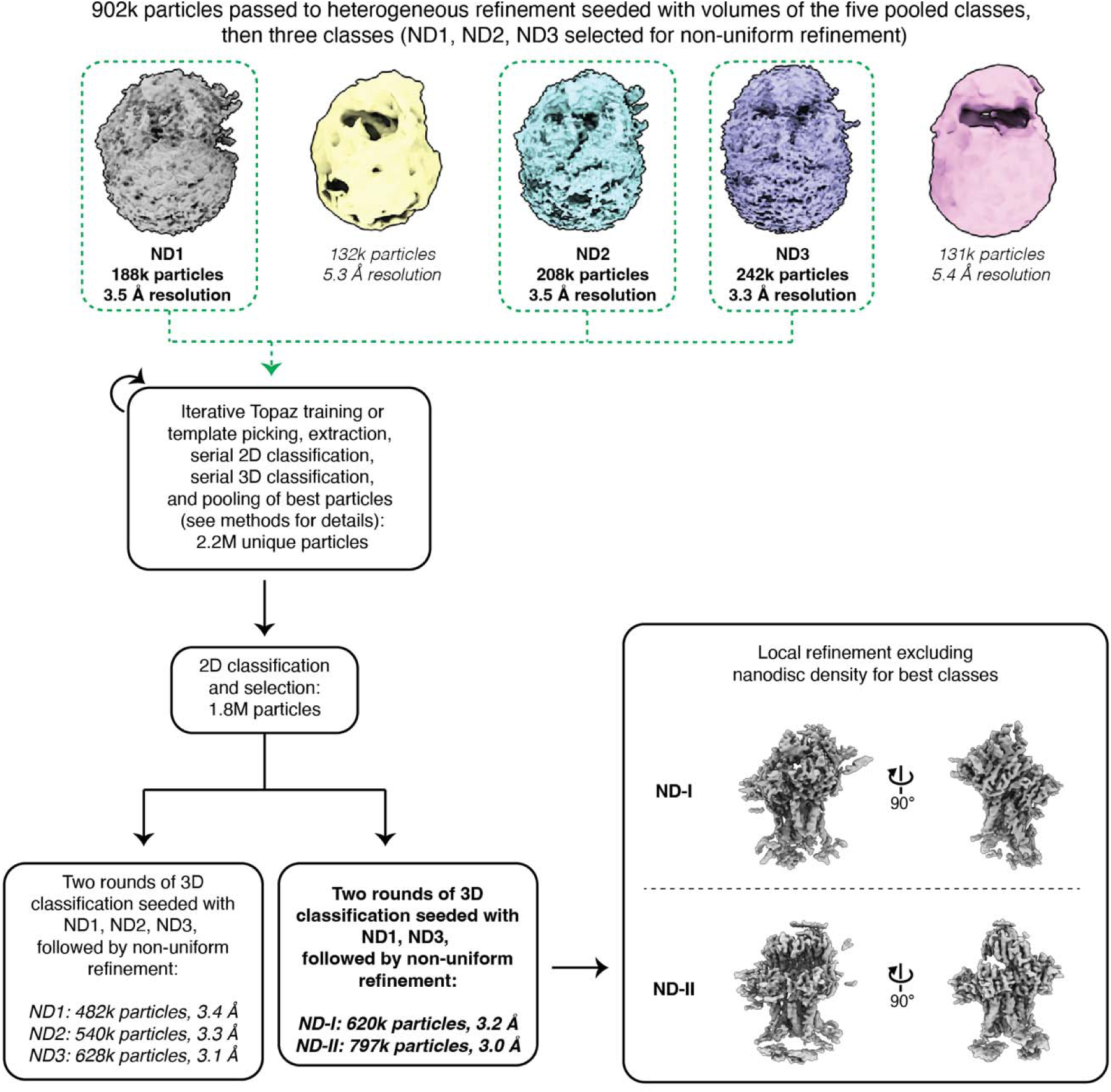
Further cryo-EM image processing for the TCR–CD3 in nanodiscs. Continued flow chart of image processing as described in Materials and Methods. Approximately 902,000 unique particles from the blob picking and TCR–CD3-in-nanodisc template picking processing streams (see Fig. S5) were pooled and their respective five maps were used to seed heterogeneous refinement in cryoSPARC. The five output maps were subjected to non-uniform refinement and the resulting maps are shown; ND1, ND2, and ND3 denote best refined classes, whose particles were used to seed further processing guided by Topaz particle picking (see Materials and Methods). Final rounds of 3D classification gave maps of superior quality and nominal resolution when only ND1 and ND3 were used as seeds. Locally refined, DeepEMhancer-sharpened maps shown for the final two classes, now denoted ND-I and ND-II.

**Fig. S7:**
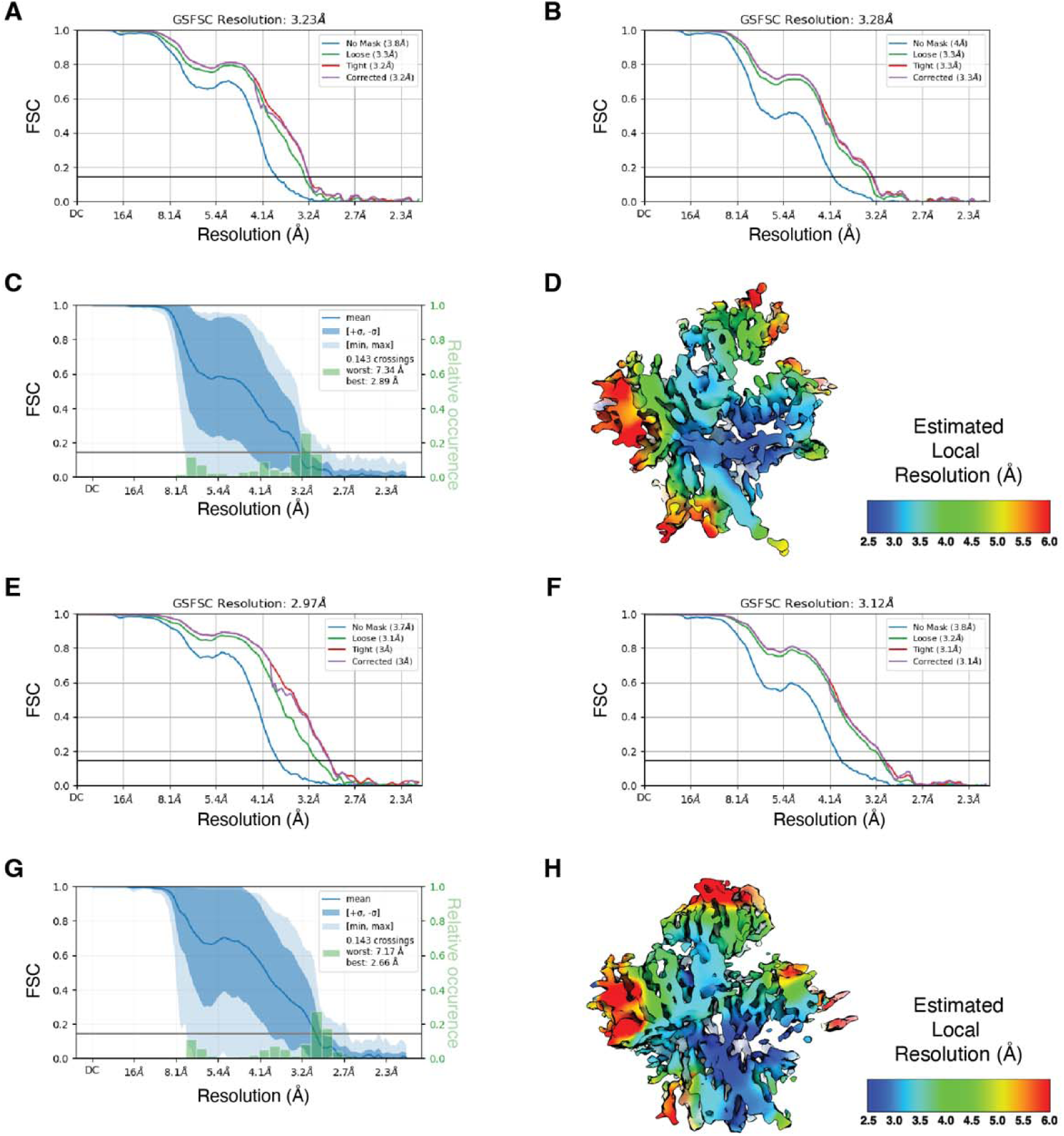
Map assessment for ND-I and ND-II. (**A**) Gold-standard FSC curves for the ND-I map after non-uniform refinement. (**B**) FSC curves for the locally refined ND-I map (mask excluded nanodisc density). While resolution was not nominally improved by masking out the nanodisc, map interpretability was qualitatively improved, and this map was used for further analysis. (**C**) 3D FSC analysis for locally refined ND-I. Note the more extensive anisotropy than was seen for the TCR–CD3 in detergent micelles. (**D**) Central slice of locally refined ND-I, colored by local resolution estimate, as indicated in the panel. (**E**) Gold-standard FSC curves for the ND-II map after non-uniform refinement. (**F**) FSC curves for the locally refined ND-II map (mask excluded nanodisc density). As with ND-I, nominal resolution was not improved by masking out the nanodisc, but map interpretability was qualitatively improved, and this map was used for further analysis. (**G**) 3D FSC analysis for locally refined ND-II. Note again the more extensive anisotropy than was seen for the TCR in detergent micelles. (**H**) Central slice of locally refined ND-II, colored by local resolution estimate, as indicated in the panel.

**Fig. S8:**
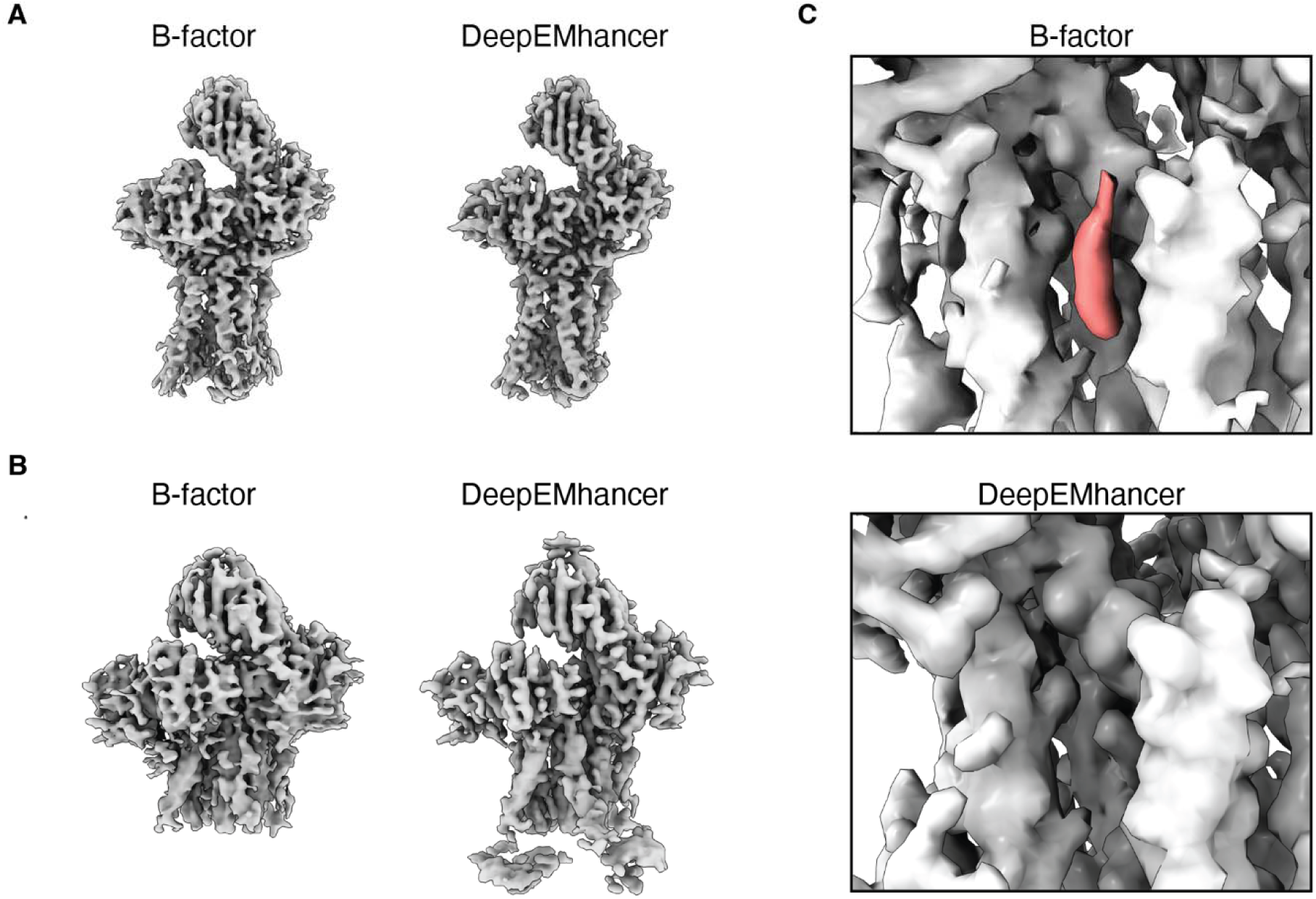
Cryo-EM map sharpening. (**A,B**) Comparison of global B-factor sharpening and DeepEMhancer sharpening for the TCR–CD3 in detergent (A) and ND-II (B). (**C**) Lipid elements are better discriminated by global B-factor sharpening than by DeepEMhancer, with the volume for the TCRβ-associated cholesterol in ND-II highlighted in red in the B-factor-sharpened map (top). In contrast, this cholesterol is not visualized after DeepEMhancer sharpening when contoured at a comparable threshold (bottom).

**Fig. S9:**
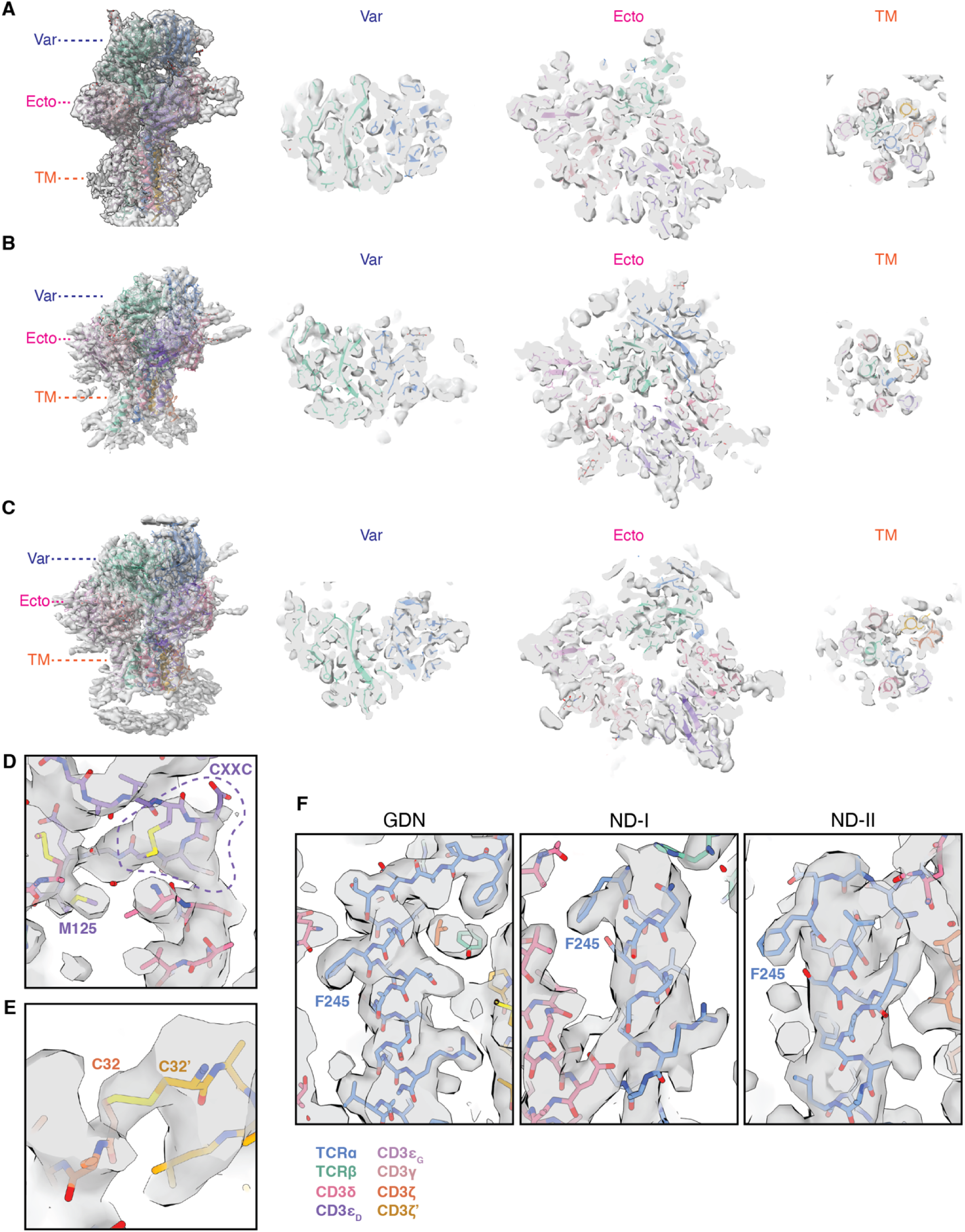
Model building and assessment for unliganded TCR–CD3 structures. (**A-C**) Overviews and sections through the DeepEMhancer-sharpened maps (gray, transparent surface) and models for the (A) GDN, (B) ND-I, and (C) ND-II conformations. Sections shown at right were taken at the approximate levels indicated on the overviews at left. Var, TCR variable domains; Ecto, TCR constant and CD3 ectodomains; TM, transmembrane helices. (**D**) The CD3 _G_ C-X-X-C lariat (enclosed by dotted line) is shown resting on the amino-terminus of the CD3δ TM helix, with additional confirmation of the register by bulky sidechains like CD3 _G_ Met125. (**E**) Modeling the interchain CD3ζ-CD3ζ’ disulfide bond establishes the register for these transmembrane helices, as the proteins lack folded ectodomains or juxtamembrane region C-X-X-C lariats. (**F**) Comparison of the density assigned to the TCRα TM helix from the indicated models, with the positioning of the Phe245 sidechain establishing the register for the helix in the absence of continuous juxtamembrane linker density in the ND-I and ND-II maps. Color legend at bottom applies to all panels in the figure.

**Fig. S10:**
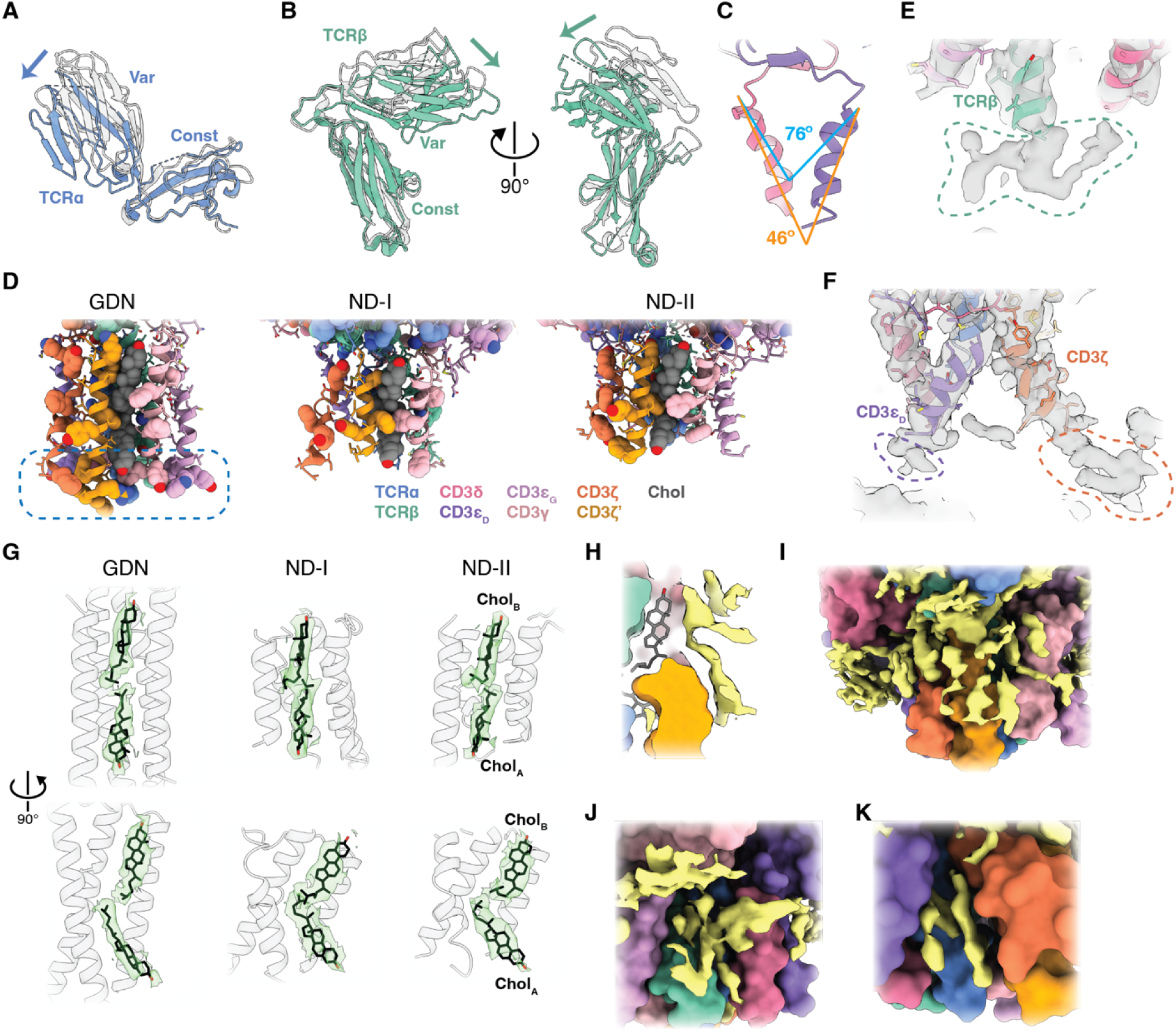
Structural detail of the TCR–CD3 in nanodiscs. (**A**) Cartoon representations of TCRα in ND-I (solid) and GDN (transparent gray), aligned on the TCRα constant domain. Arrows indicate the movements of the variable domains relative to the constant domains that comprise the “closure” of the variable domains in ND-I/II relative to GDN. (**B**) Same as in panel A but comparing TCRβ in the ND-I and GDN conformations. (**C**) The bent TM helices of the ND-I CD3 _D_ heterodimer. Splay measured at the level of the first helical turn (cyan) is greater than that of the helices overall (orange). (**D**) Cartoon representation of the indicated models with basic and aromatic residues highlighted as space-filling spheres. Blue dashed box highlights the clusters of basic and aromatic residues modeled at the carboxy-termini of the TM helices in GDN, but unstructured in ND-I and ND-II. Color legend applies to all figure panels. (**E,F**) Density projecting from the carboxy-termini of the TM helices modeled in ND-II that could not be confidently modeled, but likely corresponding to continuation of the polypeptide chains. Gray, DeepEMhancer-sharpened maps. Densities in question are encircled with dashed lines. (**G**) Density corresponding to the cholesterols modeled in ND-I, ND-II, and GDN. The transparent green surface is the indicated locally refined, B-factor-sharpened cryo-EM map, cropped to a zone of 2.2 Å around the cholesterols (black). (**H-K**) Additional densities likely attributable to lipids in the ND-II map. Proteins are shown as model surfaces, and cholesterol in stick representation. Yellow surface is the ND-II B-factor-sharpened map after local refinement (H) or non-uniform refinement (I-K) with the modeled densities subtracted to highlight unmodeled densities.

**Fig. S11:**
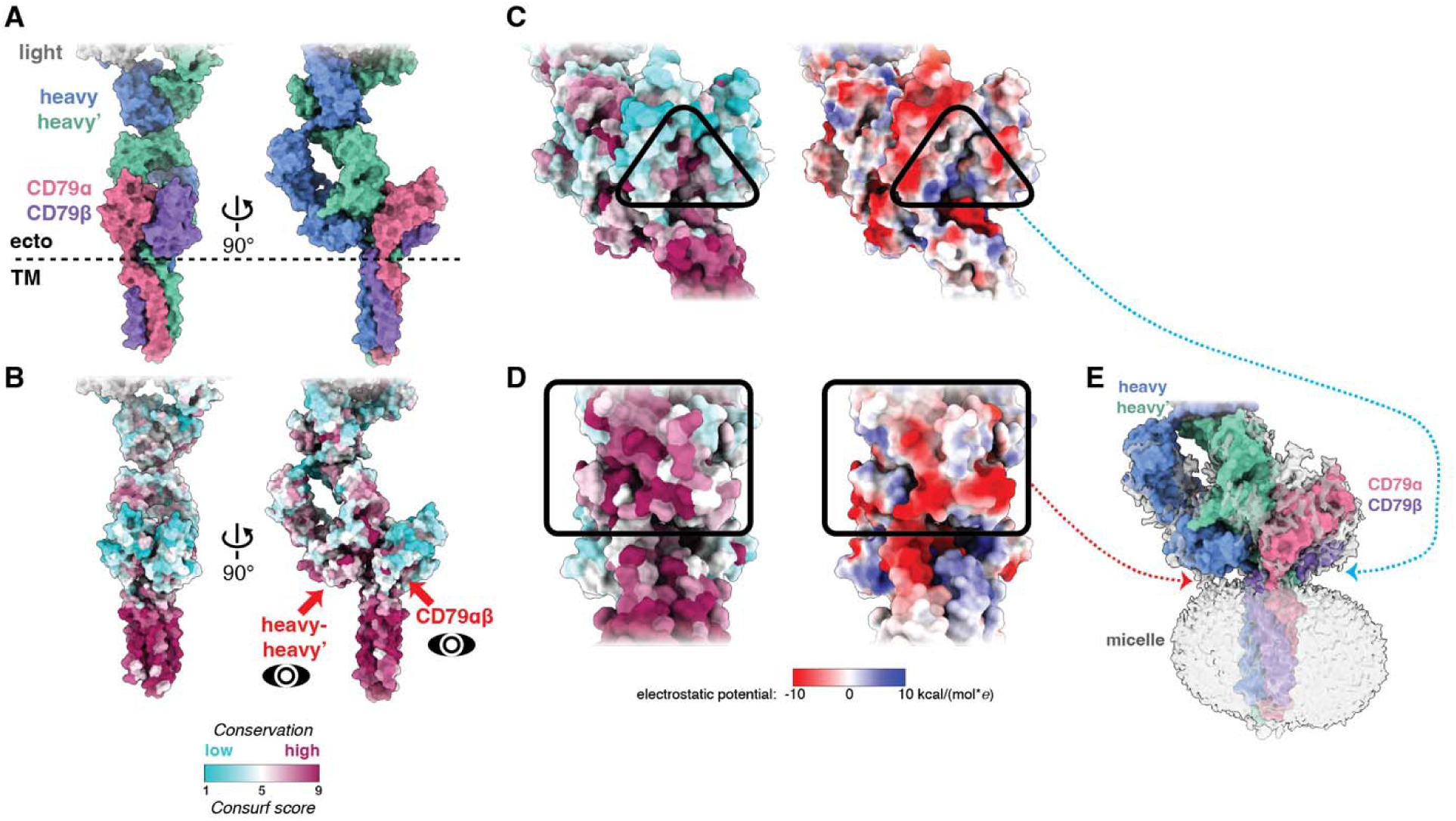
Insights into B-cell receptor structure and membrane interaction. (**A,B**) Surface representation of the human B-cell receptor (BCR) model, as determined in a detergent micelle by cryo-EM (PDB 7XQ8)(*23*). The model is colored by chain (A) and by sequence conservation (B, scale applies to remainder of figure). Red arrows and eye icons labeled CD79αβ and heavy-heavy’ indicate the direction for the magnified views in panels C and D, respectively. (**C**) Membrane-proximal face of the CD79αβ interface, colored by conservation (left, as in panel A) and electrostatic potential (right). Black triangle highlights region of high sequence conservation and corresponding surface electropositivity. However, this patch is oriented away from the micelle surface in the structure in detergent (dashed arrow to panel E). (**D**) Membrane-proximal face of the IgM heavy chains, with black box highlighting broad sequence conservation and largely electronegative conserved face. Dashed arrow indicates contact between this surface patch and the detergent micelle density in panel E. Electrostatic potential scale applies to panels C and D. (**E**) Surface model of the BCR with the unmodeled map density shown (transparent gray surface). Note the detergent micelle enshrouding the TM helices and making contact with the electronegative heavy-heavy’ interface (red dashed arrow) but not the electropositive CD79αβ interface (blue dashed arrow), which is the opposite of what might be expected in a phospholipid membrane.

**Fig. S12:**
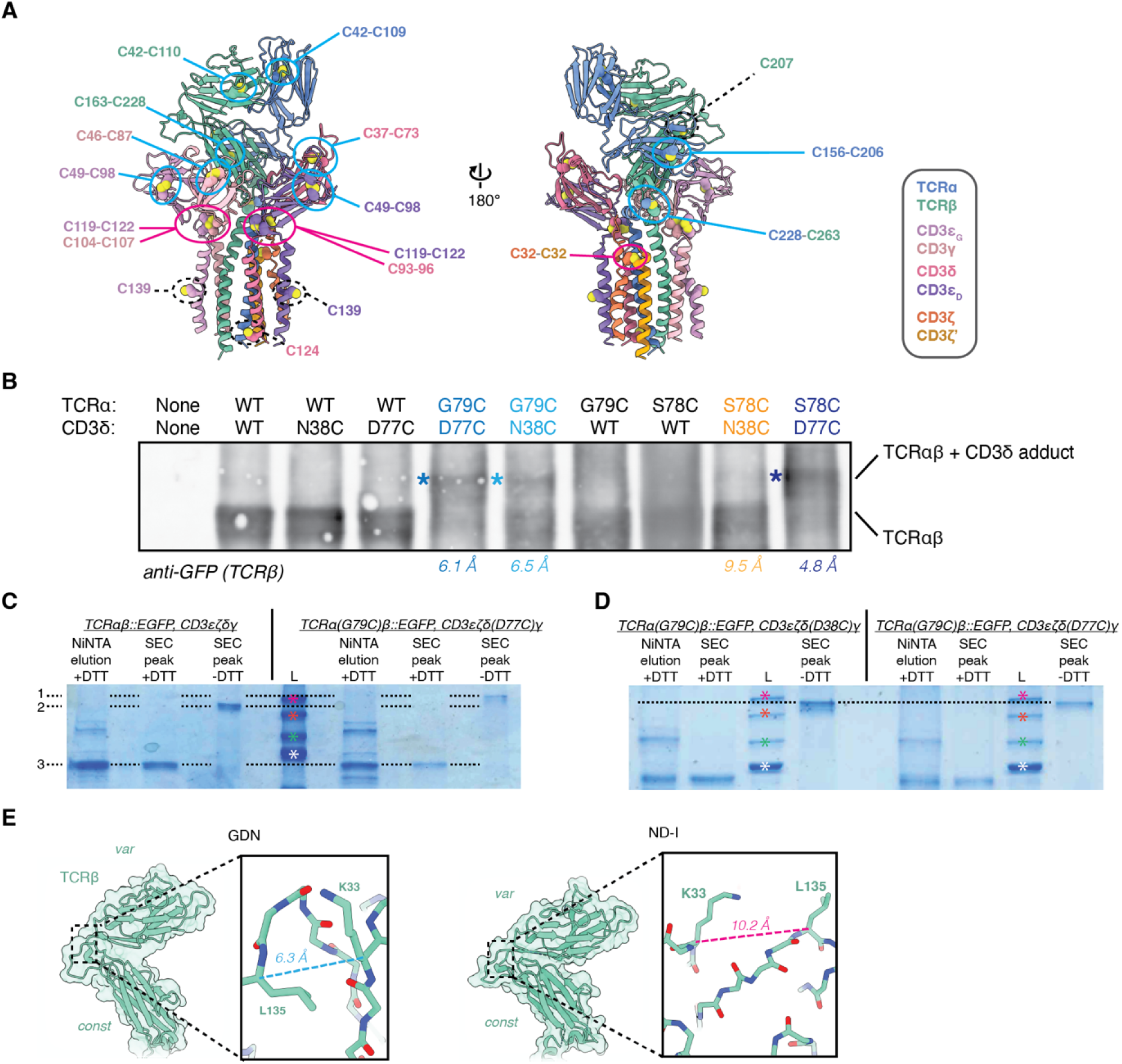
Engineered disulfide crosslinking. (**A**) Location of all native disulfides (solid cyan and magenta ovals) and unpaired cysteines (dashed black ovals) in the used TCR constructs. Disulfides indicated by cyan ovals are amenable to mass spectrometric detection as peptide pairs after EThcD, whereas disulfides indicated by magenta ovals are not unambiguously detectable by this approach, as they are C-X-X-C lariats or symmetric between CD3ζ homodimers. Subunits colored as indicated to the right. (**B**) Non-reducing SDS-PAGE gel of NiNTA affinity-purified TCR complex from HEK293T cells infected with the indicated TCR virus genotypes and Western blotted for EGFP-tagged TCRβ. Engineered disulfides yielding a TCRαβ–CD3δ covalent adduct results in an upward gel shift (asterisks). Color coding and distances as in Fig. 4A,B. (**C,D**) Acquastain-stained SDS-PAGE gels of NiNTA affinity-purified TCR complex reconstituted into nanodiscs and purified by size-exclusion chromatography (SEC). Virus genotypes indicated above gels. The presence (+) or absence (-) of reducing agent dithiothreitol (DTT) in the sample-loading buffer is indicated. In panel C, dashed line markers 1, 2, and 3 indicate the inferred electrophoretic mobility of the TCRαβ::EGFP–CD3δ adduct, TCRαβ::EGFP, and TCRβ::EGFP, respectively. Note the mobility shift between the nonreduced wild-type TCR complex and the non-reduced TCRα(G79C)–CD3δ(D77C) mutant complex. Molecular weight ladder (L) is color-coded with magenta, orange, green, and white asterisks denoting 250 kDa, 150 kDa, 100 kDa, and 75 kDa, respectively. In panel D, the non-reduced TCRα(G79C)–CD3δ(D38C) mutant complex displays an electrophoretic mobility similar to that of the non-reduced TCRα(G79C)–CD3δ(D77C) mutant complex, consistent with engineered disulfide bond formation. (**E**) Design of an engineered disulfide bond plausible in the GDN conformation but not in the closed ND-I conformation.

**Fig. S13:**
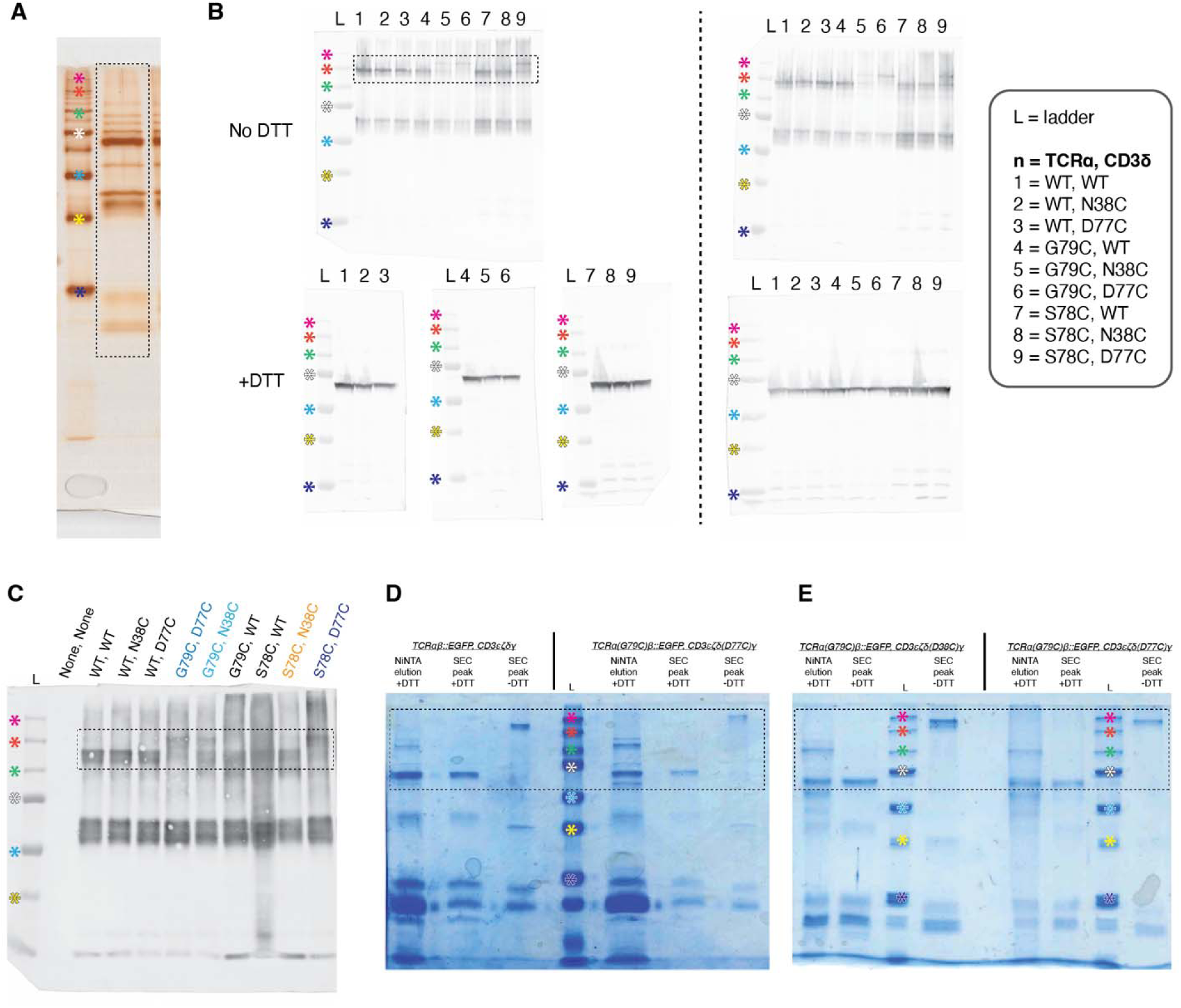
Full-length gels and blots. (**A**) Full-length silver-stained gel corresponding to Fig. S4C. Asterisks indicate the following molecular weight ladder values in all panels: magenta, 250 kDa; orange, 150 kDa; green, 100 kDa; white, 75 kDa; cyan, 50 kDa; yellow, 37 kDa; navy, 25 kDa. For all panels, black dashed box indicates approximate region shown in corresponding main text figures and preceding supplementary figures. (**B**) Full-length blots corresponding to the Western blot in Fig. 4B (upper left) and a replicate (upper right). Additionally, blots were performed using lysates from the same infected cells boiled under reducing conditions (“with DTT”), showing disruption of the TCRαβ–CD3δ covalent complexes (bottom). (**C**) Full-length blot corresponding to Fig. S12B. Lane labels indicate TCRα and CD3δ genotypes, color-coded as in Fig. S12. (**D**) Full-length gel corresponding to Fig. S12C. (**E**) Full-length gel corresponding to Fig. S12D.

**Fig. S14:**
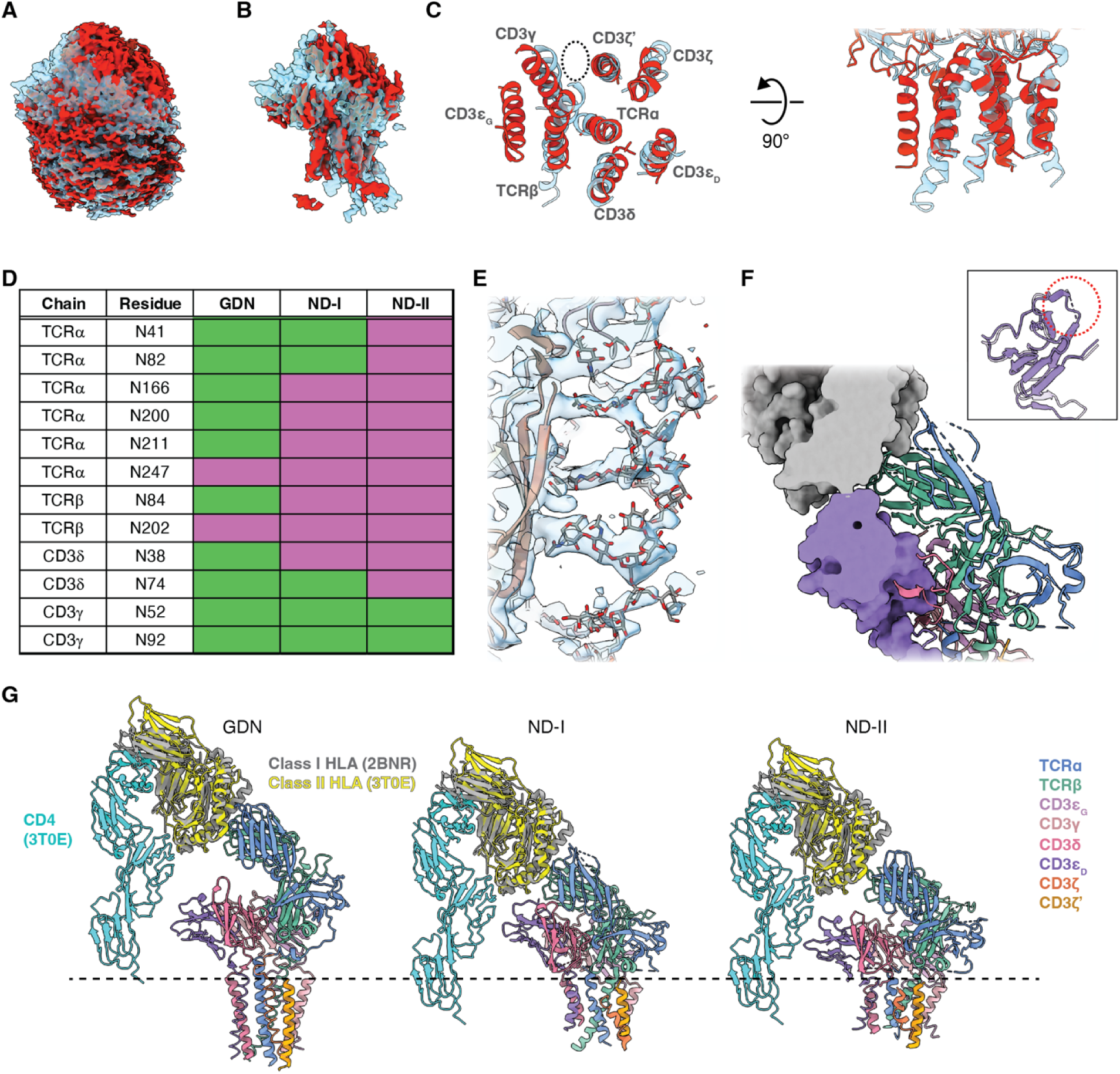
Conformational variation in the resting state and glycosylation of the TCR–CD3. (**A,B**) Cryo-EM maps for ND-I (transparent cyan surface) and ND-II (solid red surface) aligned to the TCRα transmembrane helix. The B-factor-sharpened map without local refinement, shown in panel A, is contoured to highlight alignment of the nanodisc density. The DeepEMhancer-sharpened, locally refined map, shown in panel B, is contoured to highlight protein density. (**C**) Relative positions of the TM helices for ND-I and ND-II (colored as in panel A) viewed along the axis perpendicular to the membrane plane (left) and parallel to the membrane (right). Dashed circle indicates cholesterol-binding groove. (**D**) Graphical representation of partially modeled N-linked glycans in GDN, ND-I, and ND-II. All asparagine residues within an N-X-S/T motif in the structure are included in this list for completeness; however, TCRα N247 is located in the TM domain and is unlikely to undergo glycosylation. Green, modeled; magenta, not modeled. (**E**) Glycans (gray sticks) in a structured array on the surface of HIV Env as determined by x-ray crystallography (PDB 5FYJ)(*26*). Protein shown as cartoon backbone. 2Fo-Fc map contoured to 1σ and clipped to within 5 Å of the model. (**F**) Space-filling model of HLA docked onto ND-I with the CD3 _D_ ectodomain from GDN docked in place of that from ND-I, showing minimal, if any steric clashing. Cartoon representation of the remainder of ND-I shown for orientation. Inset shows alignment of CD3 _D_ from GDN and ND-I by their ectodomains. Red dashed circle highlights surface loop (residues 107-109) not modeled in ND-I (purple dashed line) but modeled in GDN. (**G**) Alignment of the CD4–Class II HLA–TCR ternary complex structure (PDB 3T0E)(*27*) to the Class I HLA docked onto our structures as in panel F and Fig. 5. The Class II HLA from 3T0E is shown in yellow, and CD4 in cyan. Other color coding as indicated. Dashed black line represents the approximate boundary of the lipid bilayer outer leaflet.

**Fig. S15:**
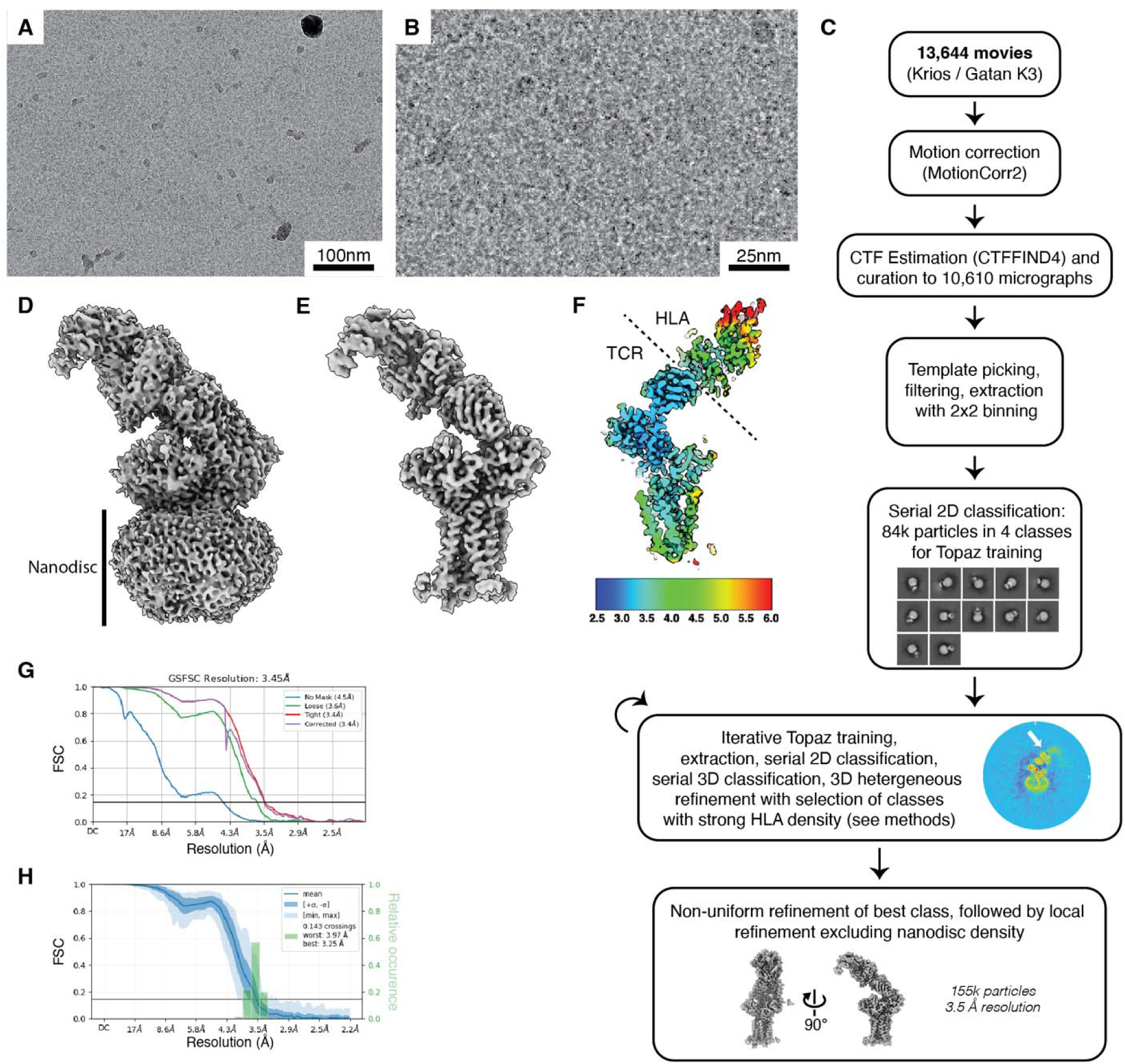
Cryo-EM image processing and map quality assessment for the TCR–CD3 in nanodiscs bound to HLA. (**A**) Representative cryo-electron micrograph from this study. (**B**) Zoomed-in region of the image in panel B. (**C**) Flow chart of data collection and image processing as described in Materials and Methods. Data were processed in cryoSPARC v4. White arrow highlights HLA density in a reconstruction central section. (**D**) B-factor sharpened map for the TCR–CD3 in nanodiscs bound to HLA shown with a low contouring threshold to highlight the density corresponding to the nanodisc. (**E**) Same view as in panel D but using the DeepEMhancer-sharpened map to highlight underlying protein density. (**F**) Central slice of the locally refined HLA-bound map colored by local resolution estimate in Å. (**G**) FSC curves for the locally refined map of HLA-bound TCR–CD3 complex (herein HLA-bound) in nanodiscs. Local refinement mask excluded nanodisc density. (**H**) 3D FSC analysis for the locally refined HLA-bound map.

**Fig. S16:**
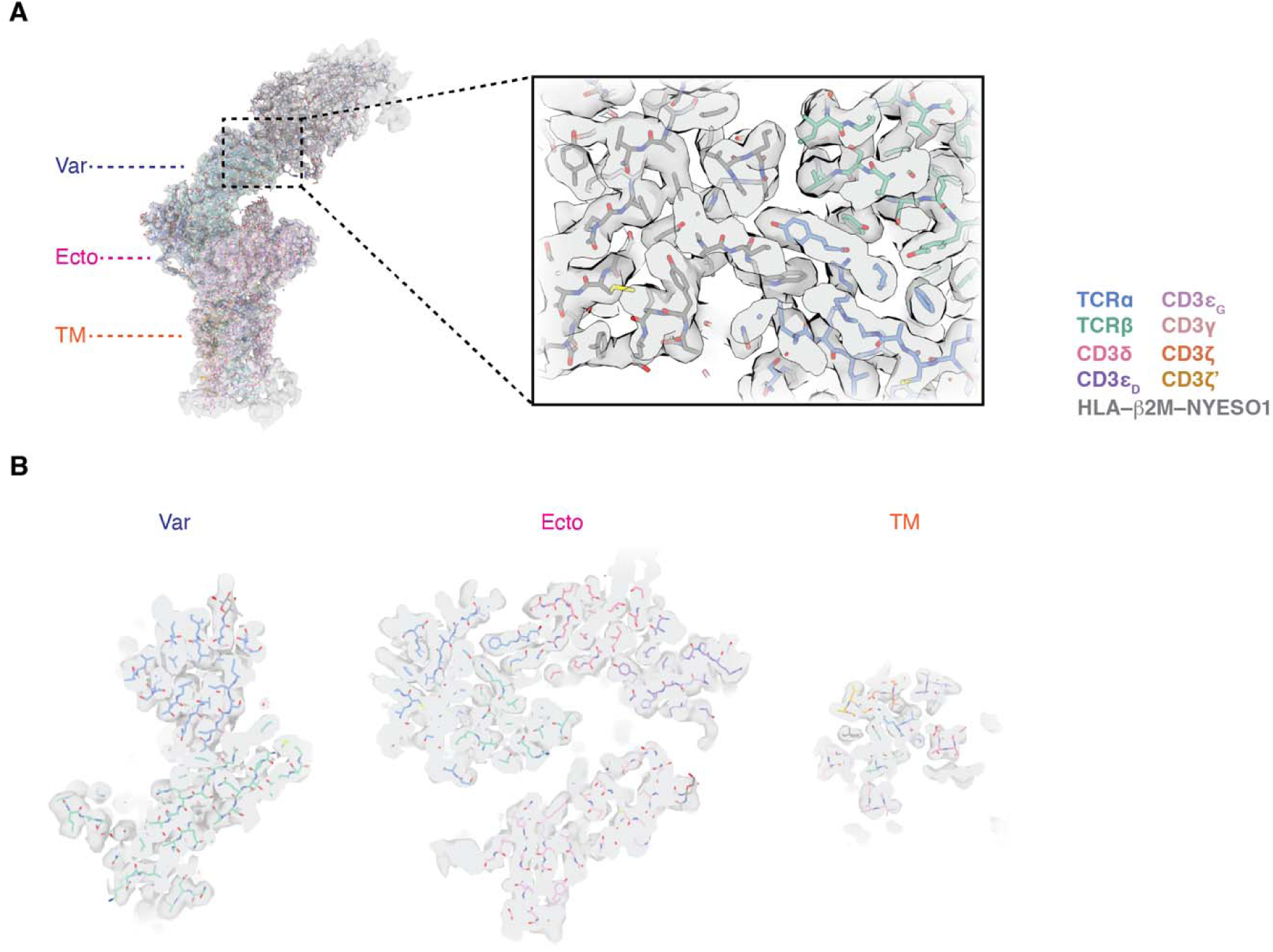
Model building and assessment for the TCR–CD3–HLA complex in nanodiscs. (**A,B**) Overview and sections through the DeepEMhancer-sharpened map (gray, transparent surface) and model. Sections shown in panel B were taken at the approximate levels indicated on the overview in panel A. Var, TCR variable domains; Ecto, TCR constant and CD3 ectodomains; TM, transmembrane helices. Inset in panel A shows a zoomed-in view of the TCRαβ–HLA–NY-ESO-1 interface. Color legend at right in panel A applies to models in both panels.

**Fig. S17:**
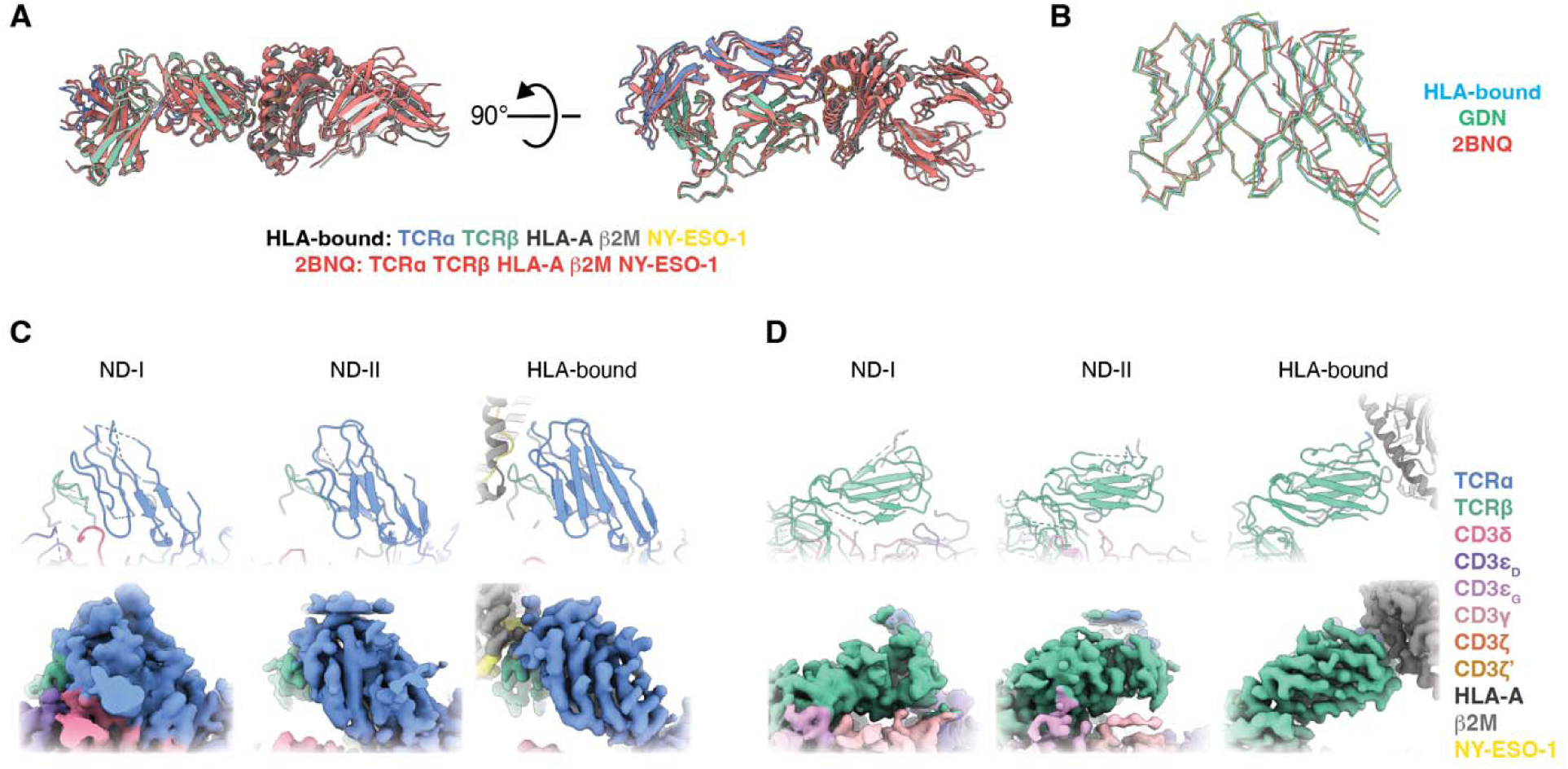
Additional structural analysis of the TCR–CD3 complex in nanodiscs bound to HLA. (**A**) Cartoon representation of the TCR variable domains and HLA from the HLA-bound structure (colored by chain) and the NY-ESO-1 TCR–HLA complex crystal structure (2BNQ(*51*)), colored red. Structures were aligned to the TCR variable domains. (**B**) Backbone traces of the TCR variable domains from HLA-bound, GDN, and the NY-ESO-1 TCR–HLA complex crystal structure (2BNQ(*51*)). Structures were aligned to the TCRα variable domain. (**C,D**) Comparison of the cryo-EM density for the TCR variable domains in HLA-bound and resting state subconformations zoomed on TCRα (C) and TCRβ (D). Top row, cartoon representation of model; bottom row, cryo-EM map. Legend in panel D also applies to panel C.

**Table S1:** Cryo-EM data collection parameters for the indicated datasets.

|  | **TCR-CD3 in GDN** | **TCR-CD3 in nanodiscs** | **TCR-CD3 in nanodiscs incubated with HLA** |
| --- | --- | --- | --- |
| Microscope | Titan Krios | Titan Krios | Titan Krios |
| Voltage (kV) | 300 | 300 | 300 |
| Camera | K3 | K3 | K3 |
| Magnification | 64,000 | 64,000 | 64,000 |
| Pixel size (Å) | 1.08 | 1.08 | 1.08 |
| Dose rate (e<sup>-</sup>/s/pixel) | 30 | 30 | 30 |
| Exposure time (s) | 2.0 | 2.2 | 2.0 |
| Frame rate (ms) | 40 | 40 | 40 |
| Electron exposure (e<sup>-</sup>/Å<sup>2</sup>) | 51 | 57 | 51 |
| Defocus range (µm) | -0.8 – -2.5 | -0.8 – -2.0 | -0.8 – -2.5 |
| Movie stacks | 15,726 | 17,766 | 13,644 |

**Table S2:** Model building and refinement statistics.

|  | **GDN** | **ND-I** | **ND-II** | **HLA-bound** |
| --- | --- | --- | --- | --- |
| **Model** |  |  |  |  |
| Composition (#) |  |  |  |  |
| Chains | 10 | 12 | 10 | 12 |
| Atoms | 8440 | 6402 | 6242 | 10620 |
| Residues | Protein: 1052 | Protein: 828 | Protein: 821 | Protein: 1356 |
|  | Nucleotide: 0 | Nucleotide: 0 | Nucleotide: 0 | Nucleotide: 0 |
| Water | 0 | 0 | 0 | 0 |
| Ligands | BMA: 3 | BMA: 1 | BMA: 1 | BMA: 1 |
|  | NAG: 15 | NAG: 9 | NAG: 3 | NAG: 10 |
|  | CLR: 2 | CLR: 2 | CLR: 2 | CLR: 2 |
| Bonds (RMSD) |  |  |  |  |
| Length (Å) (# > 4σ) | 0.003 (0) | 0.004 (0) | 0.003 (0) | 0.005 (0) |
| Angles (degrees) (# > 4σ) | 0.923 (6) | 0.862 (6) | 0.906 (3) | 0.892 (15) |
| MolProbity score | 1.9 | 2.74 | 2.67 | 2.22 |
| Clash score | 13.57 | 21.31 | 18.43 | 16.01 |
| Ramachandran plot (%) |  |  |  |  |
| Outliers | 0.1 | 0.26 | 0.4 | 0.15 |
| Allowed | 3.67 | 7.33 | 9.49 | 4.18 |
| Favored | 96.24 | 92.42 | 90.12 | 95.67 |
| Rama-Z<br>(Ramachandran plot<br>Z-score, RMSD) |  |  |  |  |
| whole (N = 1036) | 0.51 (0.26) | -2.23 (0.27) | -1.91 (0.29) | -0.18 (0.23) |
| helix (N = 214) | 0.63 (0.34) | -1.96 (0.42) | -1.14 (0.41) | 1.17 (0.31) |
| sheet (N = 286) | 1.85 (0.32) | -0.40 (0.37) | 0.54 (0.37) | 0.86 (0.25) |
| loop (N = 536) | -0.64 (0.26) | -1.87 (0.26) | -2.35 (0.28) | -1.39 (0.23) |
| Rotamer outliers (%) | 0.56 | 3.91 | 2.99 | 1.92 |
| Peptide plane (%) |  |  |  |  |
| Cis proline/general | 11.6/0.2 | 6.2/0.0 | 3.3/0.0 | 8.6/0.1 |
| Twisted<br>proline/general | 0.0/0.1 | 0.0/0.0 | 0.0/0.3 | 0.0/0.2 |
| CaBLAM outliers (%) | 3.04 | 5.63 | 7.32 | 3.45 |
| ADP (B-factors) |  |  |  |  |
| Iso/Aniso (#) | 8440/0 | 6402/0 | 6242/0 | 10620/0 |
| min/max/mean |  |  |  |  |
| Protein | 78.97/260.88/146.95 | 46.79/362.28/143.66 | 51.96/387.42/144.45 | 30.00/252.96/136.11 |
| Nucleotide | --- | --- | --- | --- |
| Ligand | 92.44/292.30/180.12 | 78.92/388.12/221.92 | 124.98/195.38/160.46 | 104.78/245.72/168.92 |
| Water | --- | --- | --- | --- |
| Occupancy |  |  |  |  |
| Mean | 1 | 1 | 1 | 1 |
| occ = 1 (%) | 100 | 100 | 100 | 100 |
| 0 < occ < 1 (%) | 0 | 0 | 0 | 0 |
| occ > 1 (%) | 0 | 0 | 0 | 0 |
| **Data** |  |  |  |  |
| Box: Lengths (Å) | (88.56, 101.52, 138.24) | (95.04, 87.48, 110.16) | (92.88, 92.88, 101.52) | (89.64, 128.52, 168.48) |
| Supplied Resolution (Å) | 3.3 | 3.3 | 3.1 | 3.5 |
| Resolution Estimates (Å), Masked |  |  |  |  |
| d FSC (half maps; 0.143) | 3.4 | 3.2 | 3.1 | 3.5 |
| d 99 (full/half1/half2) | 4.3/3.3/3.3 | 4.5/3.7/3.7 | 4.2/3.4/3.4 | 4.2/2.4/2.4 |
| d model | 3.7 | 3.3 | 3.1 | 3.9 |
| d FSC model (0/0.143/0.5) | 3.2/3.3/3.8 | 3.2/3.4/7.4 | 3.1/3.3/7.2 | 3.4/3.4/3.8 |
| Resolution Estimates (Å), Unmasked |  |  |  |  |
| d FSC (half maps; 0.143) | 3.5 | 3.3 | 3.1 | 3.6 |
| d 99 (full/half1/half2) | 4.3/2.5/2.5 | 4.5/3.5/3.4 | 4.2/3.2/3.2 | 4.2/2.3/2.3 |
| d model | 3.8 | 3.3 | 3.1 | 3.9 |
| d FSC model (0/0.143/0.5) | 3.3/3.4/3.8 | 3.2/3.5/7.6 | 3.1/3.4/7.3 | 3.4/3.5/4.0 |
| Map min/max/mean | -0.60/1.36/0.06 | -0.65/1.68/0.09 | -0.67/1.77/0.10 | -0.82/1.83/0.05 |
| **Model vs. Data** |  |  |  |  |
| CC (mask) | 0.82 | 0.57 | 0.59 | 0.79 |
| CC (box) | 0.81 | 0.65 | 0.68 | 0.74 |
| CC (peaks) | 0.70 | 0.41 | 0.48 | 0.60 |
| CC (volume) | 0.82 | 0.58 | 0.59 | 0.78 |
| Mean CC for ligands | 0.72 | 0.56 | 0.58 | 0.78 |

**Table S3:**
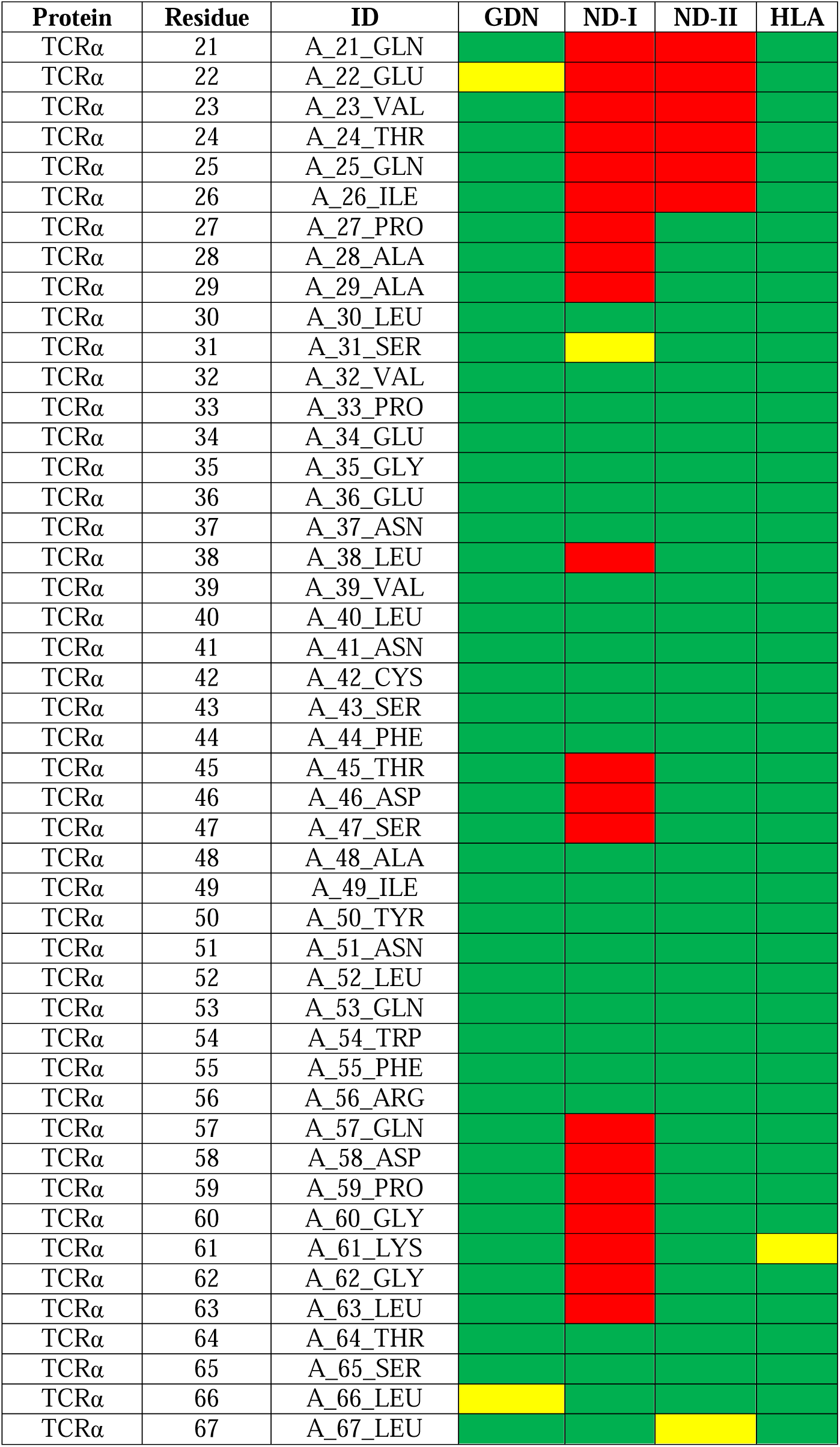
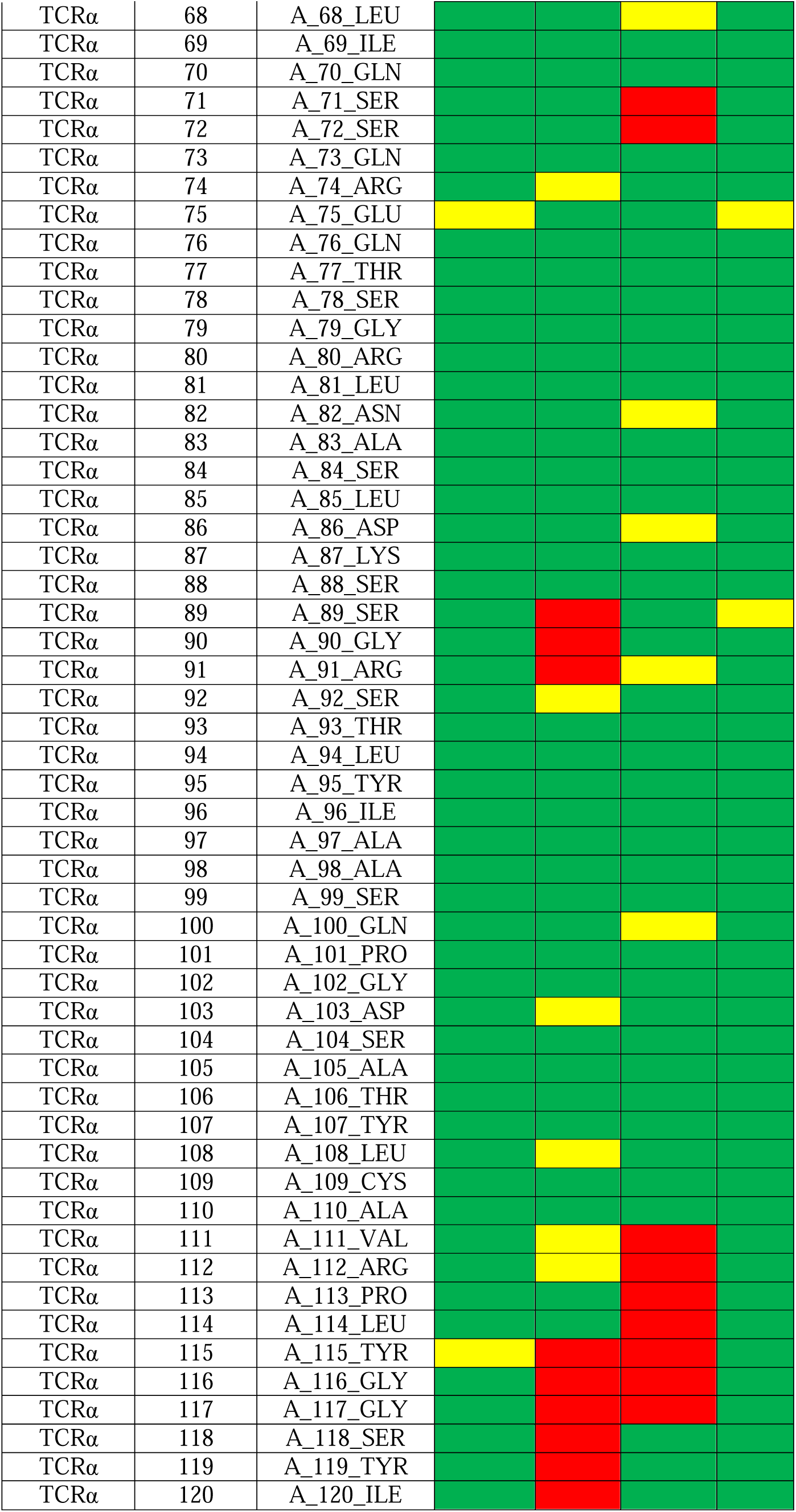
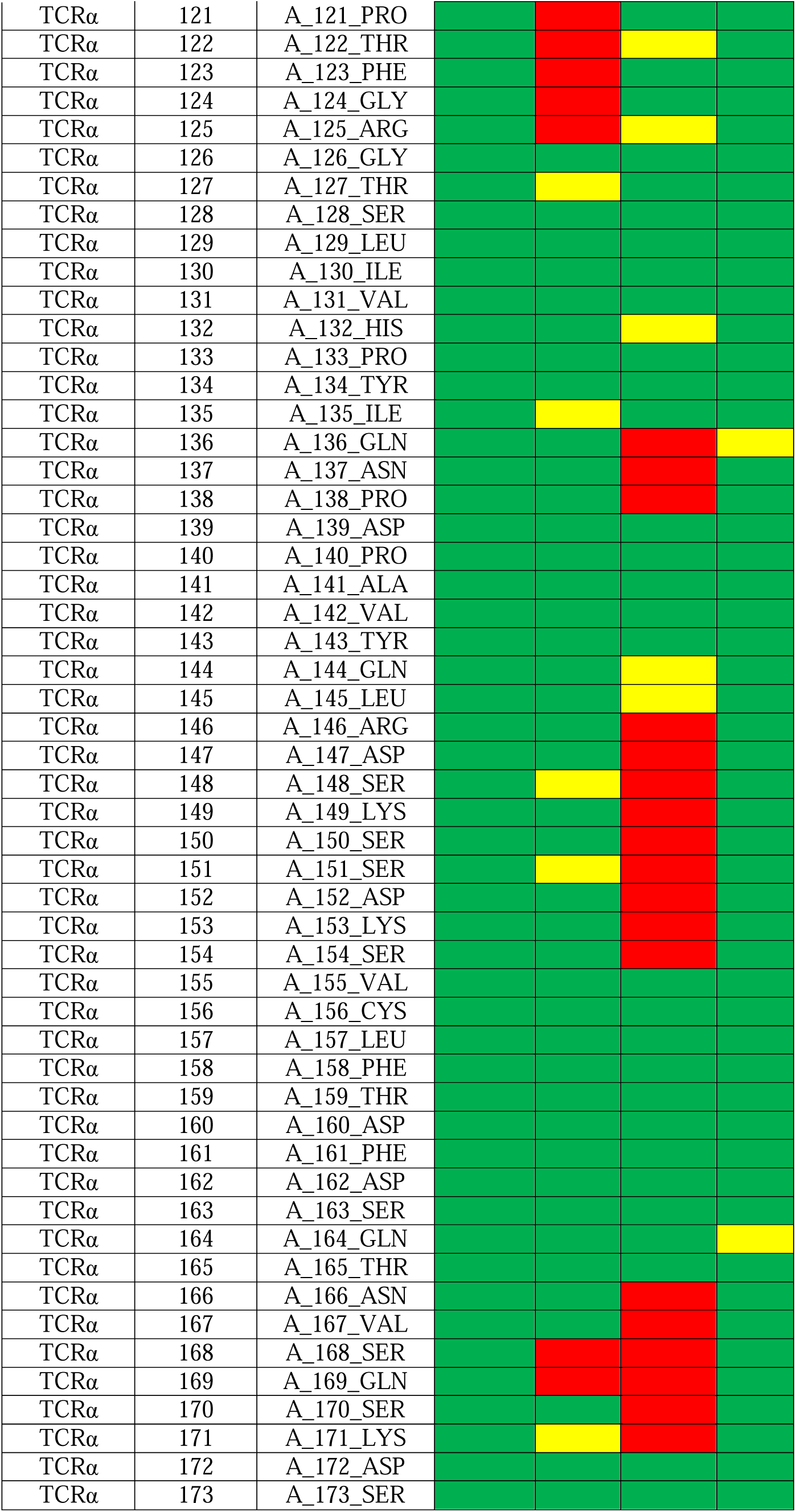
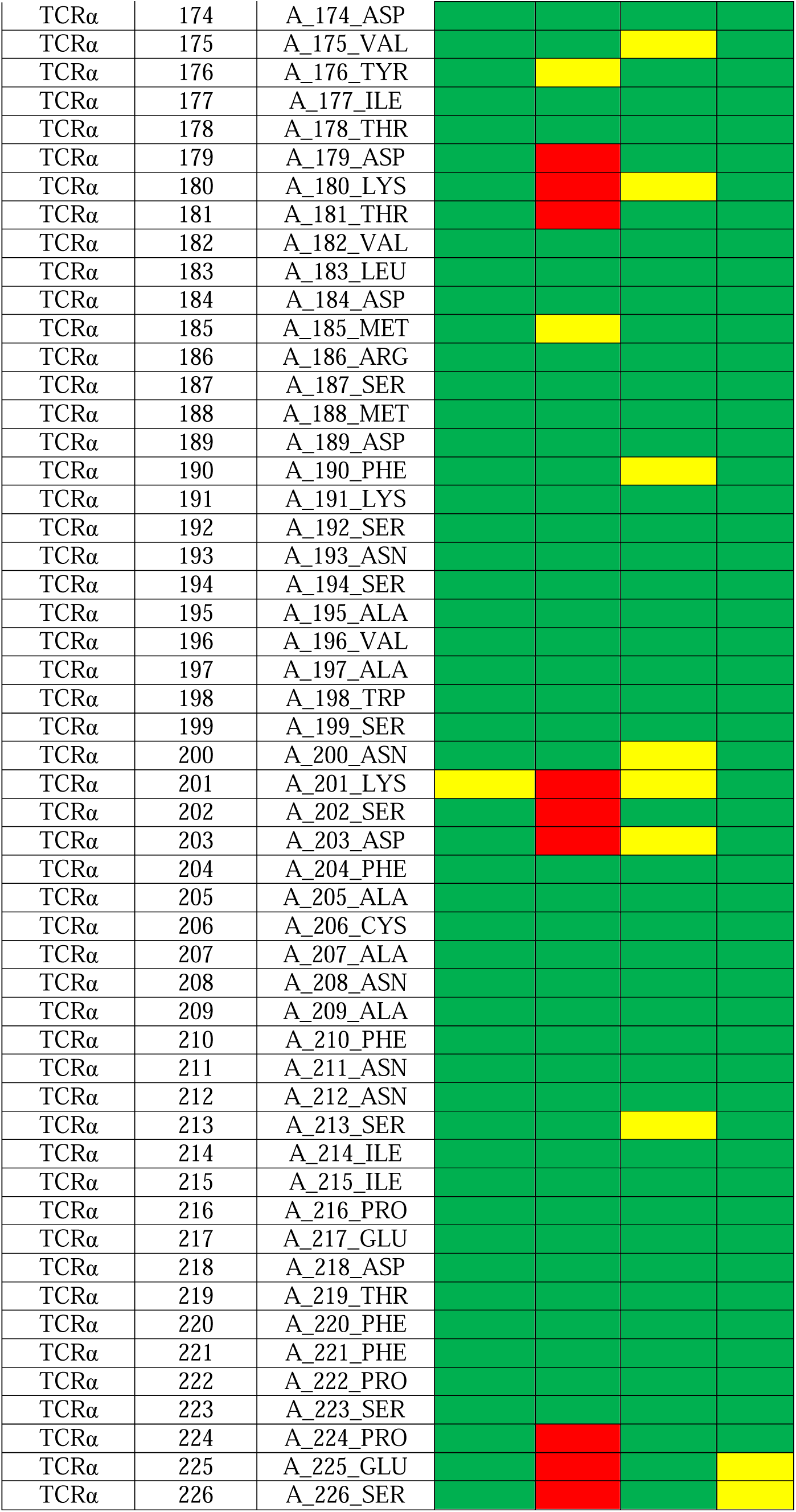
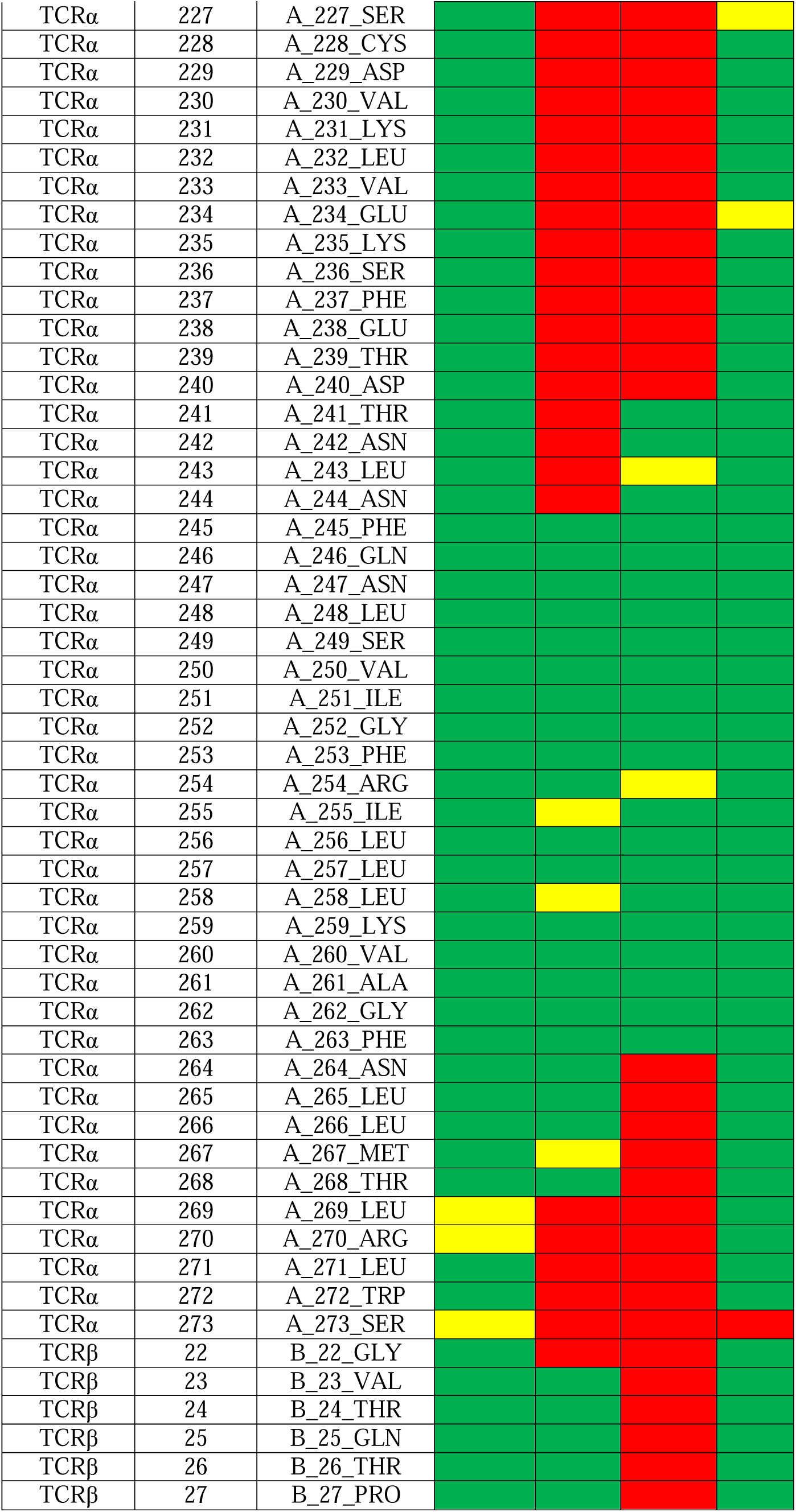
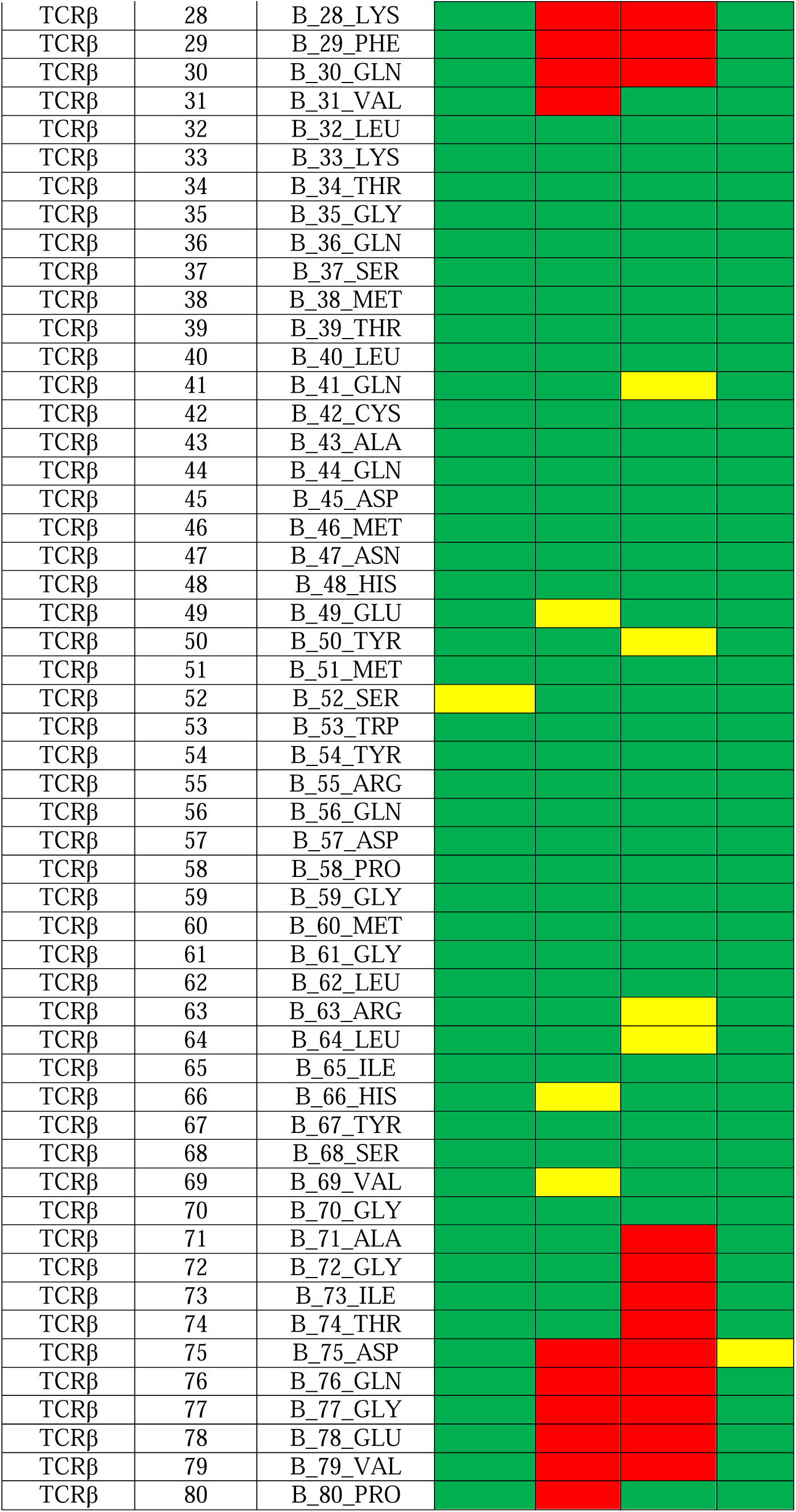
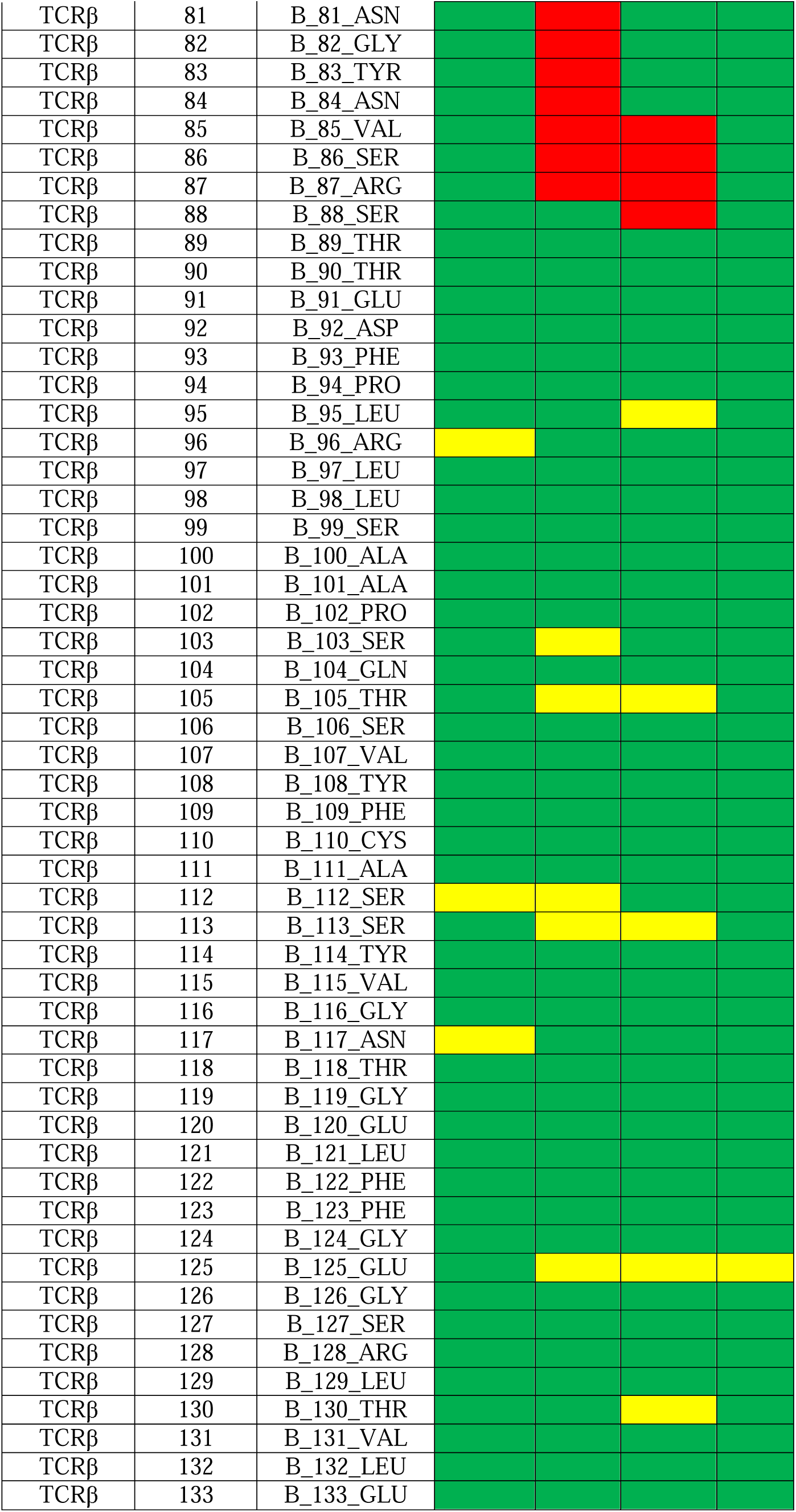
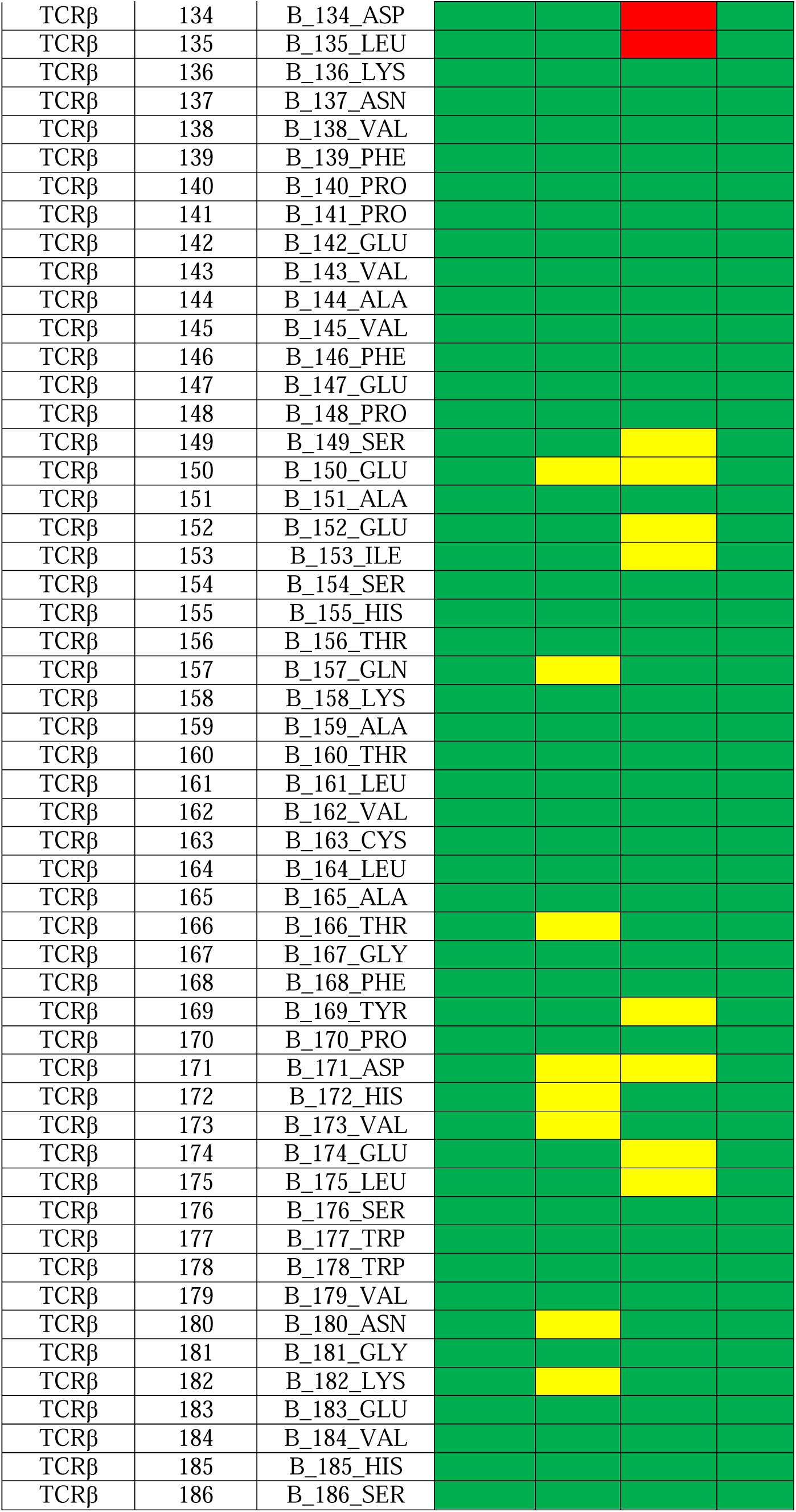
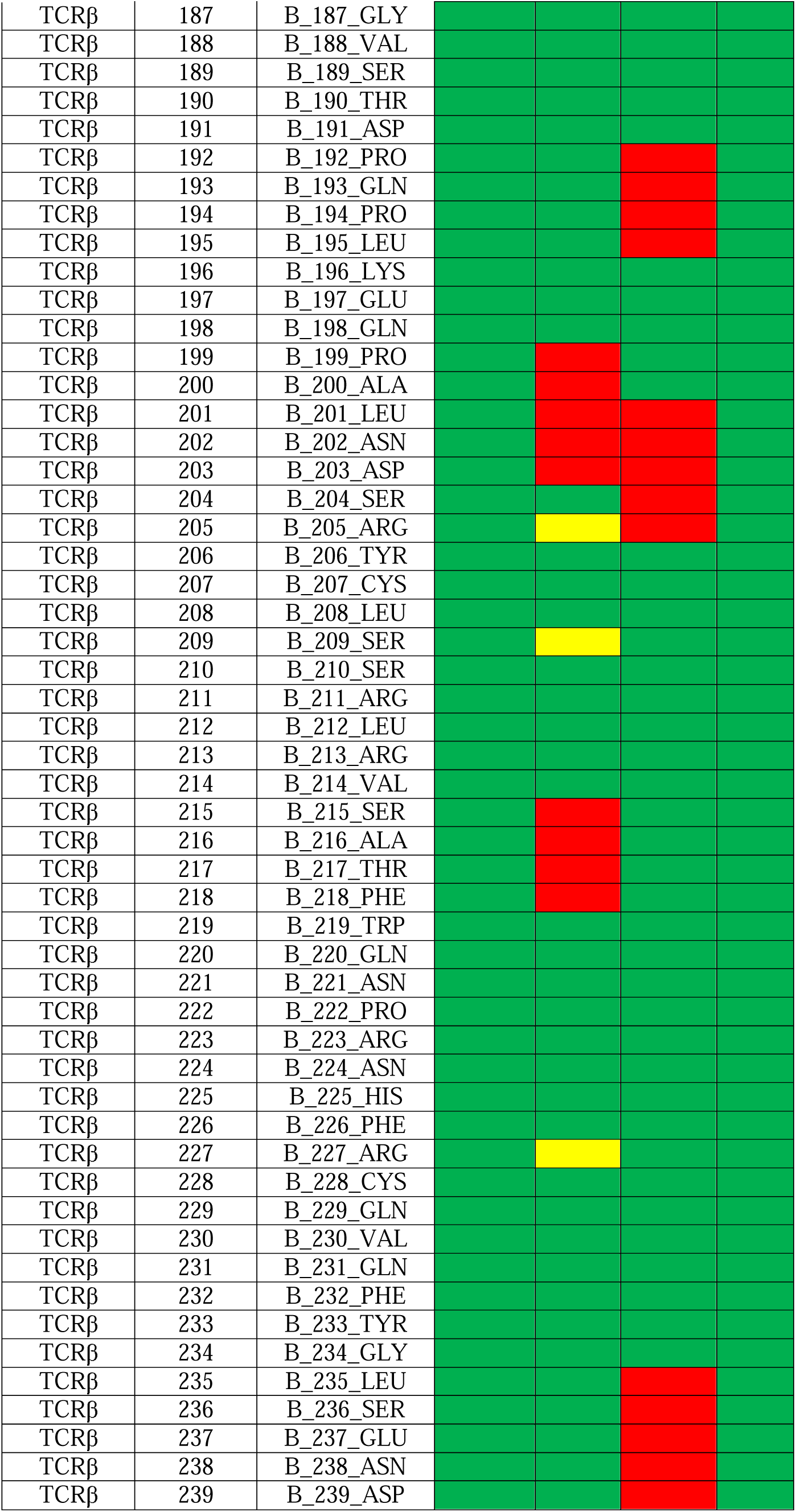
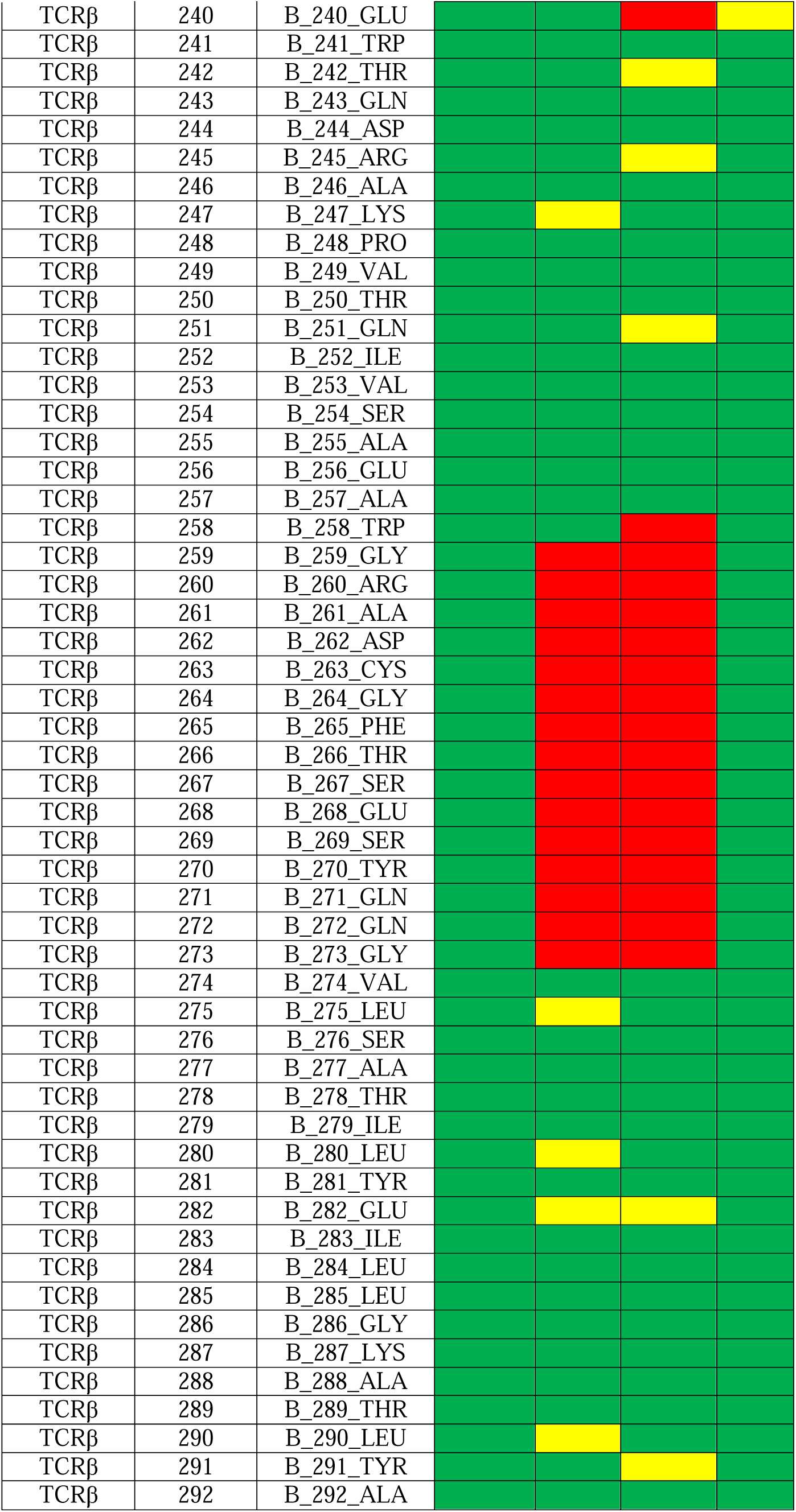
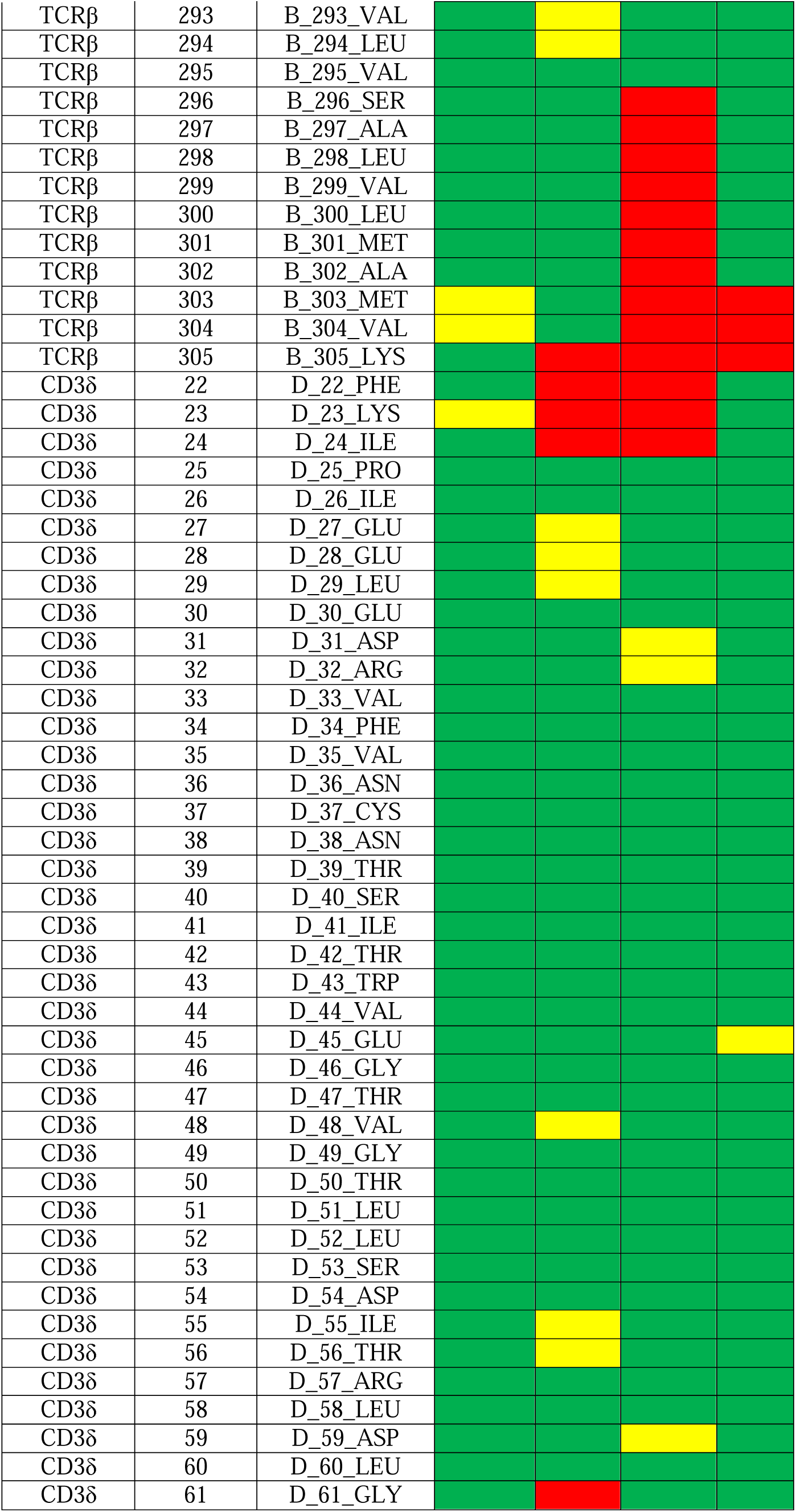
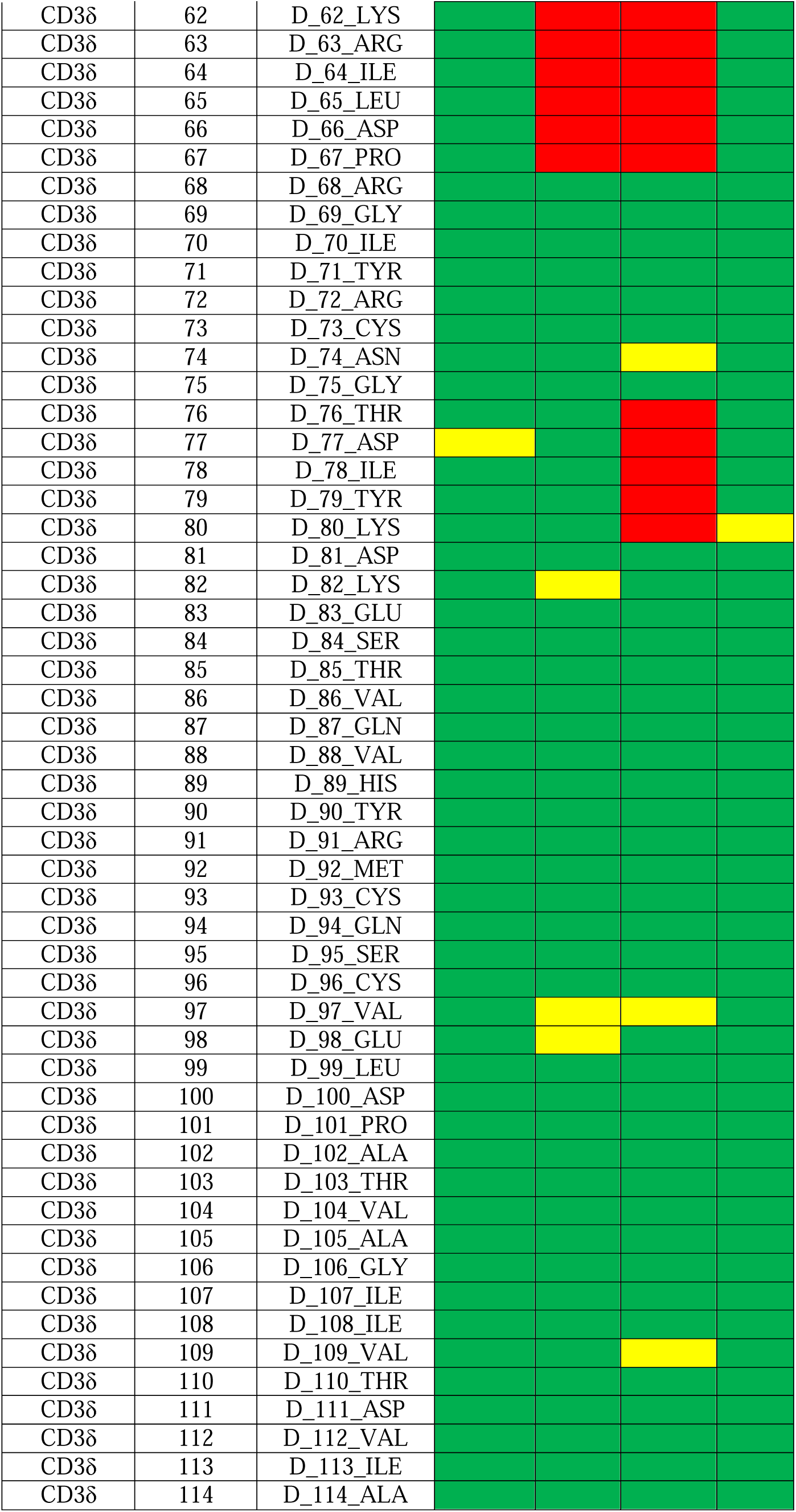
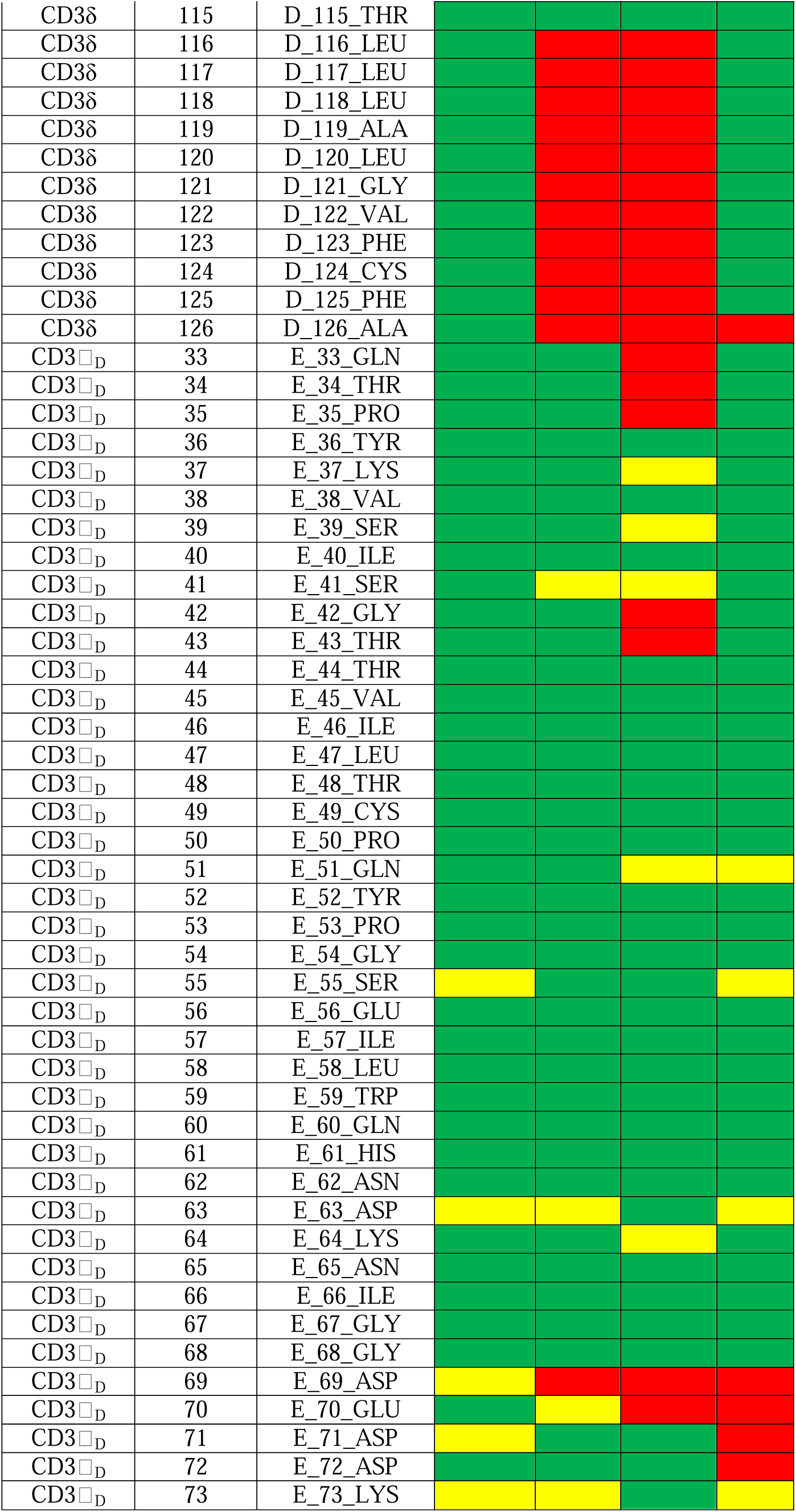
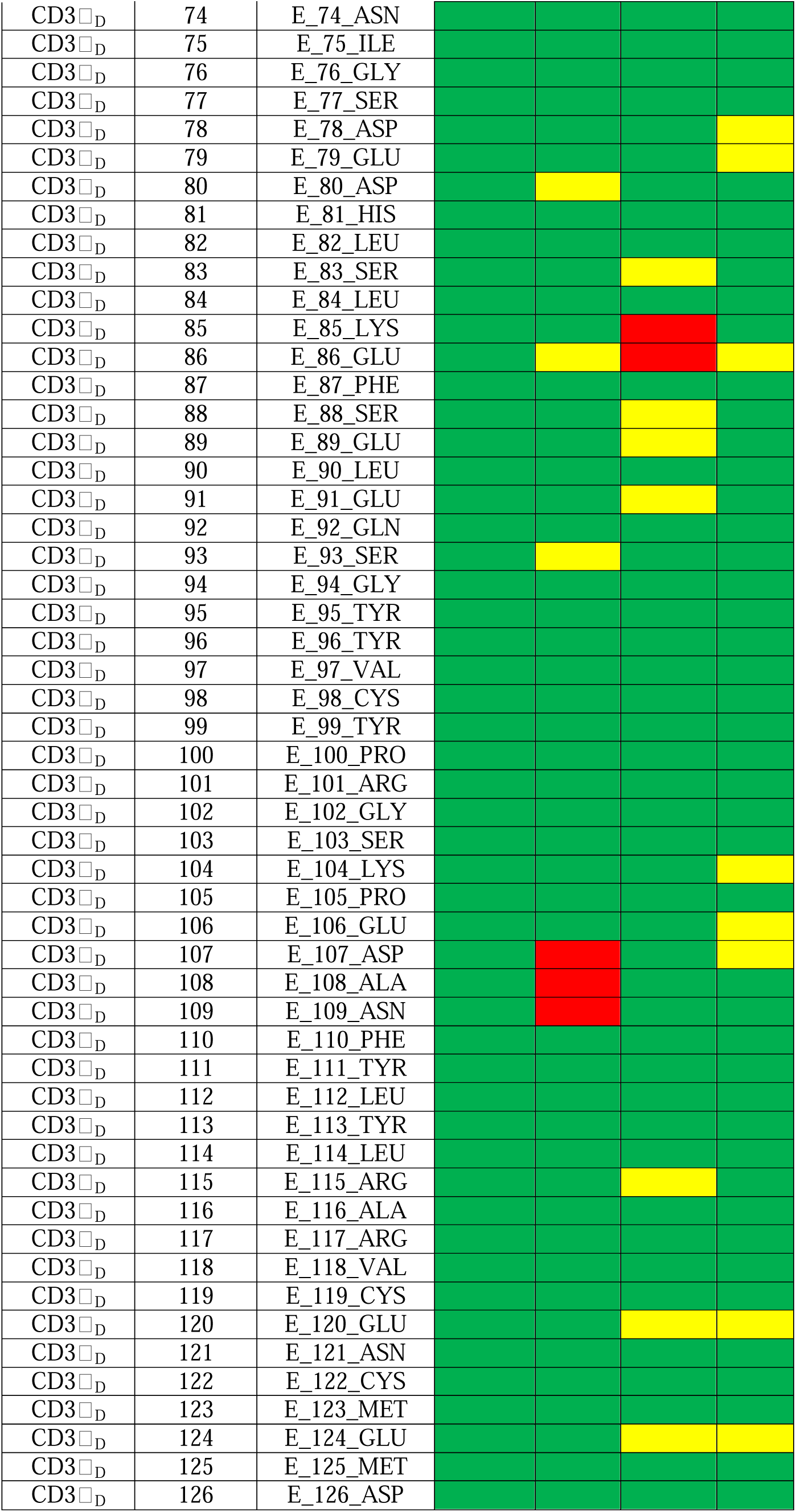
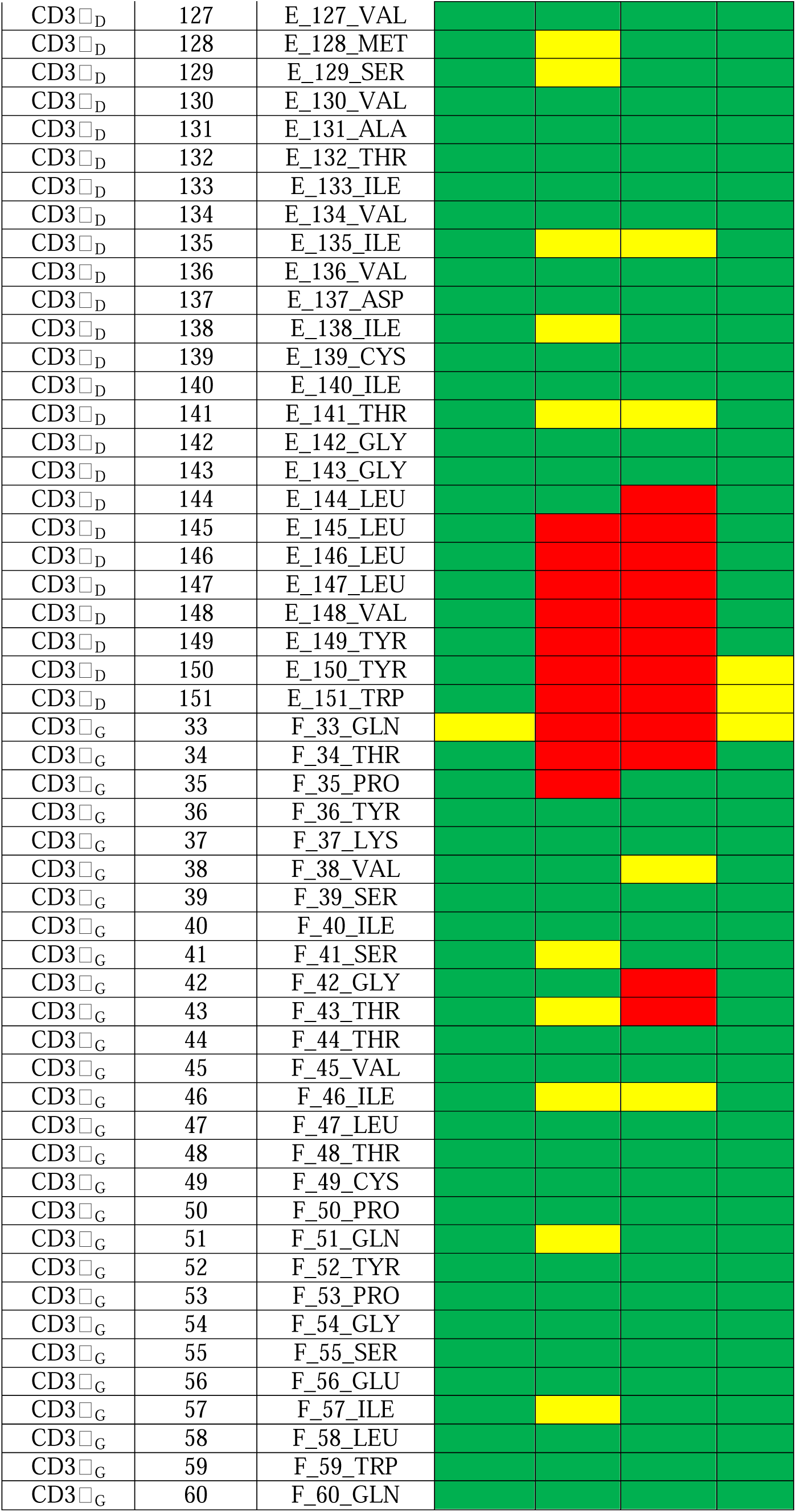
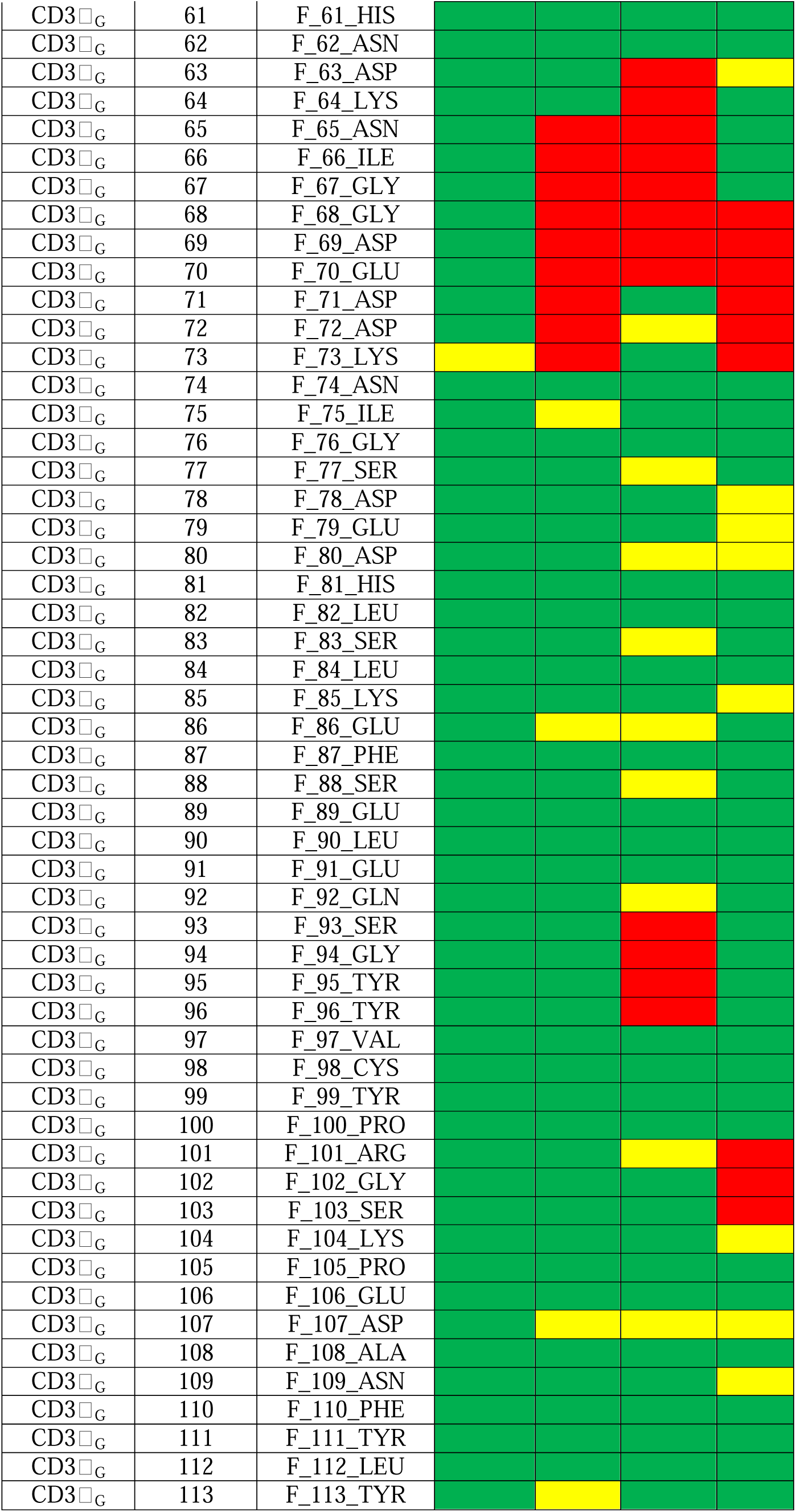
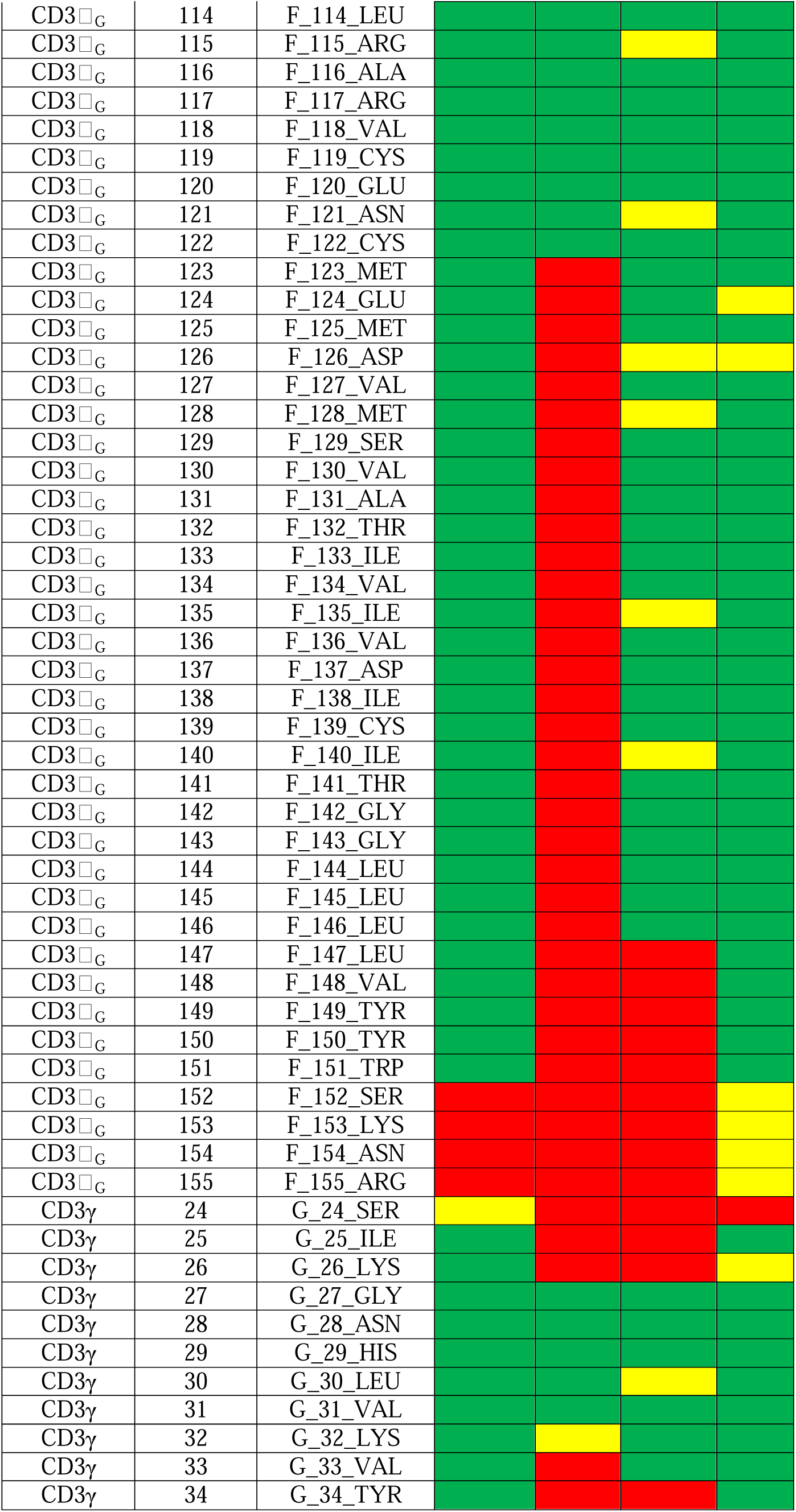
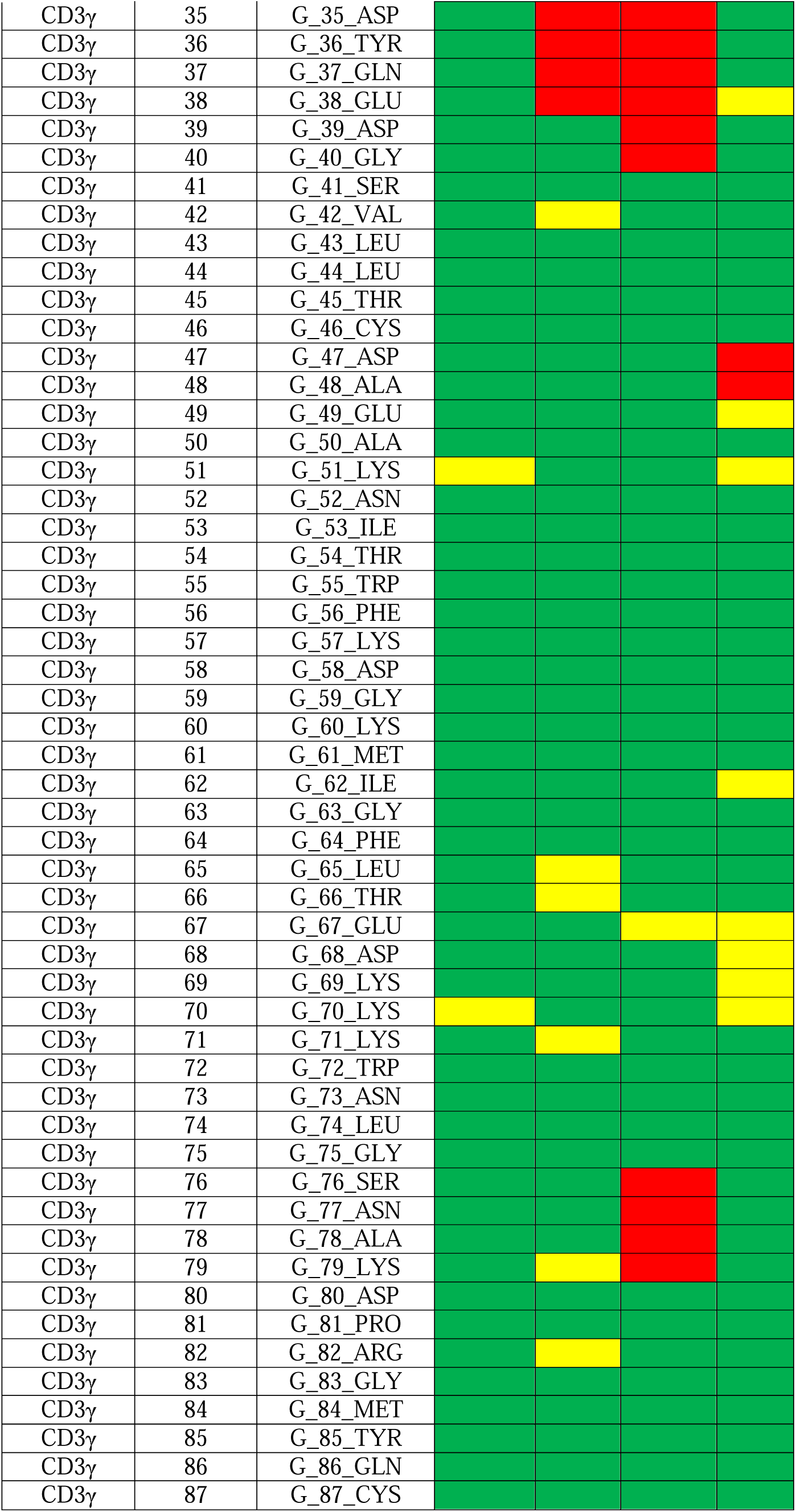
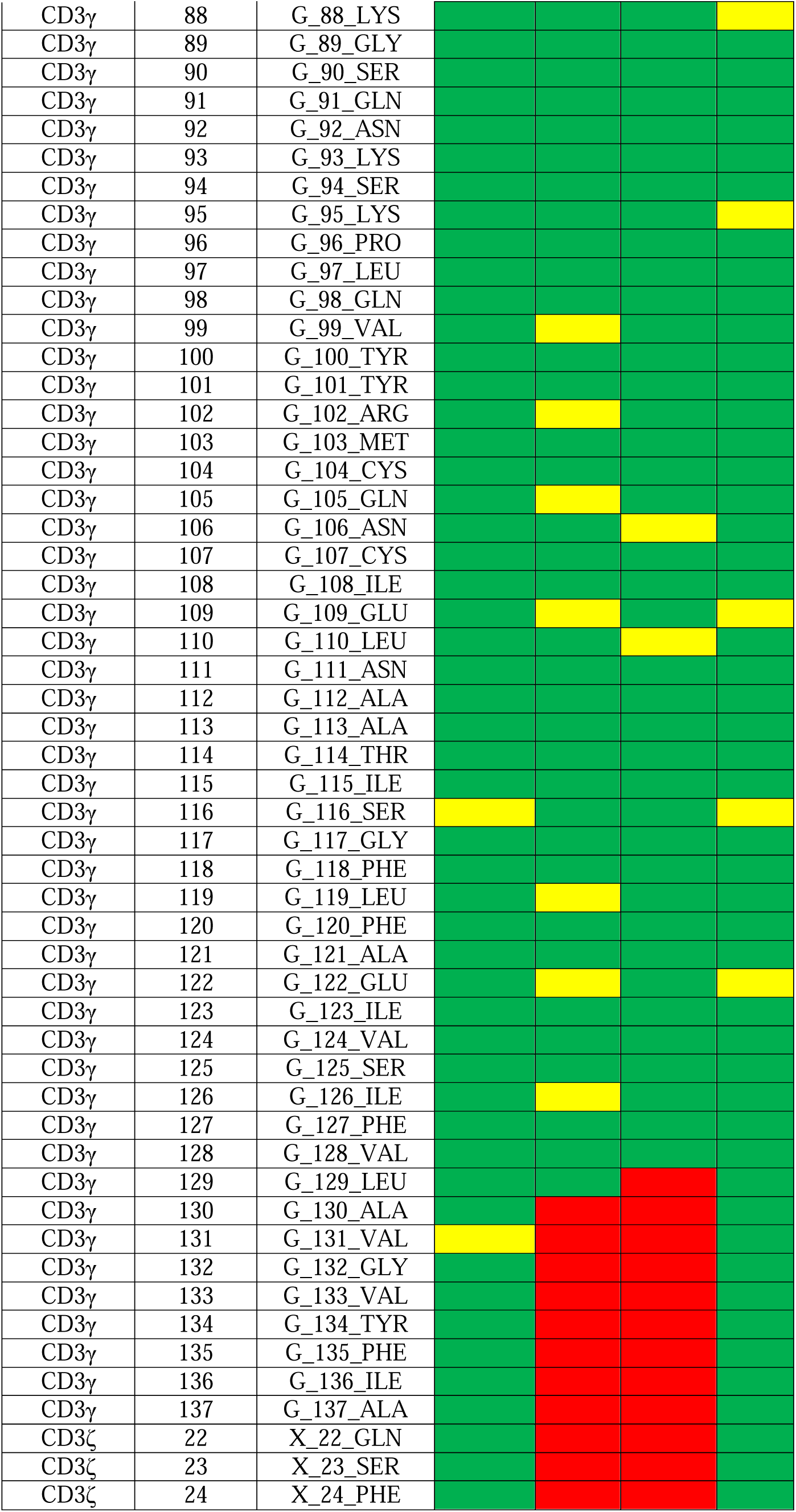
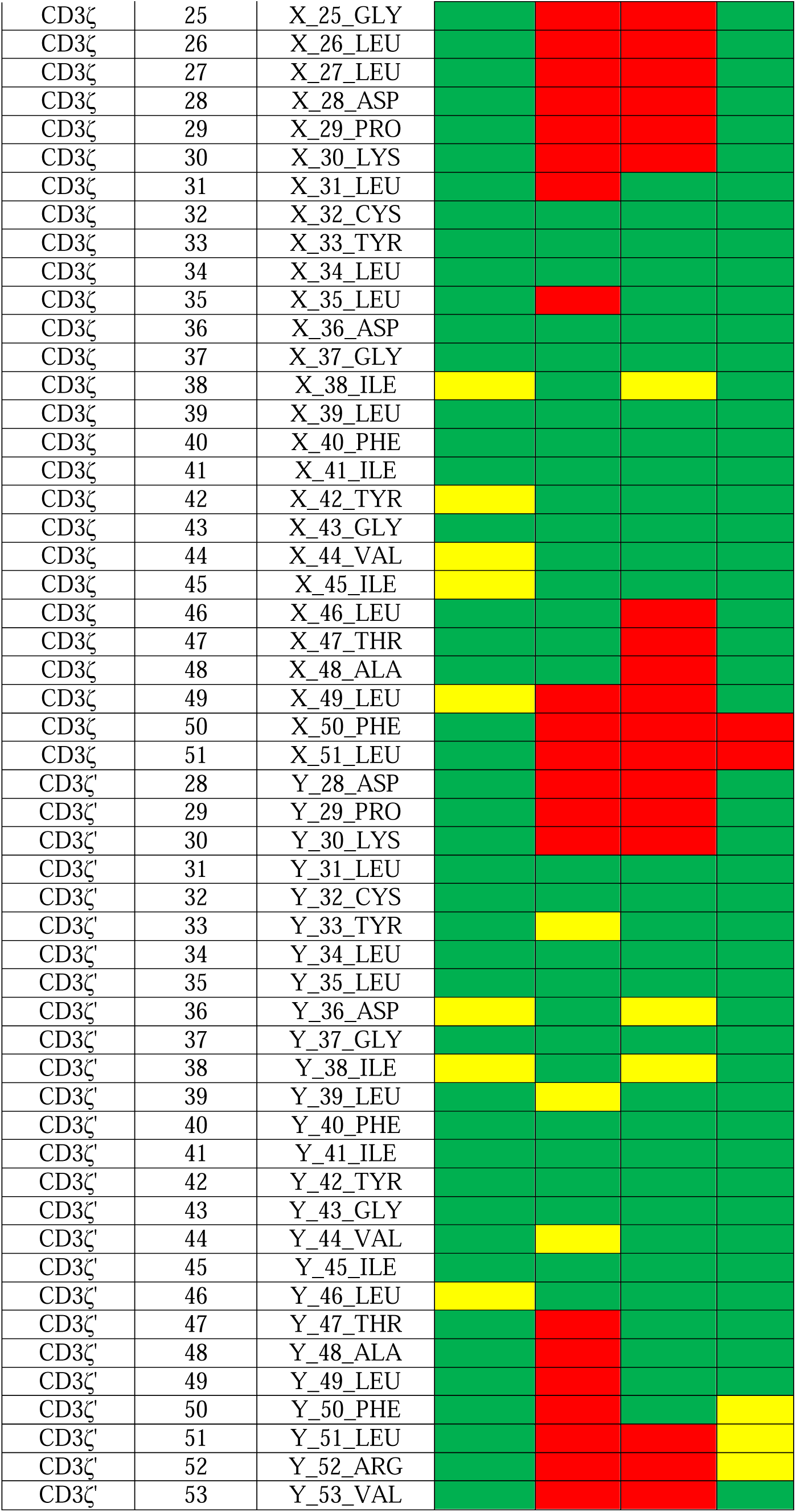
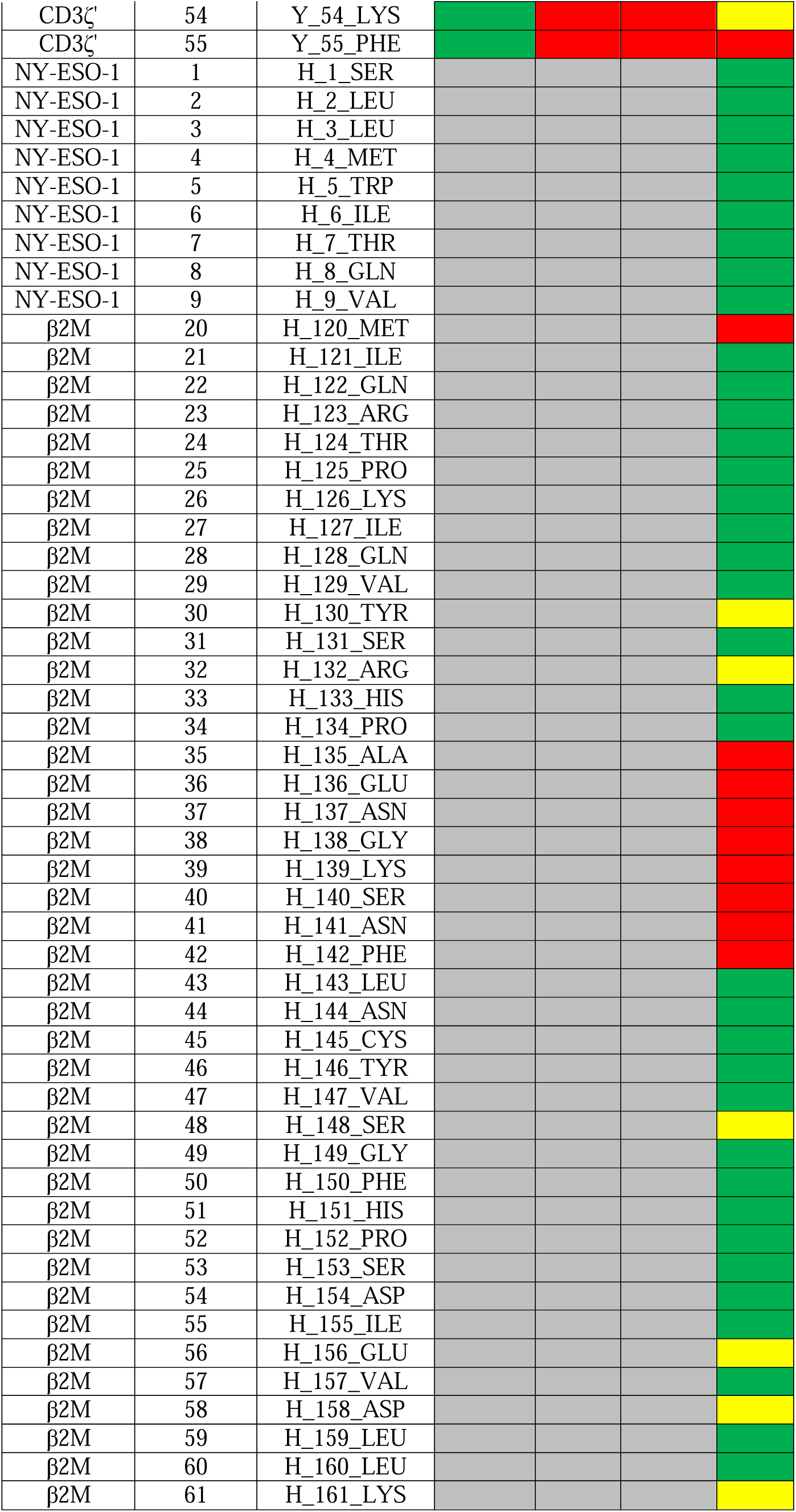
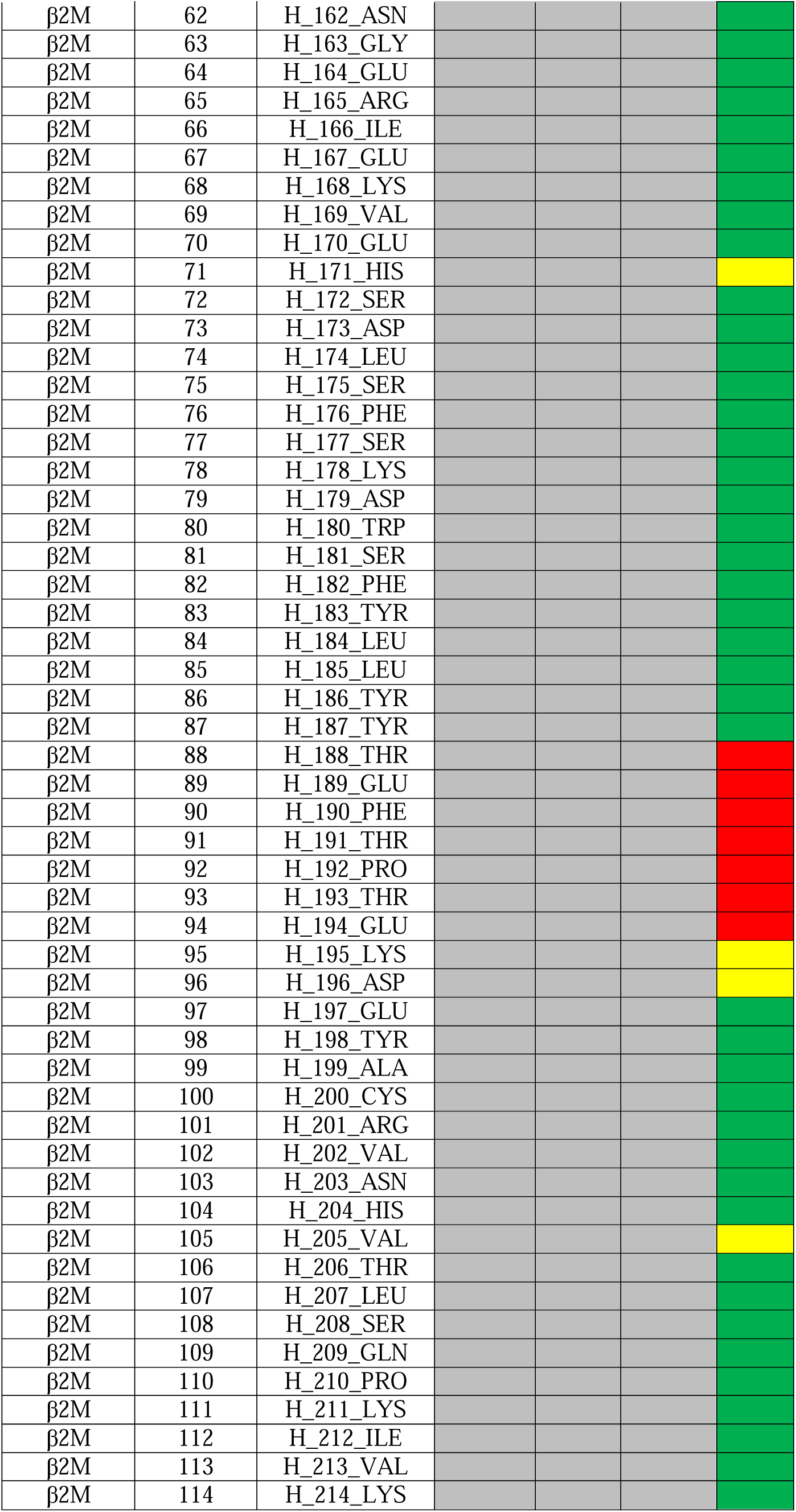
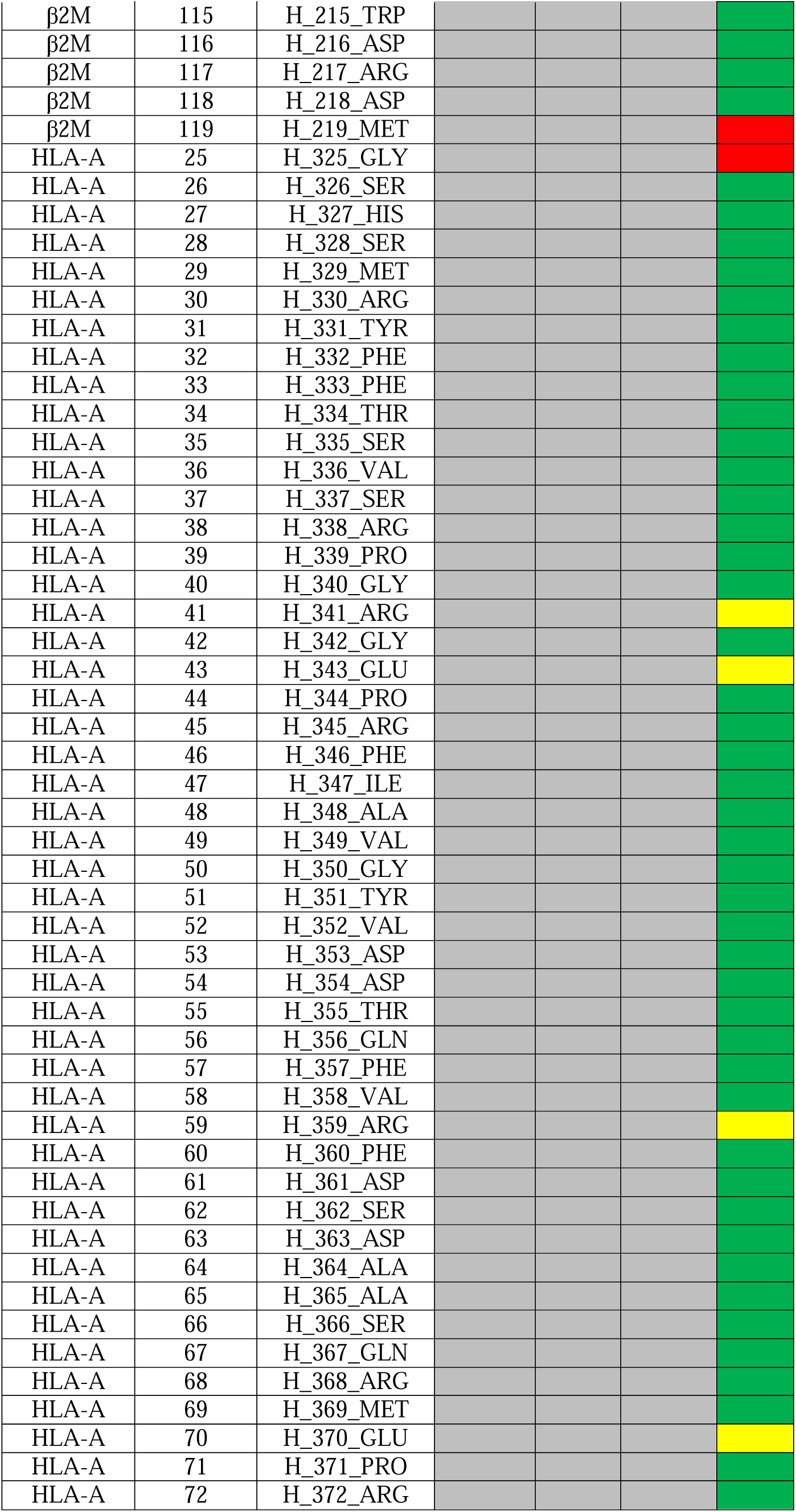
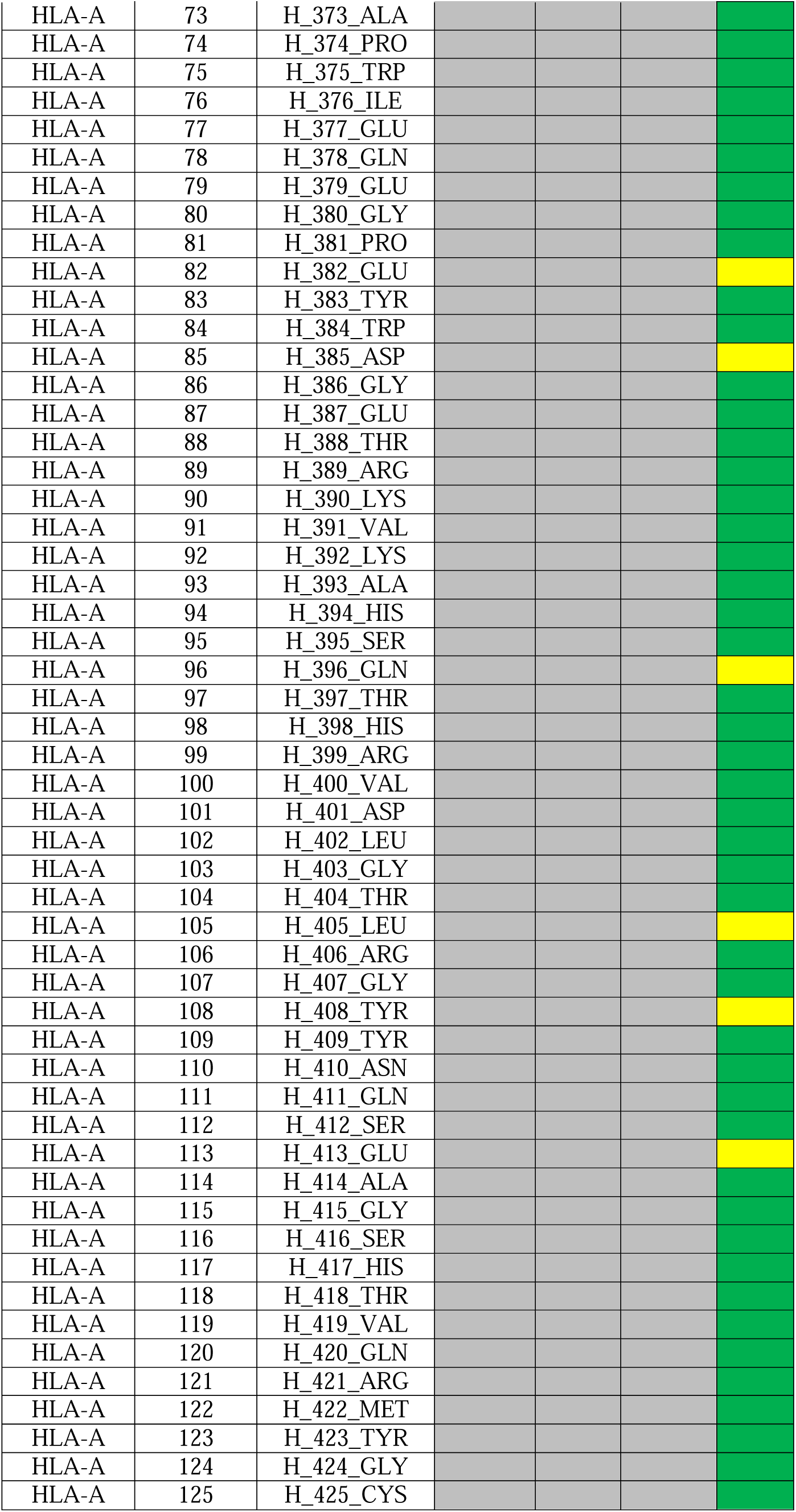
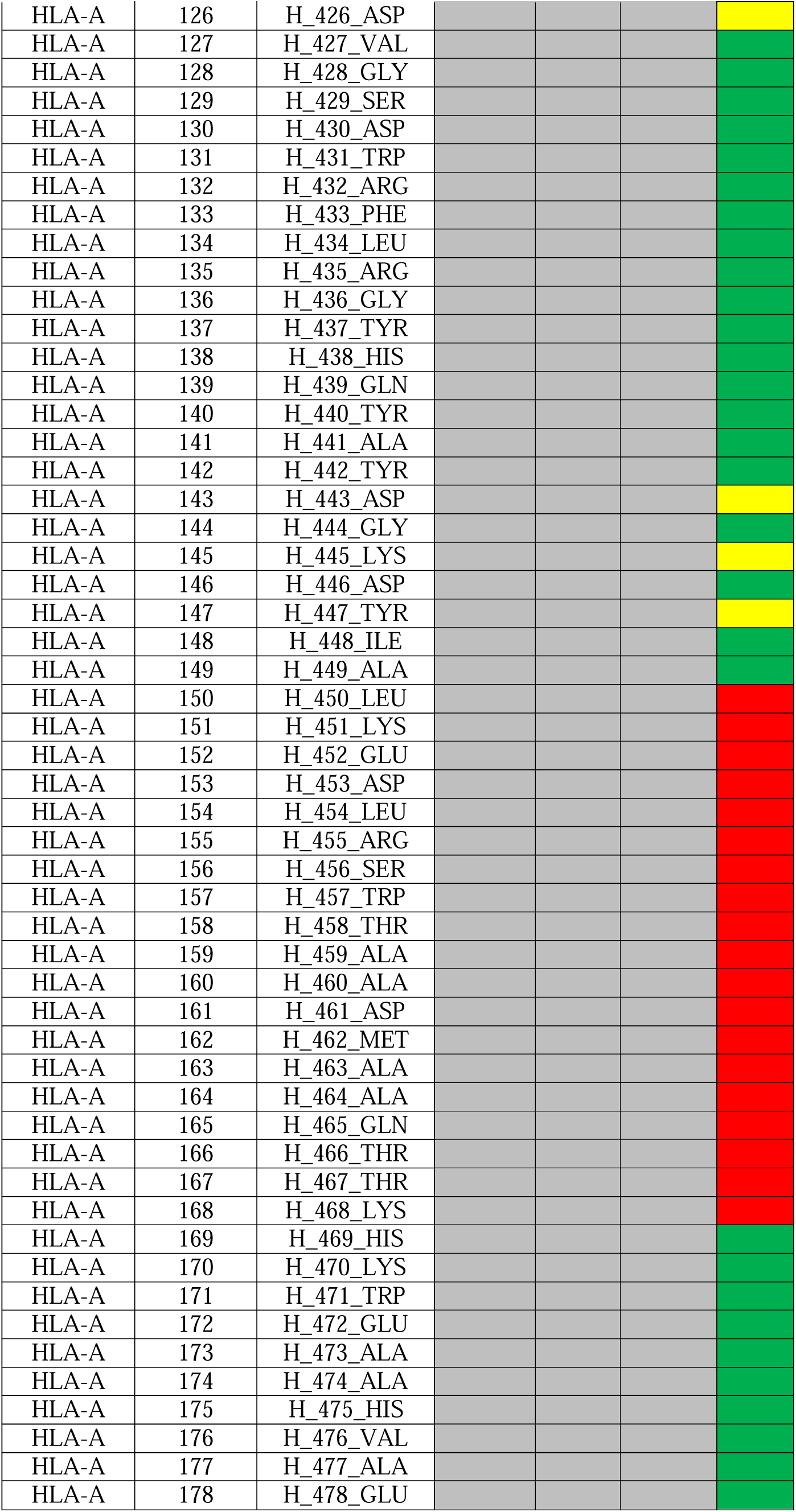
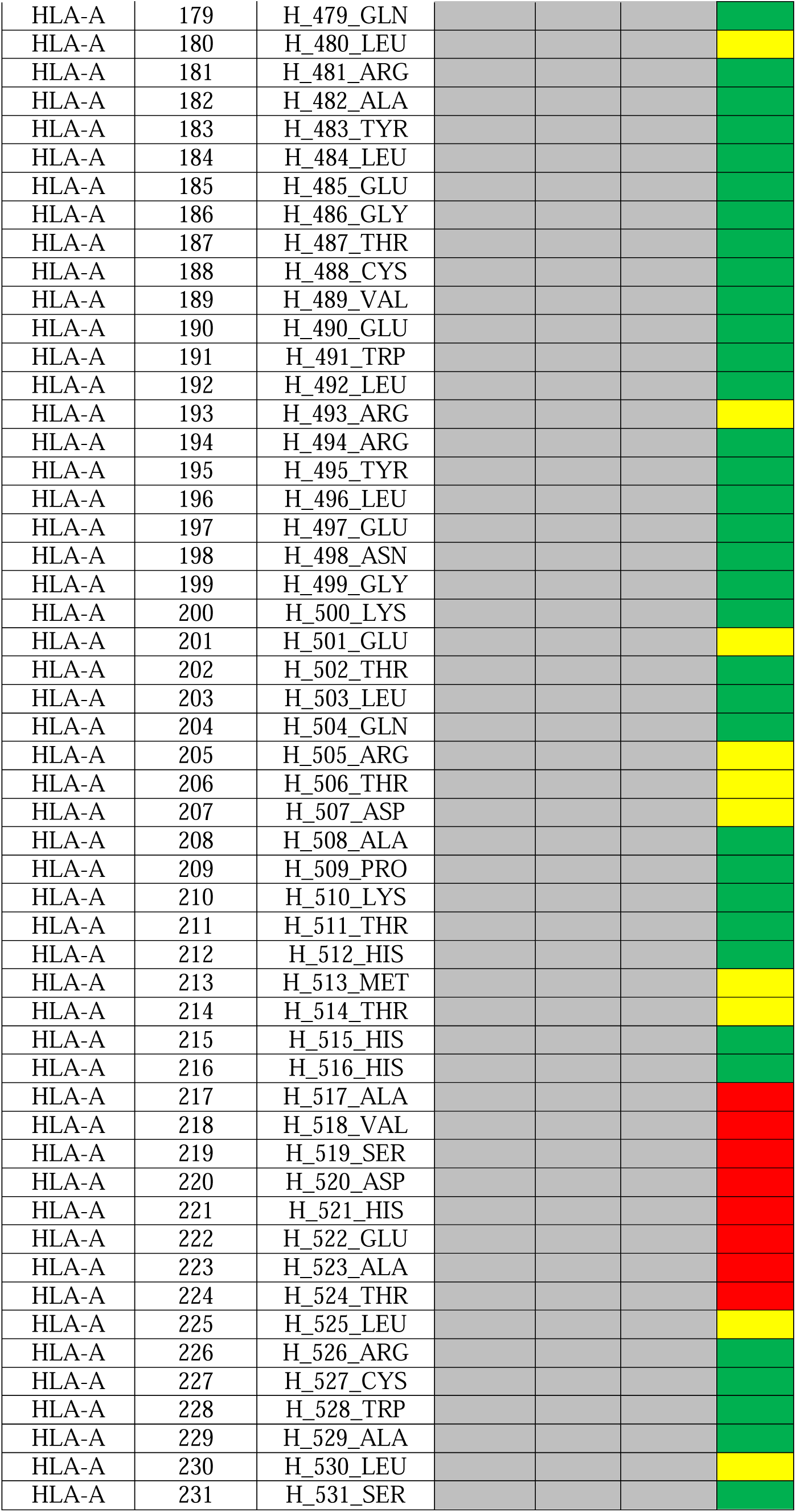
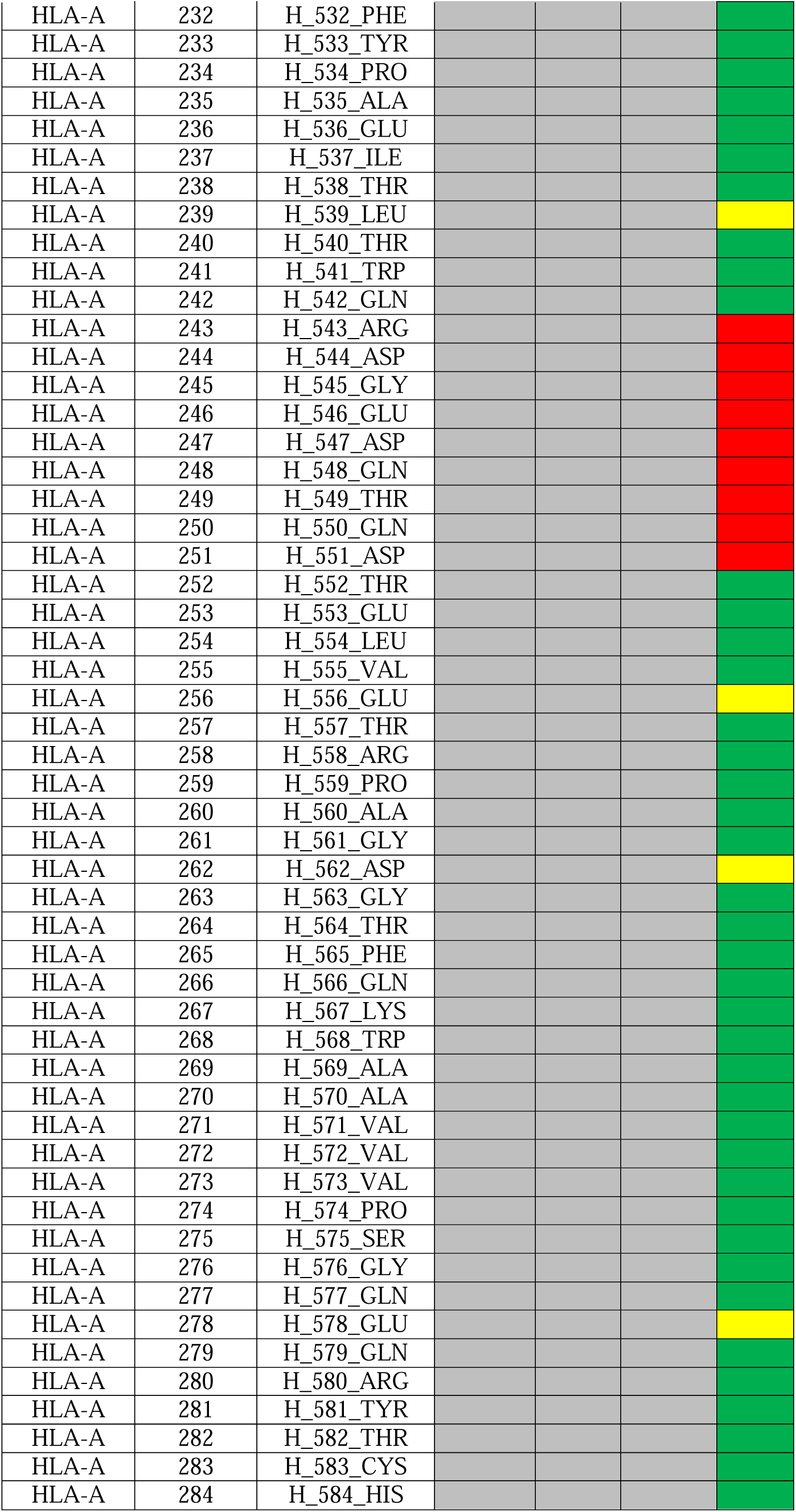
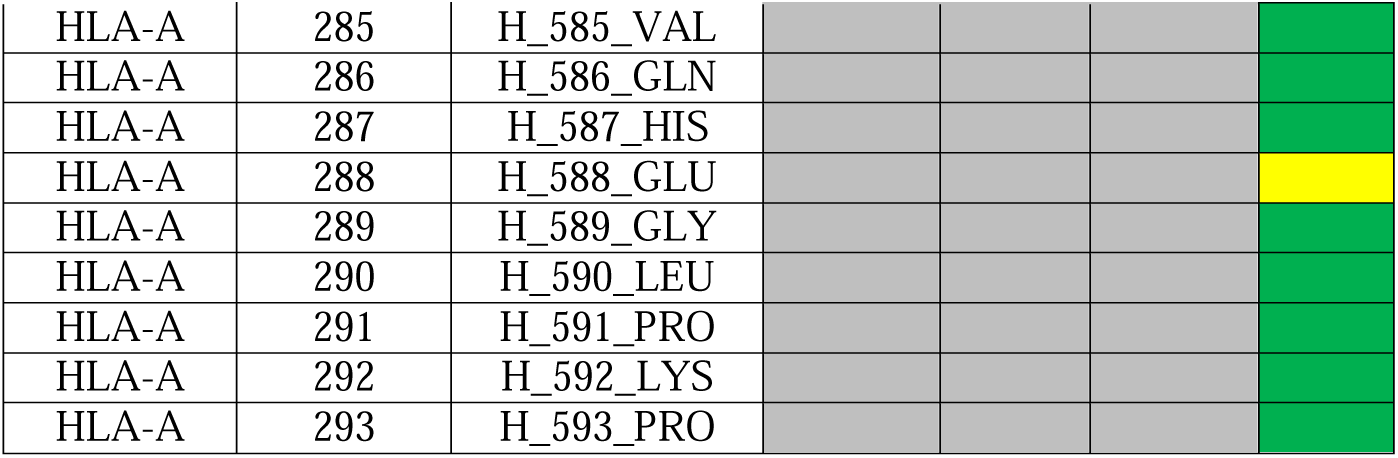
Model completeness for TCR–CD3 complexes. Green indicates a fully modeled residue, yellow indicates a stub amino acid, and red indicates residue not built. ID indicates the chain, residue number, and amino acid for a given residue in the structure. HLA-bound is abbreviated “HLA.”

**Table S4:**
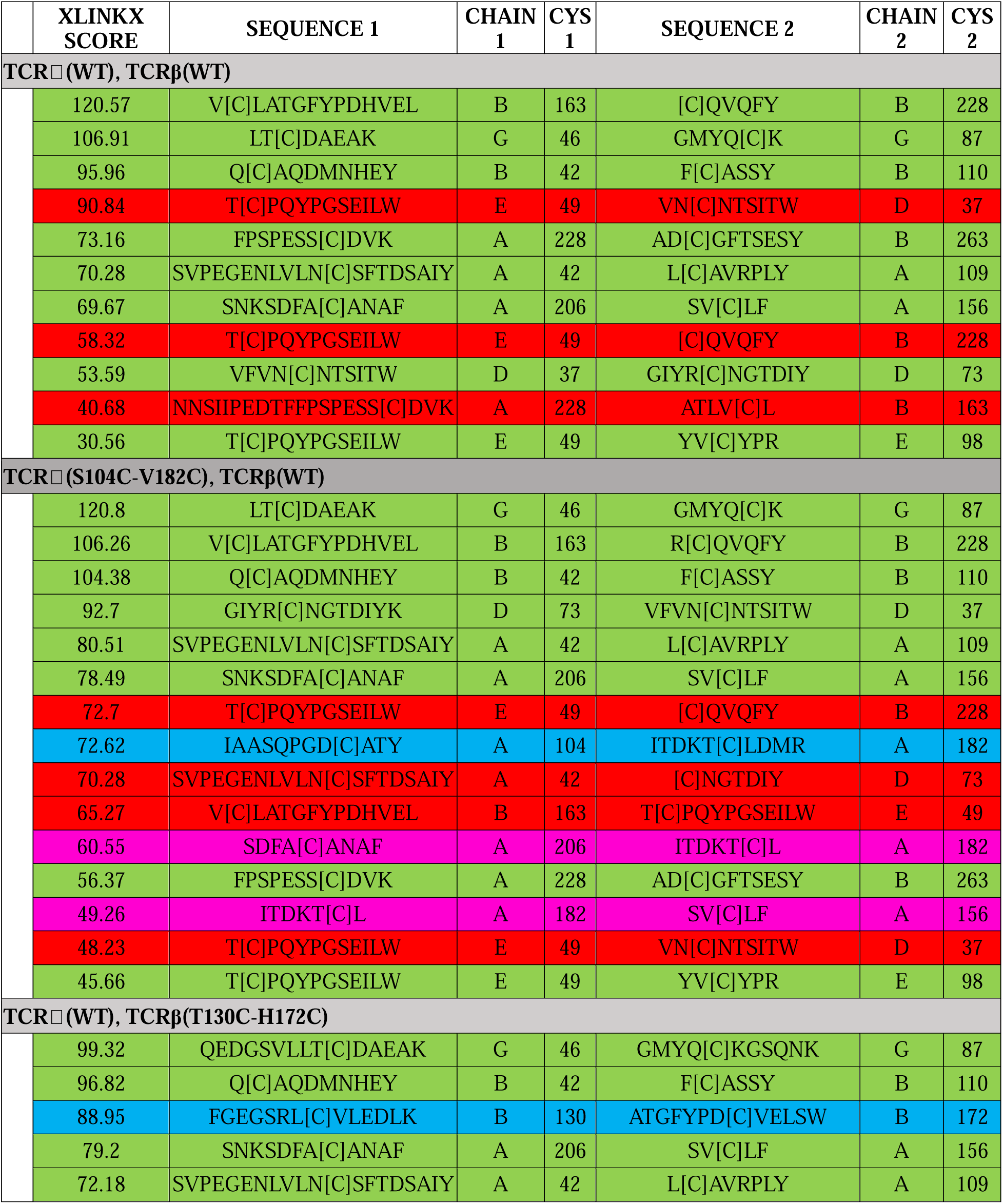
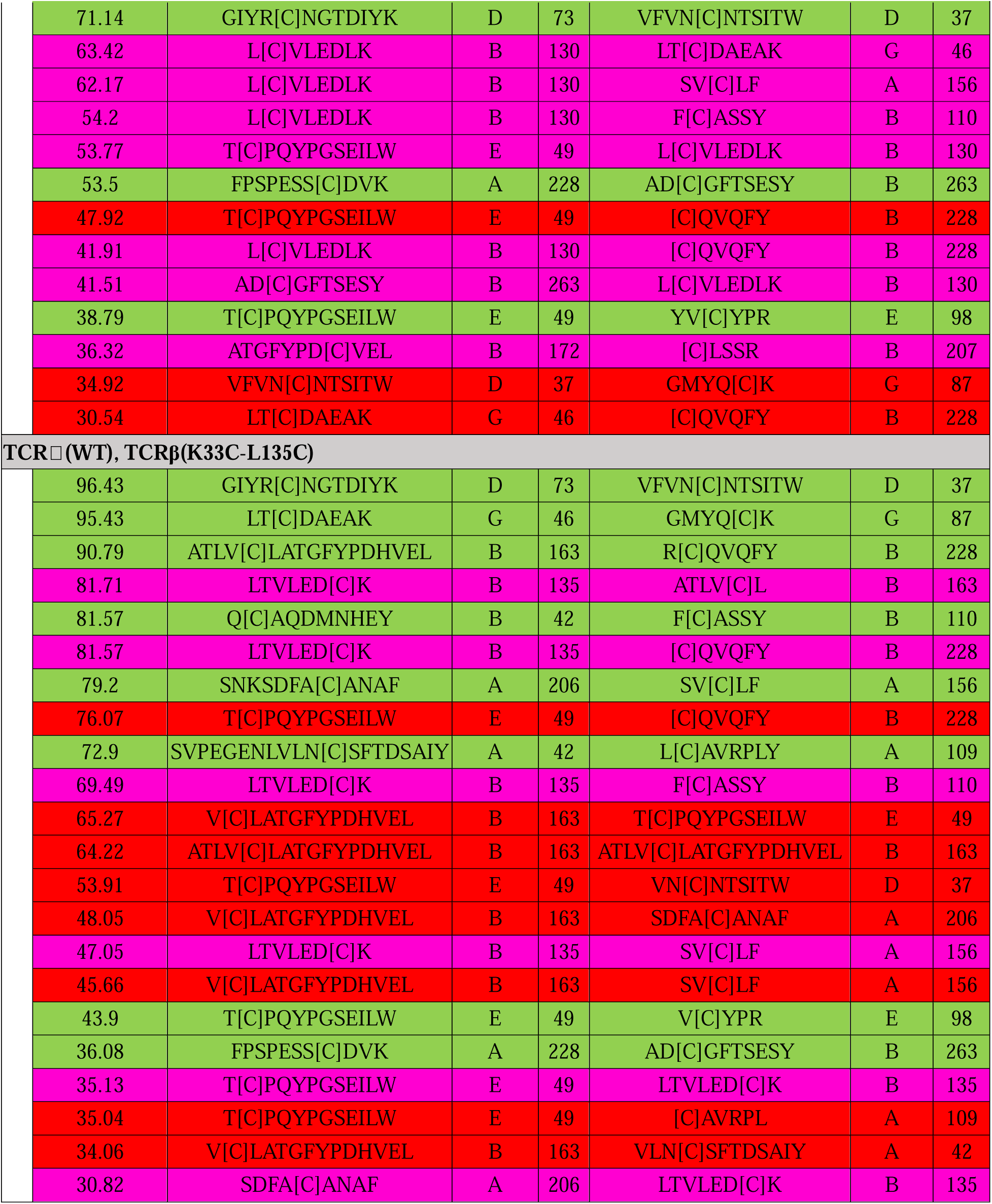
Disulfide-linked peptides identified by cross-linking mass spectrometry. Disulfide-linked peptides identified for each TCR genotype. Peptides with an XLinkX score greater than 30 are shown, as no native peptides were observed below this threshold. Color coding as follows: green, expected native disulfide; cyan, expected engineered disulfide; red, non-native disulfide or contaminant; magenta, off-target disulfide involving the engineered cysteines.

**Table S5:** Raw data for T-cell activation fluorescence reporter assay.

|  |  |  |
| --- | --- | --- |
| HLA: 8 micrograms/milliliter (peak t=16h) |  |  |
|  | WT | 104C/182C |
| n=1 | 164187.2 | 111629.3 |
| n=2 | 137446.8 | 114856.9 |
| n=3 | 167372.2 | 121945.9 |
| mean | 156335.4 | 116144.0 |
| % of WT | 100.0 | 74.3 |
| *P*-value |  | 0.0157 |
| HLA: 4 micrograms/milliliter (t=16h) |  |  |
|  | WT | 104C/182C |
| n=1 | 136138.2 | 113494.2 |
| n=2 | 128882.8 | 103848.4 |
| n=3 | 131584.7 | 119884.9 |
| mean | 132201.9 | 112409.2 |
| % of WT | 100.0 | 85.0 |
| *P*-value |  | 0.0181 |
| HLA: 0 micrograms/milliliter (t=16h) |  |  |
|  | WT | 104C/182C |
| n=1 | 424.8 | 402.0 |
| n=2 | 548.6 | 595.2 |
| n=3 | 794.2 | 765.6 |
| mean | 589.2 | 587.6 |
| % of WT | 100.0 | 99.7 |
| *P*-value |  | 0.9921 |
| PMA/Ionomycin (peak t = 10h) |  |  |
|  | WT | 104C/182C |
| n=1 | 178371.0 | 183172.8 |
| n=2 | 191040.4 | 175570.0 |
| n=3 | 184198.0 | 197041.9 |
| mean | 184536.5 | 185261.6 |
| % of WT | 100.0 | 100.4 |
| *P*-value |  | 0.9254 |

**Table S6:**
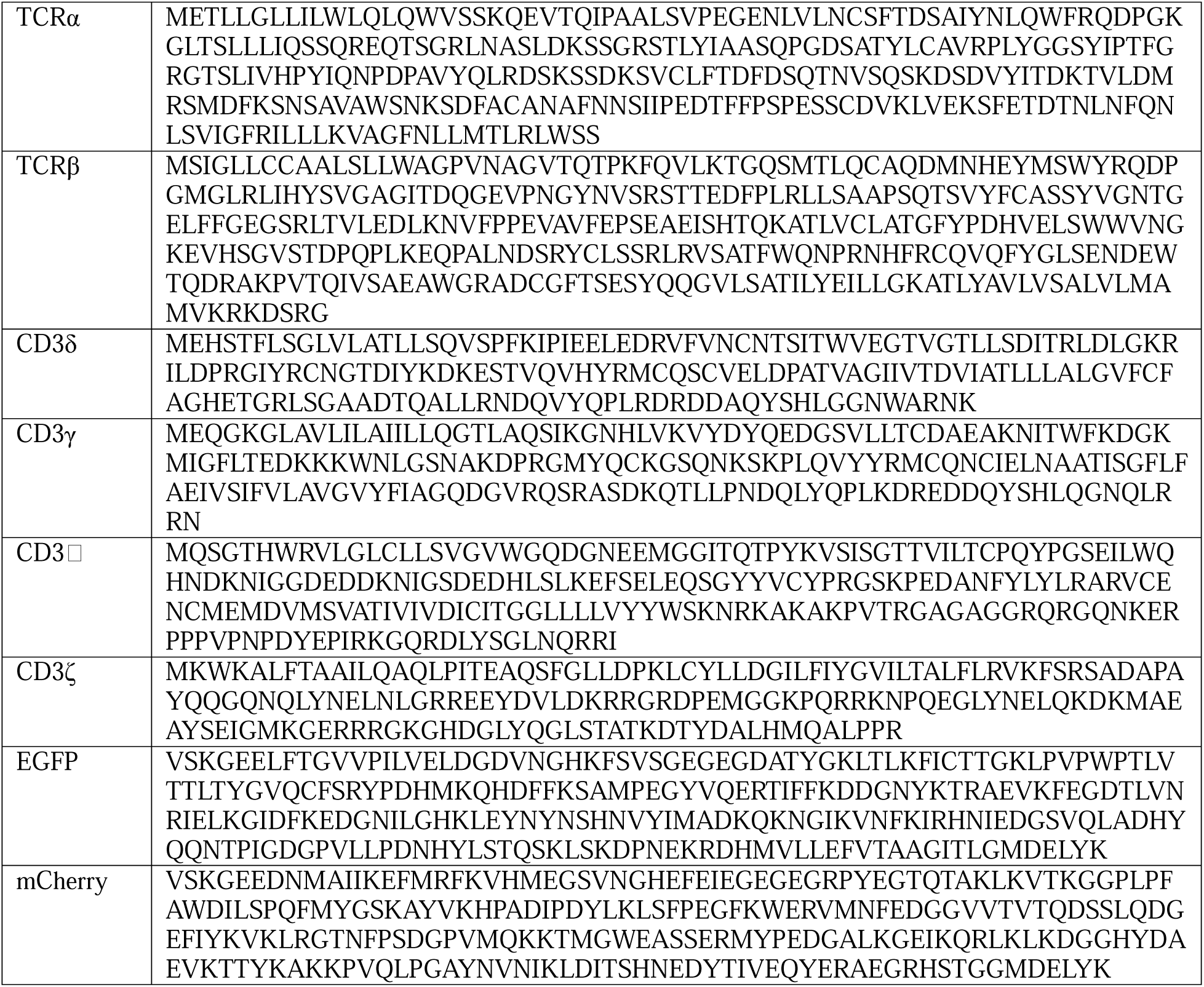
Wild-type protein sequences used in this study.

**Table S7:**
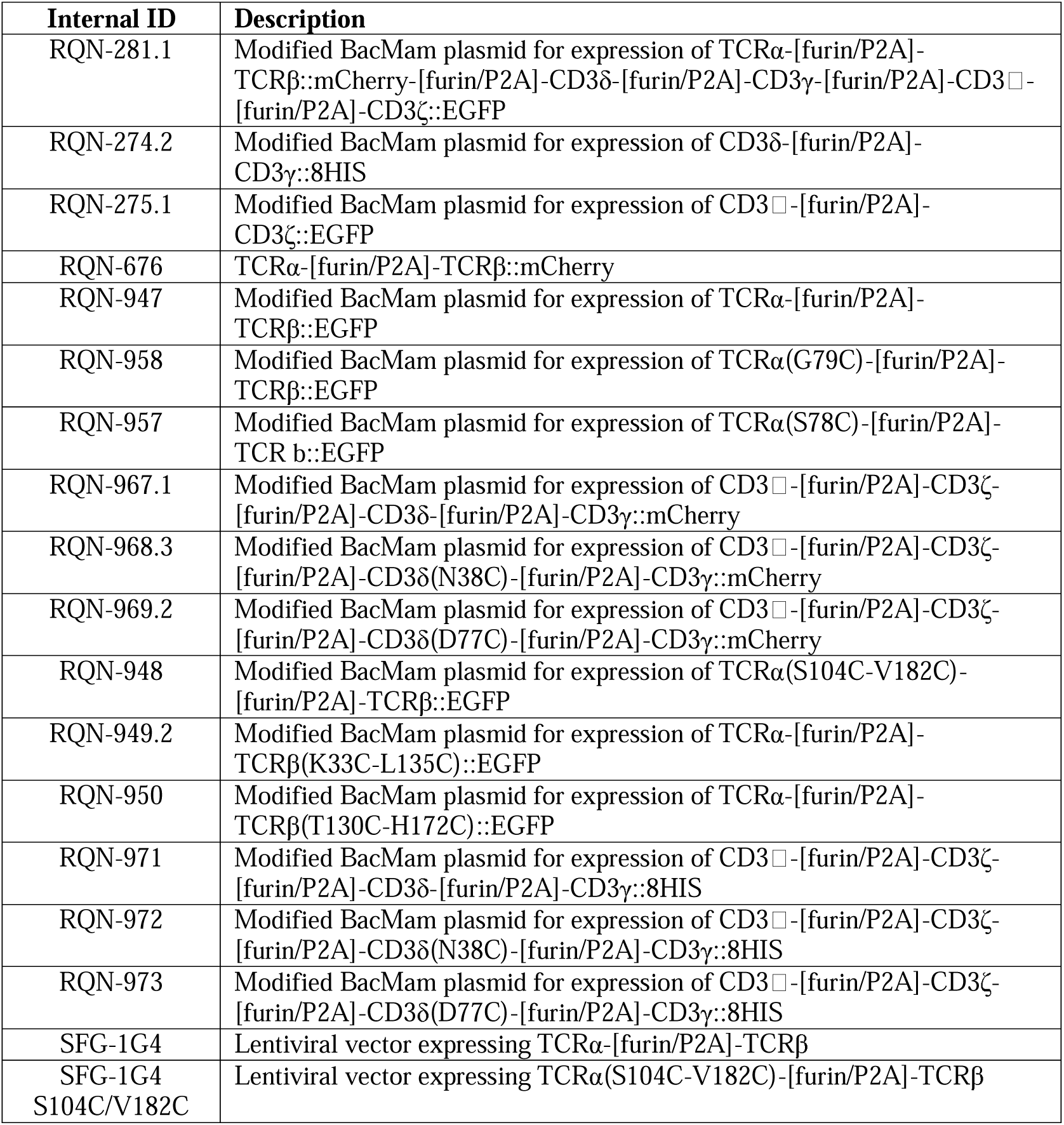
Plasmids used in this study.

**Table S8:** Antibodies used in this study.

| Antigen | Host | Label | Vendor | Catalog Number |
| --- | --- | --- | --- | --- |
| GFP | Mouse | n/a | Roche | 11814460001 |
| TCR Vβ13.1 | Mouse | Phycoerythrin | Biolegend | 362410 |
| Human CD3 □ | Mouse | Brilliant<br>Violet 785 | Biolegend | 317330 |
| Mouse IgG | Goat | Alexa700 | Invitrogen | A21036 |
| Rabbit IgG | Goat | Alexa790 | Invitrogen | A11369 |

**Movie S1: Conformational dynamics of the TCR–CD3 complex in the resting state.** Comparison of the protein constituents of the ND-I and ND-II models shown as cartoon representations and/or model surfaces. Transitions between conformations were created using the morph command in ChimeraX. The most prominent ectodomain movements are indicated by captions during transitions. Subunits colored as indicated in the legend at right.

**Movie S2: HLA-binding induces conformational change in the TCR–CD3 complex.** ChimeraX morph illustrating the transition from a resting conformation (ND-II, with HLA modeled) to the open/extended HLA-bound conformation. During the transition, the ectodomains open, the JM linkers extend, and the angle between the transmembrane helices and the membrane normal decreases. The latter is highlighted in a closeup view of the TMs towards the end of the movie. Models are shown as cartoon representations and/or model surfaces with subunits colored as indicated in the legends at right.

## References

1. C. A. Janeway, P. Travers, M. Walport, M. J. Shlomchik, C. A. J. Jr, P. Travers, M. Walport, M. J. Shlomchik, Immunobiology (Garland Science, ed. 5th, 2001).

2. E. L. Reinherz, The structure of a T-cell mechanosensor. Nature 573, 502–504 (2019).

3. A. H. Courtney, W.-L. Lo, A. Weiss, TCR Signaling: Mechanisms of Initiation and Propagation. Trends Biochem. Sci. 43, 108–123 (2018).

4. R. A. Mariuzza, P. Agnihotri, J. Orban, The structural basis of T-cell receptor (TCR) activation: An enduring enigma. J. Biol. Chem. 295, 914–925 (2020).

5. D. Dong, L. Zheng, J. Lin, B. Zhang, Y. Zhu, N. Li, S. Xie, Y. Wang, N. Gao, Z. Huang, Structural basis of assembly of the human T cell receptor–CD3 complex. Nature 573, 546– 552 (2019).

6. Y. Chen, Y. Zhu, X. Li, W. Gao, Z. Zhen, D. Dong, B. Huang, Z. Ma, A. Zhang, X. Song, Y. Ma, C. Guo, F. Zhang, Z. Huang, Cholesterol inhibits TCR signaling by directly restricting TCR-CD3 core tunnel motility. Mol. Cell 82, 1278–1287.e5 (2022).

7. L. Sušac, M. T. Vuong, C. Thomas, S. von Bülow, C. O’Brien-Ball, A. M. Santos, R. A. Fernandes, G. Hummer, R. Tampé, S. J. Davis, Structure of a fully assembled tumor-specific T cell receptor ligated by pMHC. Cell 185, 3201–3213.e19 (2022).

8. K. Saotome, D. Dudgeon, K. Colotti, M. J. Moore, J. Jones, Y. Zhou, A. Rafique, G. D. Yancopoulos, A. J. Murphy, J. C. Lin, W. C. Olson, M. C. Franklin, Structural analysis of cancer-relevant TCR-CD3 and peptide-MHC complexes by cryoEM. Nat. Commun. 14, 2401 (2023).

9. C. Xu, E. Gagnon, M. E. Call, J. R. Schnell, C. D. Schwieters, C. V. Carman, J. J. Chou, K. W. Wucherpfennig, Regulation of T Cell Receptor Activation by Dynamic Membrane Binding of the CD3 Cytoplasmic Tyrosine-Based Motif. Cell 135, 702–713 (2008).

10. H. Zhang, S.-P. Cordoba, O. Dushek, P. Anton van der Merwe, Basic residues in the T-cell receptor ζ cytoplasmic domain mediate membrane association and modulate signaling. Proc. Natl. Acad. Sci. 108, 19323–19328 (2011).

11. J. R. James, R. D. Vale, Biophysical Mechanism of T Cell Receptor Triggering in a Reconstituted System. Nature 487, 64–69 (2012).

12. S. Minguet, M. Swamy, B. Alarcón, I. F. Luescher, W. W. A. Schamel, Full activation of the T cell receptor requires both clustering and conformational changes at CD3. Immunity 26, 43–54 (2007).

13. D. Gil, W. W. A. Schamel, M. Montoya, F. Sánchez-Madrid, B. Alarcón, Recruitment of Nck by CD3 epsilon reveals a ligand-induced conformational change essential for T cell receptor signaling and synapse formation. Cell 109, 901–912 (2002).

14. K. Natarajan, A. C. McShan, J. Jiang, V. K. Kumirov, R. Wang, H. Zhao, P. Schuck, M. E. Tilahun, L. F. Boyd, J. Ying, A. Bax, D. H. Margulies, N. G. Sgourakis, An allosteric site in the T-cell receptor Cβ domain plays a critical signalling role. Nat. Commun. 8, 15260 (2017).

15. S. Rangarajan, Y. He, Y. Chen, M. C. Kerzic, B. Ma, R. Gowthaman, B. G. Pierce, R. Nussinov, R. A. Mariuzza, J. Orban, Peptide-MHC (pMHC) binding to a human antiviral T cell receptor induces long-range allosteric communication between pMHC- and CD3-binding sites. J. Biol. Chem. 293, 15991–16005 (2018).

16. D. K. Das, Y. Feng, R. J. Mallis, X. Li, D. B. Keskin, R. E. Hussey, S. K. Brady, J.-H. Wang, G. Wagner, E. L. Reinherz, M. J. Lang, Force-dependent transition in the T-cell receptor β-subunit allosterically regulates peptide discrimination and pMHC bond lifetime. Proc. Natl. Acad. Sci. 112, 1517–1522 (2015).

17. W. Wu, X. Shi, C. Xu, Regulation of T cell signalling by membrane lipids. Nat. Rev. Immunol. 16, 690–701 (2016).

18. F. Wang, K. Beck-García, C. Zorzin, W. W. A. Schamel, M. M. Davis, Inhibition of T cell receptor signaling by cholesterol sulfate, a naturally occurring derivative of membrane cholesterol. Nat. Immunol. 17, 844–850 (2016).

19. R. Q. Notti, T. Walz, Native-like environments afford novel mechanistic insights into membrane proteins. Trends Biochem. Sci. (2022).

20. P. F. Robbins, S. H. Kassim, T. L. N. Tran, J. S. Crystal, R. A. Morgan, S. A. Feldman, J. C. Yang, M. E. Dudley, J. R. Wunderlich, R. M. Sherry, U. S. Kammula, M. S. Hughes, N. P. Restifo, M. Raffeld, C.-C. R. Lee, Y. F. Li, M. El-Gamil, S. A. Rosenberg, A Pilot Trial Using Lymphocytes Genetically Engineered with an NY-ESO-1–Reactive T-cell Receptor: Long-term Follow-up and Correlates with Response. Clin. Cancer Res. 21, 1019–1027 (2015).

21. M. B. Ulmschneider, M. S. P. Sansom, Amino acid distributions in integral membrane protein structures. Biochim. Biophys. Acta BBA - Biomembr. 1512, 1–14 (2001).

22. M. Swamy, K. Beck-Garcia, E. Beck-Garcia, F. A. Hartl, A. Morath, O. S. Yousefi, E. P. Dopfer, E. Molnár, A. K. Schulze, R. Blanco, A. Borroto, N. Martín-Blanco, B. Alarcon, T. Höfer, S. Minguet, W. W. A. Schamel, A Cholesterol-Based Allostery Model of T Cell Receptor Phosphorylation. Immunity 44, 1091–1101 (2016).

23. Q. Su, M. Chen, Y. Shi, X. Zhang, G. Huang, B. Huang, D. Liu, Z. Liu, Y. Shi, Cryo-EM structure of the human IgM B cell receptor. Science 377, 875–880 (2022).

24. X. Gao, X. Dong, X. Li, Z. Liu, H. Liu, Prediction of disulfide bond engineering sites using a machine learning method. Sci. Rep. 10, 10330 (2020).

25. F. Liu, B. van Breukelen, A. J. R. Heck, Facilitating Protein Disulfide Mapping by a Combination of Pepsin Digestion, Electron Transfer Higher Energy Dissociation (EThcD), and a Dedicated Search Algorithm SlinkS*. Mol. Cell. Proteomics 13, 2776–2786 (2014).

26. G. B. E. Stewart-Jones, C. Soto, T. Lemmin, G.-Y. Chuang, A. Druz, R. Kong, P. V. Thomas, K. Wagh, T. Zhou, A.-J. Behrens, T. Bylund, C. W. Choi, J. R. Davison, I. S. Georgiev, M. G. Joyce, Y. D. Kwon, M. Pancera, J. Taft, Y. Yang, B. Zhang, S. S. Shivatare, V. S. Shivatare, C.-C. D. Lee, C.-Y. Wu, C. A. Bewley, D. R. Burton, W. C. Koff, M. Connors, M. Crispin, U. Baxa, B. T. Korber, C.-H. Wong, J. R. Mascola, P. D. Kwong, Trimeric HIV-1-Env Structures Define Glycan Shields from Clades A, B, and G. Cell 165, 813–826 (2016).

27. Y. Yin, X. X. Wang, R. A. Mariuzza, Crystal structure of a complete ternary complex of T-cell receptor, peptide-MHC, and CD4. Proc. Natl. Acad. Sci. U. S. A. 109, 5405–5410 (2012).

28. W. F. Hawse, M. M. Champion, M. V. Joyce, L. M. Hellman, M. Hossain, V. Ryan, B. G. Pierce, Z. Weng, B. M. Baker, Cutting Edge: Evidence for a Dynamically Driven T Cell Signaling Mechanism. J. Immunol. 188, 5819–5823 (2012).

29. A. A.-L. Lanz, G. Masi, N. Porciello, A. Cohnen, D. Cipria, D. Prakaash, Š. Bálint, R. Raggiaschi, D. Galgano, D. K. Cole, M. Lepore, O. Dushek, M. L. Dustin, M. S. P. Sansom, C. Kalli, O. Acuto, Allosteric activation of T cell antigen receptor signaling by quaternary structure relaxation. Cell Rep. 36, 109375 (2021).

29. S. T. Kim, K. Takeuchi, Z.-Y. J. Sun, M. Touma, C. E. Castro, A. Fahmy, M. J. Lang, G. Wagner, E. L. Reinherz, The αβ T Cell Receptor Is an Anisotropic Mechanosensor *. J. Biol. Chem. 284, 31028–31037 (2009).

31. R. A. Fernandes, D. A. Shore, M. T. Vuong, C. Yu, X. Zhu, S. Pereira-Lopes, H. Brouwer, J. A. Fennelly, C. M. Jessup, E. J. Evans, I. A. Wilson, S. J. Davis, T Cell Receptors are Structures Capable of Initiating Signaling in the Absence of Large Conformational Rearrangements*. J. Biol. Chem. 287, 13324–13335 (2012).

32. J. H. Lee, N. de Val, D. Lyumkis, A. B. Ward, Model Building and Refinement of a Natively Glycosylated HIV-1 Env Protein by High-Resolution Cryoelectron Microscopy. Structure 23, 1943–1951 (2015).

33. N. Chouaki Benmansour, K. Ruminski, A.-M. Sartre, M.-C. Phelipot, A. Salles, E. Bergot, A. Wu, G. Chicanne, M. Fallet, S. Brustlein, C. Billaudeau, A. Formisano, S. Mailfert, B. Payrastre, D. Marguet, S. Brasselet, Y. Hamon, H.-T. He, Phosphoinositides regulate the TCR/CD3 complex membrane dynamics and activation. Sci. Rep. 8, 4966 (2018).

34. P. S. Costello, M. Gallagher, D. A. Cantrell, Sustained and dynamic inositol lipid metabolism inside and outside the immunological synapse. Nat. Immunol. 3, 1082–1089 (2002).

35. S. Xu, O. Chaudhary, P. Rodríguez-Morales, X. Sun, D. Chen, R. Zappasodi, Z. Xu, A. F. M. Pinto, A. Williams, I. Schulze, Y. Farsakoglu, S. K. Varanasi, J. S. Low, W. Tang, H. Wang, B. McDonald, V. Tripple, M. Downes, R. M. Evans, N. A. Abumrad, T. Merghoub, J. D. Wolchok, M. N. Shokhirev, P.-C. Ho, J. L. Witztum, B. Emu, G. Cui, S. M. Kaech, Uptake of oxidized lipids by the scavenger receptor CD36 promotes lipid peroxidation and dysfunction in CD8+ T cells in tumors. Immunity 54, 1561–1577.e7 (2021).

36. A. Kent, N. V. Longino, A. Christians, E. Davila, Naturally Occurring Genetic Alterations in Proximal TCR Signaling and Implications for Cancer Immunotherapy. Front. Immunol. 12 (2021).

37. C. A. Klebanoff, S. S. Chandran, B. M. Baker, S. A. Quezada, A. Ribas, T cell receptor therapeutics: immunological targeting of the intracellular cancer proteome. Nat. Rev. Drug Discov. 22, 996–1017 (2023).

38. A. Goehring, C.-H. Lee, K. H. Wang, J. C. Michel, D. P. Claxton, I. Baconguis, T. Althoff, S. Fischer, K. C. Garcia, E. Gouaux, Screening and large-scale expression of membrane proteins in mammalian cells for structural studies. Nat. Protoc. 9, 2574–2585 (2014).

39. Y. Zhang, C. Daday, R.-X. Gu, C. D. Cox, B. Martinac, B. L. de Groot, T. Walz, Visualization of the mechanosensitive ion channel MscS under membrane tension. Nature 590, 509–514 (2021).

40. M. Ohi, Y. Li, Y. Cheng, T. Walz, Negative Staining and Image Classification – Powerful Tools in Modern Electron Microscopy. Biol. Proced. Online 6, 23–34 (2004).

41. D. N. Mastronarde, Automated electron microscope tomography using robust prediction of specimen movements. J. Struct. Biol. 152, 36–51 (2005).

42. A. Cheng, E. T. Eng, L. Alink, W. J. Rice, K. D. Jordan, L. Y. Kim, C. S. Potter, B. Carragher, High resolution single particle cryo-electron microscopy using beam-image shift. J. Struct. Biol. 204, 270–275 (2018).

43. S. Q. Zheng, E. Palovcak, J.-P. Armache, K. A. Verba, Y. Cheng, D. A. Agard, MotionCor2: anisotropic correction of beam-induced motion for improved cryo-electron microscopy. Nat. Methods 14, 331–332 (2017).

44. A. Punjani, J. L. Rubinstein, D. J. Fleet, M. A. Brubaker, cryoSPARC: algorithms for rapid unsupervised cryo-EM structure determination. Nat. Methods 14, 290–296 (2017).

45. A. Rohou, N. Grigorieff, CTFFIND4: Fast and accurate defocus estimation from electron micrographs. J. Struct. Biol. 192, 216–221 (2015).

46. S. Jo, T. Kim, V. G. Iyer, W. Im, CHARMM-GUI: a web-based graphical user interface for CHARMM. J. Comput. Chem. 29, 1859–1865 (2008).

47. Y. Qi, J. Lee, J. B. Klauda, W. Im, CHARMM-GUI Nanodisc Builder for modeling and simulation of various nanodisc systems. J. Comput. Chem. 40, 893–899 (2019).

48. A. Punjani, D. J. Fleet, 3D variability analysis: Resolving continuous flexibility and discrete heterogeneity from single particle cryo-EM. J. Struct. Biol. 213, 107702 (2021).

49. A. Punjani, H. Zhang, D. J. Fleet, Non-uniform refinement: adaptive regularization improves single-particle cryo-EM reconstruction. Nat. Methods 17, 1214–1221 (2020).

50. R. Sanchez-Garcia, J. Gomez-Blanco, A. Cuervo, J. M. Carazo, C. O. S. Sorzano, J. Vargas, DeepEMhancer: a deep learning solution for cryo-EM volume post-processing. Commun. Biol. 4, 1–8 (2021).

51. J.-L. Chen, G. Stewart-Jones, G. Bossi, N. M. Lissin, L. Wooldridge, E. M. L. Choi, G. Held, P. R. Dunbar, R. M. Esnouf, M. Sami, J. M. Boulter, P. Rizkallah, C. Renner, A. Sewell, P. A. van der Merwe, B. K. Jakobsen, G. Griffiths, E. Y. Jones, V. Cerundolo, Structural and kinetic basis for heightened immunogenicity of T cell vaccines. J. Exp. Med. 201, 1243– 1255 (2005).

52. S. Aiyer, C. Zhang, P. R. Baldwin, D. Lyumkis, Evaluating Local and Directional Resolution of Cryo-EM Density Maps. Methods Mol. Biol. Clifton NJ 2215, 161–187 (2021).

53. T. D. Goddard, C. C. Huang, E. C. Meng, E. F. Pettersen, G. S. Couch, J. H. Morris, T. E. Ferrin, UCSF ChimeraX: Meeting modern challenges in visualization and analysis. Protein Sci. Publ. Protein Soc. 27, 14–25 (2018).

54. P. V. Afonine, B. K. Poon, R. J. Read, O. V. Sobolev, T. C. Terwilliger, A. Urzhumtsev, P. D. Adams, Real-space refinement in PHENIX for cryo-EM and crystallography. Acta Crystallogr. Sect. Struct. Biol. 74, 531–544 (2018).

55. A. Casañal, B. Lohkamp, P. Emsley, Current developments in Coot for macromolecular model building of Electron Cryo microscopy and Crystallographic Data. Protein Sci. Publ. Protein Soc. 29, 1069–1078 (2020).

56. T. I. Croll, ISOLDE: a physically realistic environment for model building into low-resolution electron-density maps. Acta Crystallogr. Sect. Struct. Biol. 74, 519–530 (2018).

57. P. Emsley, M. Crispin, Structural analysis of glycoproteins: building N-linked glycans with Coot. Acta Crystallogr. Sect. Struct. Biol. 74, 256–263 (2018).

58. H. Ashkenazy, S. Abadi, E. Martz, O. Chay, I. Mayrose, T. Pupko, N. Ben-Tal, ConSurf 2016: an improved methodology to estimate and visualize evolutionary conservation in macromolecules. Nucleic Acids Res. 44, W344–W350 (2016).

59. J. Rappsilber, Y. Ishihama, M. Mann, Stop and go extraction tips for matrix-assisted laser desorption/ionization, nanoelectrospray, and LC/MS sample pretreatment in proteomics. Anal. Chem. 75, 663–670 (2003).

